# 7q11.23 CNV alters protein synthesis and REST-mediated neuronal intrinsic excitability

**DOI:** 10.1101/2022.10.10.511483

**Authors:** Marija Mihailovich, Pierre-Luc Germain, Reinald Shyti, Davide Pozzi, Roberta Noberini, Yansheng Liu, Davide Aprile, Erika Tenderini, Flavia Troglio, Sebastiano Trattaro, Sonia Fabris, Ummi Ciptasari, Marco Tullio Rigoli, Nicolò Caporale, Giuseppe D’Agostino, Alessandro Vitriolo, Daniele Capocefalo, Adrianos Skaros, Agnese Franchini, Sara Ricciardi, Ida Biunno, Antonino Neri, Nael Nadif Kasri, Tiziana Bonaldi, Rudolf Aebersold, Michela Matteoli, Giuseppe Testa

## Abstract

Copy number variations (CNVs) at 7q11.23 cause Williams-Beuren (WBS) and 7q microduplication syndromes (7Dup), two neurodevelopmental disorders with shared and opposite cognitive-behavioral phenotypes. Using patient-derived and isogenic neurons, we integrated transcriptomics, translatomics and proteomics to elucidate the molecular underpinnings of this dosage effect. We found that 7q11.23 CNVs cause opposite alterations in neuronal differentiation and excitability. Genes related to neuronal transmission chiefly followed 7q11.23 dosage and appeared transcriptionally controlled, while translation and ribosomal protein genes followed the opposite trend and were post-transcriptionally buffered. Mechanistically, we uncovered REST regulon as a key mediator of observed phenotypes and rescued transcriptional and excitability alterations through REST inhibition. We identified downregulation of global protein synthesis, mGLUR5 and ERK-mTOR pathways activity in steady-state in both WBS and 7Dup, whereas BDNF stimulation rescued them specifically in 7Dup. Overall, we show that 7q11.23 CNVs alter protein synthesis and neuronal firing-established molecular and cellular phenotypes of neurodevelopmental disorders.

**Graphical abstract:** 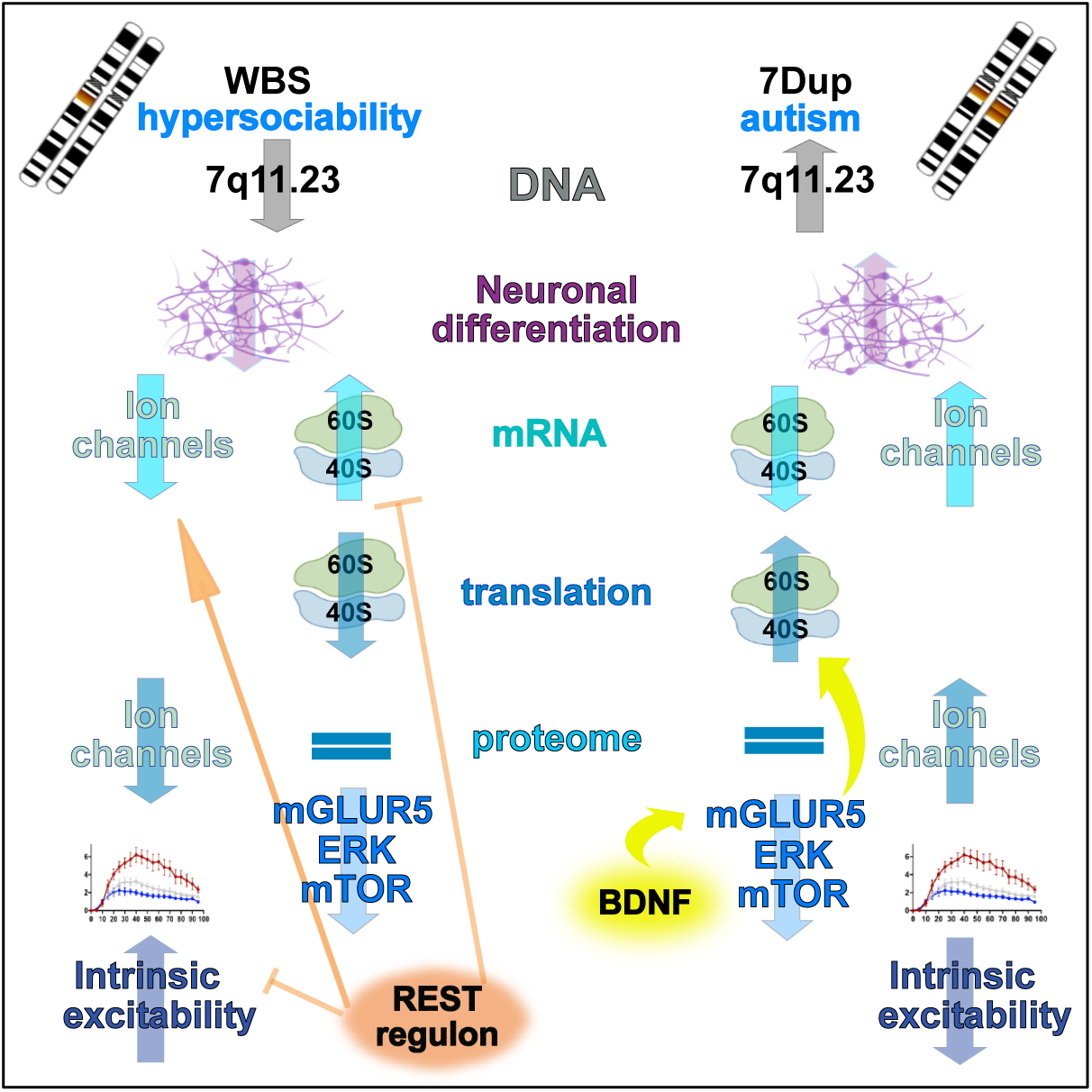

## Introduction

Neurodevelopmental disorders (NDDs), in particular autism spectrum disorder (ASD) and intellectual disability (ID), are among the most conspicuous instances of what Hyman called a ‘*terra incognita*’, referring to the “daunting polygenicity” of mental illness (Hyman, 2018). Over the past two decades, formidable advances in neurogenomics are increasingly breaking down long established neuropsychiatric diagnoses into a vast universe of, on the one hand, monogenic rare conditions and, on the other, of polygenic inheritance itself fragmented into a multiplicity of individual loadings of risk variants, with in between the expanding recognition of the combinatorial contribution of rare and polygenic variants in terms of phenotypic expressivity. With more than 1000 genes associated with ASD/ID (https://www.sfari.org/), including a number of multigenic copy number variations (CNVs) and a rising score of polygenic risk variants, understanding the molecular basis of their pathogenic convergence and defining the phenotypic scale at which it makes most sense to probe it mechanistically are two key challenges for the field. Against this backdrop, rare conditions, through their well-defined and amenable genetics, offer unique opportunities for exploring the reach of disease modeling approaches and test their potential for both mechanistic dissection and translational inroads. Among them, syndromes caused by symmetrically opposite CNV are particularly informative, since they offer the opportunity, by way of reciprocal cross-referencing, to precisely tease out phenotypic cascades in terms of their dosage-dependent components. Fueled by the power of large sample sizes, recent initiatives linking genome-wide regions of high dosage sensitivity with specific disorders are shedding light into CNVs’ properties and identifying thousands dosage sensitive genes, thereby deciphering the genetic causes of complex NDDs (Collins et al., 2022).

Within them, the 7q11.23 CNVs lead to a pair of multisystemic syndromes with shared and opposite neurocognitive and behavioral profiles. The hemideletion causes Williams-Beuren syndrome (WBS; OMIM (Online Mendelian Inheritance in Man) 194050), while its hemiduplication causes 7q microduplication syndrome (7Dup; OMIM 609757). Both NDDs feature mild to moderate intellectual disability, anxiety and facial dysmorphic features. However, while WBS is characterized by deficits in visuospatial construction, relative strength in language and hypersociability (Pober, 2010), 7Dup is associated with speech delay and ASD (Klein-Tasman and Mervis, 2018; Sanders et al., 2011; Velleman and Mervis, 2011). Despite some shared features, the strikingly opposite patterns in complex cognitive features of these two conditions suggest that the symmetry is maintained all the way, from the original CNV through the various layers of biological organization and regulation, up to behavior and cognition. The 7q11.23 locus comprises 26-28 genes including several key regulators of transcription and translation. Indeed, we previously dissected 7q11.23-related transcriptional dysregulation at the induced pluripotent stem cells (iPSC) stage, which was then selectively amplified upon the onset of neuronal differentiation (Adamo et al., 2015). We found that GTF2I, a key transcription factor (TF) of the 7q11.23 region, forms a repressive chromatin complex with KDM1A/RCOR2/HDAC2 and silences its dosage-sensitive genes.

Transcription and translation constitute key layers of dysfunction in NDDs (Ayhan and Konopka, 2019; Crino, 2016; Magdalon et al., 2017; Santos-Terra et al., 2021). Mutations in many TFs affect neurodevelopment by altering the activity of master regulators such as the RE-1 silencing transcription factor (REST), as shown for the high-confidence ASD/ID genes *CHD8* and *MECP2* (Ballas et al., 2005; Katayama et al., 2016). *REST* is a master regulator of neuronal differentiation and maturation. It represses neuronal genes in non-neuronal tissues and is a key factor in multistep repression complexes that control the orderly expression of genes during neuronal differentiation (Ballas et al., 2005; Chen et al., 1998). Likewise, deregulated protein synthesis and mutations in the mTOR (mechanistic target of rapamycin) pathway, a signaling hub that integrates internal and external signals to regulate cellular physiology and local protein synthesis, were initially linked to highly penetrant monogenic forms of ASD, and later on, proposed as a hallmark of the disorder (Kapur et al., 2017; Liu-Yesucevitz et al., 2011; Santini and Klann, 2014). The interplay between signaling pathways, local protein synthesis and proper synapse functioning was supported also by evidence for the key role of the excessive mGluR5 signaling in the cognitive phenotype of Fragile X syndrome, the most common mendelian form of ID/ASD (Dölen et al., 2007; Michalon et al., 2012; Osterweil et al., 2010). In addition, ribosome biogenesis has recently emerged as a possible underlying mechanism linking various NDDs (Hetman and Slomnicki, 2019). Therefore, a growing body of evidence supports the idea that mutations strongly associated with NDDs affect all layers of gene expression.

Although several teams, including our own, have already made significant contributions to understanding the molecular mechanisms underlying 7q11.23-related pathophysiologies (Adamo et al., 2015; Barak et al., 2019; Capossela et al., 2012; Chailangkarn et al., 2016; Deurloo et al., 2019; Khattak et al., 2015; Tebbenkamp et al., 2018; Zanella et al., 2019), the interplay between different layers of gene expression shaped by 7q11.23 CNV remains elusive. We hypothesized that the symmetry between 7q11.23 CNVs can act as a governing principle in deciphering clinically relevant pathways underlying sociability and language competence. By integrating transcriptomics, translatomics, proteomics, and electrophysiological analysis of glutamatergic neurons derived from 7q11.23 neurotypical and CNV-patients, along with a fully isogenic allelic series that recapitulates the dosages of the 7q11.23 locus, we were able to uncover the mechanisms that link 7q11.23 genetic dosage imbalances to key NDD phenotypes.

## Results

### Generation of an isogenic allelic series of 7q11.23 CNVs

Human genetic heterogeneity poses a formidable challenge for disease modeling, being at once the very aspect that one would wish to capture for patient-tailored precision but also a significant potential confounder for the dissection of gene-specific generalizable mechanisms of disease causation. Isogenic and patient-derived settings can thus provide complementary insights, allowing to focus, with the former, on the phenotypes exclusively imputable to the mutation at hand while excluding, with the latter, any potential artifact arising from the genetic manipulation *per se* or from the spurious interaction of the mutation with the chosen genetic background. Building on the empirical benchmarks we had previously derived from the meta-analysis of the two largest iPSC resources worldwide, we thus set out to complement our cohort of patient-derived iPSC lines (Adamo et al., 2015; Cavallo et al., 2020) with a fully isogenic allelic series that recapitulates, in the same genetic background, the three dosages of the 7q11.23 interval (hemiduplicated, control and hemideleted) CNVs (**Fig. 1A**). To this end, we exploited the presence of the 7q11.23 duplication in the cells originating from 7Dup patients and targeted the Cas9 onto the Williams-Beuren syndrome critical region (WBSCR). We used a single guide RNA (gRNA) that simultaneously recognized both duplicated sequences and consequently introduced an intrachromosomal deletion, thus effectively generating an isogenic healthy control in a female 7Dup background (isoCTL; **Fig. 1A**). Successively, we performed a second round of CRISPR/Cas9 editing starting from isoCTL to generate an isoWBS line, using two gRNAs that circumscribe the whole WBSCR (**Fig. 1A**). The deletions were screened by digital PCR assays on genomic DNA and validated by FISH analysis (**Fig. S1A** and **Fig. 1B**, respectively). Western blot analysis confirmed that isogenic iPSC lines preserved a 7q11.23 dosage imbalance of proteins encoded by genes located in the WBSCR interval (**Fig. S1B**), while a Short Tandem Repeat (STR) analysis confirmed the identity of the isogenic lines (**Fig. S1C**). Next, we generated neurogenin 2 (Ngn2)-driven induced cortical glutamatergic neurons (iNeurons; Zhang et al., 2013) by ectopic expression of Ngn2 delivered with a PiggyBac transposon system (Cavallo et al., 2020; Kim et al., 2016), which ensured high reproducibility between different rounds of differentiation (**Fig. S1D**). Isogenic iNeurons, faithfully recapitulated the dosage imbalances of WBSCR genes similar to those of patient-derived iNeurons, at the level of both transcriptome and proteome (**Fig. 1C-D; Fig. S1E**). Correlation analysis of transcriptome signatures from 30 days old patient-derived and isogenic iNeurons with the signature of the human developing neocortex (Mayer et al., 2019) revealed that our differentiation paradigm faithfully recapitulates cortical early upper layers neurons (GW: 16-18; **Fig. S1F**), in line with previous reports (Frega et al., 2017; Schörnig et al., 2021; Zhang et al., 2013). We further performed karyostat analysis to assess the genomic integrity of neurons derived from isogenic lines. The analysis uncovered a large amplification of the Chromosome 14 (Chr14) in isogenic lines, which probably originated in the mosaicism of the original patient line, used for generation of the isogenic lines (**Fig. S2A**; the list of amplified genes is provided in **Supplementary Table S1**). In line with the Chr14 amplification in all three genotypes, the comparison of the transcriptomes did not show any substantial change in the expression of the amplified region between the original 7Dup line and the isogenic derivatives (**Fig. S2B**). This unique series of isogenic lines thus provides the opportunity to study the effect of the CNVs in a highly controlled setting and in conjunction with patient-derived lines to reveal disease-relevant endophenotypes.

**Fig. 1.**
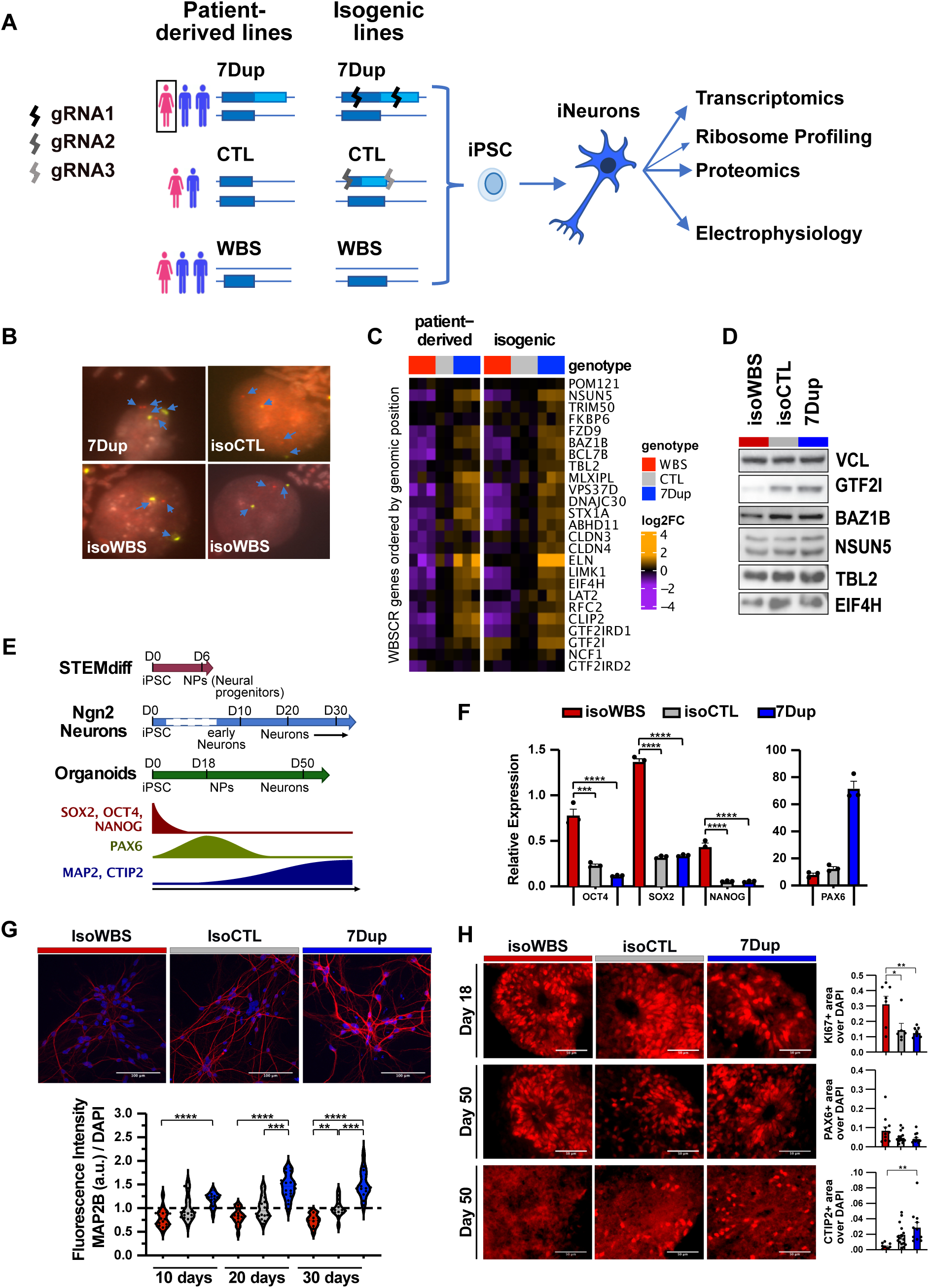
7q11.23 hemideletion delays, whereas hemiduplication accelerates neuronal differentiation. **(A)** Scheme of experimental design and generation of 7q11.23 isogenic lines (successive deletions leading from 7Dup to isoCTL and isoWBS; gRNAs - guide RNA; isoCTL - isogenic control; isoWBS - isogenic WBS line). **(B)** Two-color FISH analyses using 7 Alpha Satellite and ELN probes (green and red, respectively). ELN, which is located within WBS critical region (WBSCR), showed three signals in 7Dup, two in isoCTL, and one in isoWBS, corresponding to the WBSR copy number in respective clones; 7 Alpha Satellite was used as a control for the chromosomal number. Considering that the distance between two identical WBSCR on a duplicate chromosome is approximately the same as WBSCR length (∼ 1,8 Mb), the 7Dup signal pattern was resolved in interphase nuclei (metaphase chromosomes are thousands of times more compacted than interphase chromosomes). **(C)** WBSCR genes maintain the dosage at the level of RNA. RNAseq data for WBSCR genes are shown for both, patient-derived and isogenic neurons for all three genotypes. Although GTF2I transcripts were not downregulated in isoWBS, both the translatome and proteome data showed 7q11.23 dosage-dependent expression (Fig. S1E) that was confirmed also by western blot (Fig. 1D), suggesting that the upregulation observed at mRNA level is probably an artifact of sequencing or genomic engineering of non-coding and non-regulatory elements. **(D)** WBSCR genes maintain the dosage at the level of protein in induced glutamatergic Neurons. **(E)** Diagram showing the timing of neuronal differentiation in three different neuronal models: STEMdiff^TM^ differentiation, Ngn2-driven differentiation, and cortical brain organoids. The expected change in profiled neuronal markers during neuronal development is schematized. Arrows represent time points of analysis. **(F)** 7Dup, isoCTL and isoWBS were differentiated for five days using the dual Smad differentiation protocol (STEMdiff^TM^) and profiled for stem markers (OCT4, SOX2 and NANOG) and neural progenitor marker PAX6. To compare samples, we used one-way ANOVA followed by Tukey’s multiple comparison test. The significance level was set at p<0.05. *p<0.05; **p<0.01; ***p<0.001. **(G)** Representative immunofluorescence images of 30 days old Ngn2 iNeurons from isoWBS, isoCTL and 7Dup stained for the mature neuronal marker MAP2B (red) and DAPI (blue; upper panel). Graph showing the quantification of MAP2B fluorescence intensity over the cell number in Ngn2 iNeurons assessed at 10, 20 and 30 days of differentiation. IsoWBS and 7Dup were normalized over controls; data are means ± SEM from 2 independent differentiation protocols; 14-18 fields of view. The statistical comparisons were done with one-way ANOVA followed by Tukey’s multiple comparison test; **p<0.01; ***p<0.001 ****p<0.0001 (lower panel). **(H)** Immunofluorescence in cryosections of cortical organoids from isogenic lines on day 18 and 50 depict differences in the expression of the proliferative marker KI67, neural progenitor marker PAX6 and neuronal postmitotic marker CTIP2. First row: quantification of KI67, isoWBS n=3 organoids, isoCTL n=4, 7Dup n=3; second row: quantification of PAX6, isoWBS n=5 organoids, isoCTL n=4, 7Dup n=3; third row: quantification of CTIP2, isoWBS n=5 organoids, isoCTL n=4, 7Dup n=3. Data points are organoids’ sections. All the data are from two independent experiments. Data are shown as mean ± SEM. The statistical comparisons were done with one-way ANOVA followed by Tukey’s multiple comparison test. The significance level was set at p<0.05. *p<0.05; **p<0.01.

### 7q11.23 dosage alters neuronal differentiation in a symmetrically opposite manner

Upon differentiation into iNeurons, the occurrence of dosage-dependent differences in the dynamics of morphological changes (**Fig. S2C**) prompted us to systematically compare the kinetics of differentiation between genotypes starting from the earliest stages. We thus, exploited three different neuronal models (aligned timelines depicted in **Fig. 1E**), and assessed the expression of: *i*) pluripotency (OCT4, SOX2 and NANOG) and neural progenitor (PAX6) markers in early neural progenitor cells (NPCs) differentiated with STEMdiff™ (day 5; **Fig. 1F**), *ii*) mature neuron marker MAP2B in Ngn2-iNeurons (day 10, 20 and 30; **Fig. 1G**) and *iii*) proliferative marker KI67, PAX6 and postmitotic deep-layer neuron marker CTIP2 in day 18 and 50, respectively, in brain organoids (**Fig. 1H**). In all three neuronal models we found symmetrically opposite kinetics of differentiation, with isoWBS and 7Dup being, respectively, delayed and accelerated in comparison to controls. In particular, we found pluripotency markers still expressed in isoWBS while almost absent in isoCTL and 7Dup, whereas PAX6 followed an opposite trend, being higher in 7Dup and lower in isoWBS and isoCTL in NPCs (**Fig. 1F**). The expression of MAP2B followed 7q11.23 dosage, being higher in 7Dup and lower in isoWBs when compared to isoCTL at all three time-points assessed in Ngn2-driven iNeurons (**Fig. 1G**). Finally, KI67 was enriched in isoWBS compared to isoCTL and 7Dup brain organoids on day 18 (**Fig. 1H**). Conversely, on day 50-when brain organoids contain a mixed population of neural progenitors and postmitotic neurons (López-Tobón et al., 2019) we did not observe a significant difference in PAX6 expression, whereas the postmitotic neuronal marker CTIP2 was enriched in organoids derived from 7Dup line, confirming symmetrically opposite kinetics of differentiation (**Fig. 1H**). In agreement with increased KI67 expression, we observed a trend towards a larger size of isoWBS organoids compared with isoCTL and 7Dup (**Fig. S2D**). Together these results are in agreement with the longitudinal analysis of brain organoids and *GTF2I* dosage-specific murine models in the companion article by (Lopez-Tobon et al.), where we also found symmetrically opposite dynamics of neural progenitor proliferation and accelerated production of excitatory neurons in 7Dup. Therefore, these data robustly underscore that 7q11.23 gene dosage imbalances regulate the timing of neuronal differentiation both across models and patient-specific versus isogenic designs.

In addition to MAP2B (**Fig. 1G**), we investigated the expression of additional mature neuron markers by western blot analysis (**Fig. S3A**). While CAMK2A and CAMK2B showed the same trend as MAP2B, thus supporting further symmetrically opposite kinetics of differentiation, some other synaptic proteins such as SYP, NLGN1 and, in part, SYN1 showed an opposite trend (**Fig. S3A**), suggesting a more complex picture. Of note, SYN1 and NLGN1 are found among GTF2I targets (Kopp et al., 2020; McCleary-Wheeler et al., 2020). We next compared the iNeurons transcriptomes to a single-cell RNAseq time-course of Ngn2 differentiation (Lin et al., 2021), which revealed a slight differentiation delay of WBS lines (**Fig. S3B**), while no acceleration was observed for 7Dup. Sholl analysis at day 30 iNeurons showed no differences in branching and neurite length between genotypes (**Fig. S3C-D**), consistent with what we reported for the patient-derived lines in (Cavallo et al., 2020).

### Symmetrically opposite transcriptional regulation of translation and neuronal transmission genes in WBS and 7Dup

Isogenic and patient-derived iNeurons showed remarkably consistent 7q11.23-associated transcriptome alterations (**Fig. S4A**), though -as expected- not all alterations were significant in both, highlighting the value of using the two models as complementary ones. A merged analysis of the datasets revealed a highly reproducible signature of the CNVs (**Fig. 2A**) across over 2000 differentially-expressed genes (DEGs; FDR<0.01), which were largely, though not exclusively, dysregulated in symmetrically opposite directions in WBS and 7Dup. This revealed a largely linear dosage sensitivity, in line with the symmetrically opposite phenotypes of the syndromes in the neural domain. The 7q11.23-sensitive genes showed highly significant enrichments for a number of biological processes, with ribosomal genes and translation initiation factors being downregulated when 7q11.23 copy number dosage was increased (**Fig. 2B**, in red and **Fig. 2C**), whereas ion channels and synaptic transmission genes were upregulated (**Fig. 2B**, in green and **Fig. 2D**). Enrichment analysis for genotype-specific DEGs was less clearly related to neuronal function (**Fig. S4B**), further suggesting the relevance of symmetrically opposite alterations for the neurological phenotypes. In addition, the enrichments for cell cycle-related terms in WBS (**Fig. S4B**, upper panel) are in agreement with the observed differences in neuronal differentiation and increased proliferation of NPCs in isoWBS organoids (**Fig. 1E-H**; **Fig. S2D** and Lopez-Tobon et al.). Finally, the DEGs were significantly enriched for ASD-associated genes (Fisher p∼10^-12; the most significant ASD-associated DEGs are shown in **Fig. 2E**), underscoring the impact of 7q11.23 dosage on critical genes associated with social and cognitive phenotypes.

**Fig. 2.**
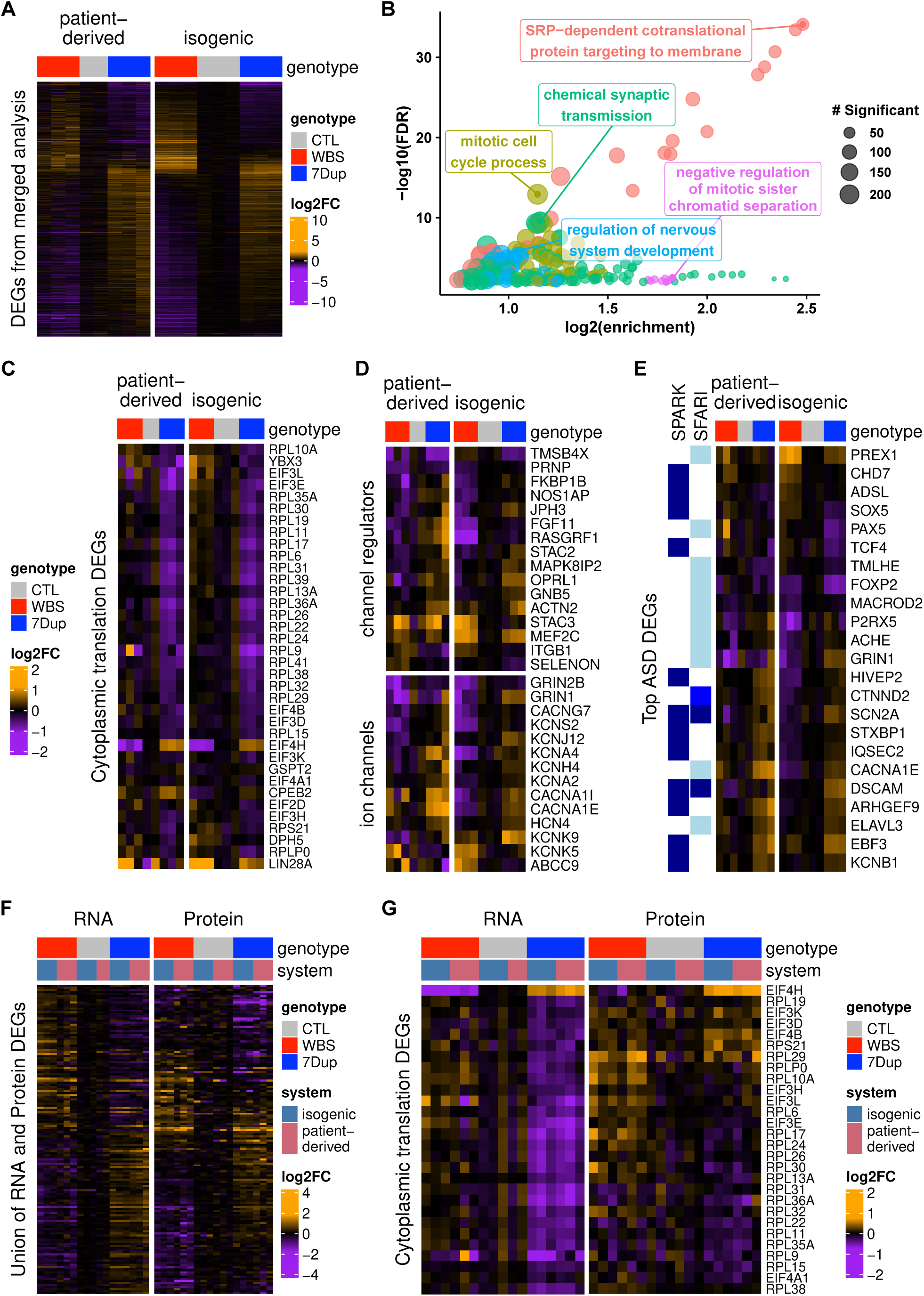
Robust transcriptional alterations in translation-and neural transmission-related genes. **(A)** Fold-changes of differentially-expressed genes (DEGs) in the merged analysis of isogenic and patient-derived lines (any condition), showing robust transcriptional signatures that are largely symmetrically opposite between genotypes. **(B)** Enriched Gene Ontology (GO) terms in the regression on 7q11.23 copy-number. Similar terms are clustered (denoted by colors) and only the top term per cluster is shown. **(C-E)** Top DEGs associated with translation (C), ion channels and their regulation (D) or ASD (E). **(F)** Fold-changes of the union of RNA and Protein DEGs (quantified in both assays), revealing largely consistent patterns across the two layers. **(G)** Expression of the transcriptionally-deregulated translation genes that could also be measured at the proteome level, highlighting a buffering in 7Dup.

### Neuronal transmission genes are transcriptionally controlled, while translation-related genes show dosage-dependent post-transcriptional regulation

As expected from the higher measurement noise and lower coverage of proteomics (2300 unique proteins quantified across all samples, and 3057 quantified across at least 75% of the samples), only a subset of differentially expressed genes could be detected as significantly altered also at the protein level. Nevertheless, for those DEGs that could be measured in both transcriptome and proteome, we observed that the proteome of both patient-derived and isogenic lines mostly recapitulated the transcriptome, albeit with partial buffering (**Fig. 2F**), i.e., mitigation of the impact of mRNA alterations on the proteome (Buccitelli and Selbach, 2020; Kusnadi et al., 2022). The distributions of fold-change differences between RNA and protein were consistent with an overall weak buffering, especially in genes downregulated in 7Dup or upregulated in WBS (**Fig. S4E**). Although proteins forming complexes often show stronger buffering, we observed only a weak effect in this direction. Of note, most of the buffering appeared condition-specific, like in the case of ribosomal proteins and translation initiation factors buffered specifically in 7Dup (**Fig. 2G**).

To investigate the post-transcriptional regulation underlying the observed buffering, we performed ribosome-protected fragments (RPF) analysis in isogenic iNeurons. Integrated analysis across the three layers (transcriptome, translatome and proteome) revealed significant buffering at the level of translation for a subset of genes and confirmed that these were different in WBS and 7Dup (**Fig. S4C-D**). To explore these different sets of genes, we clustered the union of differentially expressed genes with significant differences at both RNA and protein levels, according to their binarized fold change in each condition and layer (**Fig. 3A, Supplementary Table S2**). Genes whose expression by and large increased with copy number both at the RNA and protein levels (’forwarded opposite’, i.e., their symmetrically opposite dysregulation is forwarded from the transcriptome to the proteome) were related to protein polymerization, neuronal projections, synaptic plasticity and ion transport (**Fig. 3B**). In contrast, genes decreasing with copy number at the RNA level but which were mostly buffered in 7Dup at the protein level were related to translation, mostly ribosomal proteins and translation initiation factors (**Fig. 3C, Supplementary Table S2**). Other gene clusters did not show statistically significant enrichments. Hence while the expression of genes related to neuronal transmission is primarily transcriptionally controlled, translation-related genes display a more complex, multi-layered regulation that entails significant post-transcriptional buffering. Of note, many deregulated translation-related genes belong to the group of TOP mRNAs, which undergoes coordinated translation control through a common cis-regulatory element, the 5’ Terminal Oligo Pyrimidine motif (Meyuhas and Kahan, 2015). We thus probed the fold change distribution (isoWBS vs isoCTL and 7Dup vs isoCTL) at the level of RNA, RPF and proteins of the whole core set of genes with a 5’ TOP motif (Philippe et al., 2020). While 5’ TOP genes tended to be transcriptionally downregulated in 7Dup and upregulated in WBS, we observed a highly significant (KS p-value < 2e-16) opposite trend at the level of translation efficiency, which buffered their expression at the protein level, pointing to translation remodeling to counteract transcriptional imbalances (**Fig. 3D**).

**Fig. 3.**
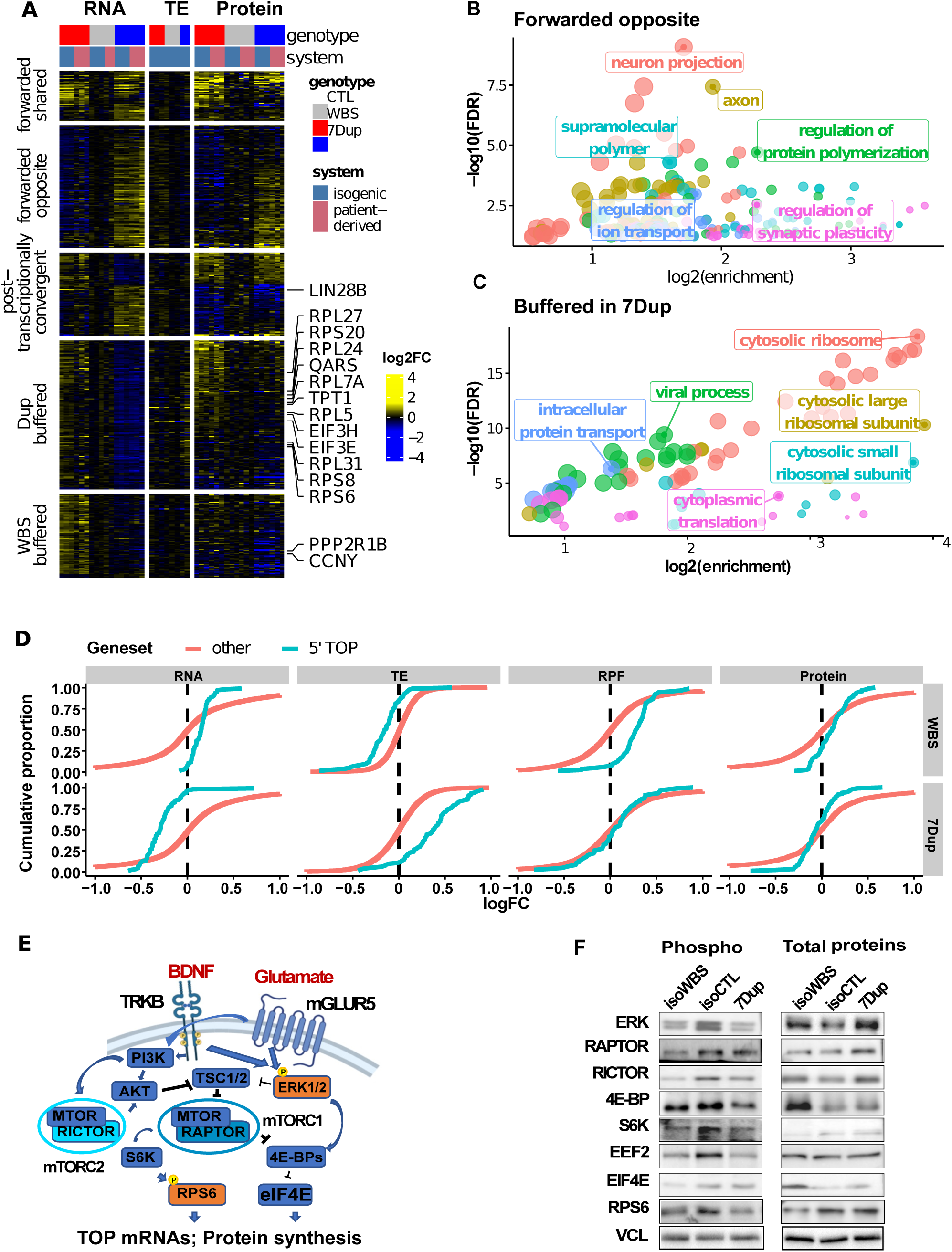
Translational buffering of TOP mRNAs in 7Dup. **(A)** Cross-layer clustering of DEGs reveals distinct patterns of regulation, emphasizing symmetrically-opposite DEGs forwarded to the proteome and 7Dup-specific buffering. The genes labeled are those found to have significant changes in translation efficiency. **(B-C)** Enriched Gene Ontology (GO) terms in the forwarded opposite (B) and buffered in 7Dup (C) clusters. **(D)** Translational buffering opposes the transcriptional dysregulation of genes encoding 5’ Terminal Oligo-Pyrimidine motif (TOP) containing mRNAs to maintain stable protein levels in 7Dup. Cumulative distribution plots comparing the fold-changes in both conditions and across gene expression layers of genes encoding 5’ TOP mRNAs to that of other genes are shown. Translation efficiency: TE; Ribosome protected fragments: RPF. **(E)** Scheme of ERK-mTOR signaling. **(F)** Shared downregulation of the mTOR pathway activity in WBS and 7Dup. Western blotting of total levels and phosphorylated, active, forms of major components of the mTOR pathway. Vinculin (VCL) was used as a loading control.

### Genotype-specific dysregulation of the mGLUR5-ERK-mTOR-protein synthesis axis

The mTOR and ERK signaling pathways (**Fig. 3E**) are the key regulators of the translation of TOP mRNAs (Meyuhas and Kahan, 2015). To assess the activity of both signaling pathways, we profiled the total protein levels and the phosphorylated (active) forms for their major components. While we did not observe any major change in the total protein levels, we found a consistent reduction of phosphorylated forms in both genotypes, indicating that this alteration in the activity of both signaling pathways is a shared feature of the 2 syndromes (**Fig. 3F**, see **Fig. S4F** for replicates). Given the fact that both pathways also regulate protein synthesis, we next examined the rate of global protein synthesis by using a puromycin incorporation assay (Aviner, 2020). We found a significant decrease in translation efficiency in isoWBS, while the effect was less pronounced in 7Dup iNeurons (**Fig. 4A**; replicates whose quantification is included in Fig 4A are shown in Fig. S4G). We observed no change in phospho-eIF2α (**Fig. S4F**), excluding its contribution to the observed downregulation in protein synthesis. Interestingly, we also found shared downregulation of the spontaneous excitatory postsynaptic current (sEPSC) frequency (**Fig. 4B**), which could be related to the observed downregulation of protein synthesis. The sEPSC amplitude was not changed (**Fig. 4B**).

**Fig. 4.**
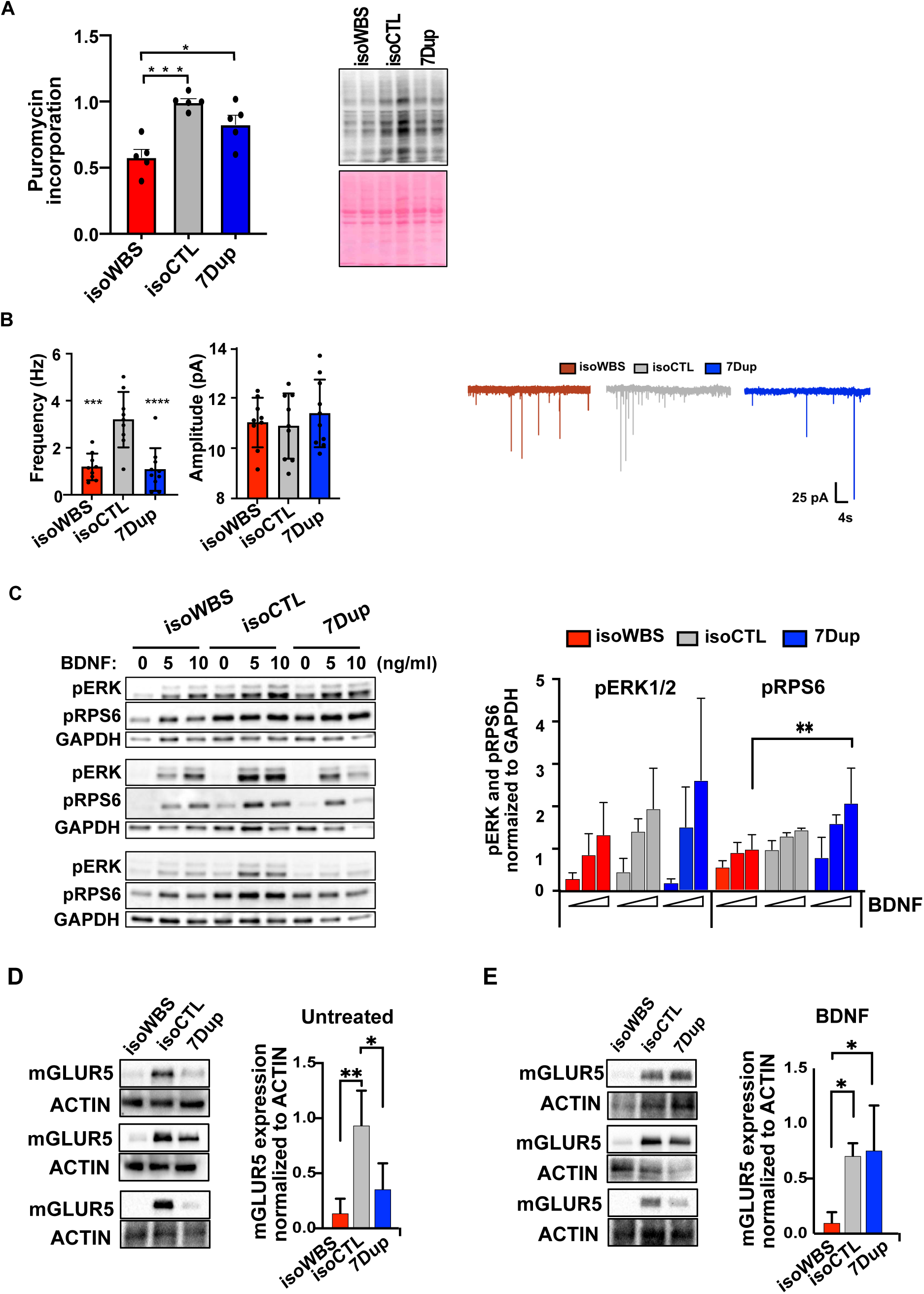
A short BDNF treatment rescues mGLUR5 expression and ERK-mTOR signaling. **(A)** Dosage imbalance of 7q11.23 triggers alterations in global protein synthesis. The quantification of five independent puromycin incorporation experiments is shown in a bar graph on the left. Puromycin incorporation assays were done in three independent rounds of neuronal differentiation. A representative western blot is shown on the right. Two-way ANOVA followed by Tukey’s multiple comparison test was used for the sample comparisons. The significance level was set at p<0.05. *p<0.05; **p<0.01; ***p<0.001. Additional replicates are shown in Fig. S4G. Red: isoWBS, gray: isoCTL, blue: 7Dup. **(B)** Symmetrical downregulation of spontaneous excitatory postsynaptic currents (sEPSCs) in isogenic iNeurons. Left, quantification of frequency and amplitude of sEPSCs. Right, representative whole-cell voltage-clamp recordings. Data represent mean ± SEM. ***p < 0.001, two-way ANOVA with post hoc Bonferroni correction. **(C)** A short BDNF treatment rescues ERK and mTOR signaling in 7Dup. Western blot profiling of pRPS6 (Serin 240-244) and pERK after 20 min treatment with increasing concentrations of BDNF is shown for three independents rounds of differentiation (left panel); pRPS6 (Serin 240-244) and pERK were used as a readout for the activity of mTOR and ERK pathways respectively. Quantification is on the right. GAPDH was used as a loading control. Two-way ANOVA followed by Tukey’s multiple comparison test was used for the sample comparison. The significance level was set at p<0.05. *p<0.05; **p<0.01; ***p<0.001. **(D)** mGLUR5 is downregulated in both WBS and 7Dup at steady-state. Western blot results from three independent rounds of differentiation (left panel) and corresponding quantifications (right panel) are shown. ACTIN was used as a loading control. Two-way ANOVA followed by Tukey’s multiple comparison test was used for the sample comparison. The significance level was set at p<0.05. *p<0.05; **p<0.01; ***p<0.001. **(E)** A short BDNF treatment rescues the mGLUR5 expression in 7Dup. Western blot results for three independent rounds of differentiation (left panel) and corresponding quantifications (right panel) are shown. ACTIN was used as a loading control. Two-way ANOVA followed by Tukey’s multiple comparison test was used for the sample comparison. The significance level was set at p<0.05. *p<0.05; **p<0.01; ***p<0.001.

In neurons, the mTOR and ERK pathways are activated by a brain-derived neurotrophic factor (BDNF) and glutamate through the stimulation of, respectively, the tyrosine receptor kinase B (TRKB) and metabotropic glutamate receptors 1 and 5 (mGLUR1 and 5; Lipton and Sahin, 2014). We thus probed how the decrease in the baseline activity of both pathways common to the two syndromes responded to activation by profiling phospho-RPS6 (pRPS6) at Serine 240/244 and phospho-ERK (pERK) upon stimulation with BDNF. At steady-state, we confirmed that pERK and pRPS6 were downregulated in both genotypes. However, upon stimulation with increasing concentrations of BDNF, we observed a 7Dup-specific rescue in both pERK and pRPS6 (**Fig. 4C**), which suggested additional, genotype-specific, dysregulation of upstream components of both signaling pathways. We, therefore, profiled TRKB and mGLUR5 expression in untreated and BDNF-stimulated conditions. In line with the diminished activity of the mTOR and ERK signaling pathways at steady-state, we found a significant downregulation of mGLUR5 in untreated samples of both isoWBS and 7Dup (along with the same trend for TRKB, **Fig. 4D** and **Fig. S4H**, respectively). However, only in 7Dup iNeurons, a short (20 min) BDNF treatment completely rescued mGLUR5 expression (**Fig. 4E**). BDNF treatment did not lead to a significant change in the TRKB expression (**Fig. S4H**). In conclusion, the mechanistic dissection of the translation control highlighted the mGLUR5-mTOR-ERK-protein synthesis axis centered on mGLUR5, thus further supporting the role of mGLUR5 in the cognitive-behavioral phenotypes of NDDs.

### Symmetrically opposite intrinsic excitability in WBS and 7Dup iNeurons

Next, in order to examine neuronal firing- a defining attribute of excitable cells and a key determinant of neuronal function-we performed whole-cell current-clamp recordings to quantify iNeurons’ intrinsic excitability. Specifically, we assessed the number of action potentials (AP) elicited by a series of incremental steps of current injection in iNeurons plated at low density (**Fig. 5A**). Induced neurons from WBS and 7Dup patients elicited a consistently higher and lower number of APs respectively, compared with iNeurons from CTL, across all current steps above 35pA (**Fig. 5B-C**). Conversely, passive properties (input resistance and resting potential) were comparable between genotypes, confirming the healthy state of the recorded neurons. In line with intrinsic excitability differences, WBS iNeurons exhibited a higher AP amplitude and a lower Rheobase compared to those from 7Dup (**Fig. 5D-G**). To gain more insights into the altered intrinsic excitability of iNeurons, we recorded Na^+^ and K^+^ currents (**Fig. S5**). Na^+^ and K^+^ currents were comparable in neurons from all the 3 genotypes. To confirm that the observed effect in intrinsic excitability was the result of gene dosage differences at the 7q11.23 locus and not due to cell line variability, we repeated the experiments with isogenic iNeurons (**Fig. 5H**) and confirmed that isoWBS iNeurons generated a higher number of APs compared with iNeurons from either isoCTL or 7Dup across all current steps above 15pA (**Fig. 5I-J**). Similarly, AP amplitude was also consistently higher in iNeurons from isoWBS compared to those from isoCTL and 7Dup, while passive properties were unaltered (**Fig. 5K-N**). These results uncover a 7q11.23 CNV-dependent selective impact on neuronal excitability that is highly robust across patient-derived and isogenic settings.

**Fig. 5.**
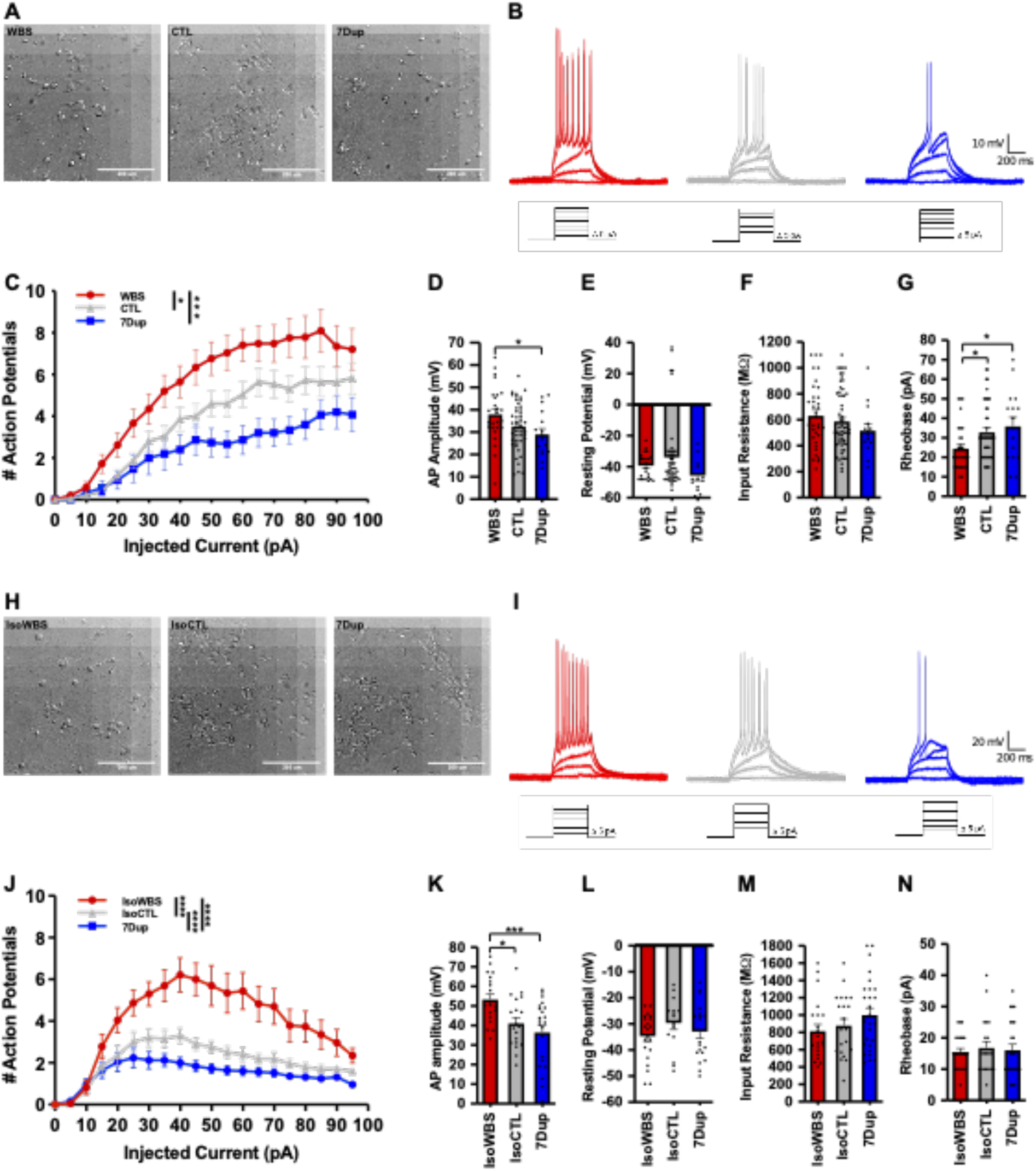
7q11.23 CNVs cause symmetrically opposite intrinsic neuronal excitability dynamics. **(A)** Brightfield images of CTL, WBS and 7Dup patient-derived iNeurons **(B)** Representative action potential (AP) trains in response to steps of 5 pA depolarizing current lasting 500 ms from −60 mV in WBS, CTL and 7Dup patient-derived iNeurons recordings, respectively. **(C-D)** Quantitative analysis depicting the number (C) and bar graph depicting the amplitude (D) of elicited APs in the current-clamp configuration in the 3 genotypes of patient-specific iNeurons (WBS: 4 cell lines n=29 neurons; CTL: 3 cell lines n=45 neurons; 7Dup: 4 cell lines n=16 neurons) in response to increasing current steps. **(E-G)** Input resistance was calculated in the current-clamp mode without any current injection. Resting potential was calculated in voltage-clamp mode using a pulse test of 10 mV. Rheobase was calculated as the minimum current required to elicit at least one AP. **(H)** Brightfield images of isogenic iNeurons **(I)** Representative AP trains in response to steps of 5 pA depolarizing current lasting 500 ms from −60 mV in isogenic iNeurons. **(J-K)** Quantitative analysis depicting the number (J) and bar graph depicting the amplitude (K) of elicited APs in the current-clamp configuration in the isogenic iNeurons (isoWBS: n=23 neurons, isoCTL: n=22 neurons, 7Dup: n=25 neurons). **(L-N)** Passive properties and Rheobase of the iNeurons recordings from isogenic lines, calculated as above. All data in Fig. 5 are shown as mean ± SEM and is the average of 3 independent experiments. For comparing AP frequency, we used two-way ANOVA followed by Tukey’s multiple comparison test, while for comparing passive properties we used one-way ANOVA followed by Tukey’s multiple comparison test. The significance level was set at p<0.05. *p<0.05; **p<0.01; ***p<0.001.

### The REST regulon mediates dosage-dependent pathophysiological phenotypes

The CNV-dependent symmetrically opposite endophenotypes in transcription and intrinsic excitability prompted us to search for the mediating factors. We thus performed a master regulator analysis, estimating TFs activities based on their curated targets (see methods). This predicted a number of TFs as changing in activity linearly with 7q11.23 copy-number. Interestingly, several of them were also differentially expressed at the transcriptional level in a 7q11.23 copy-number dependent manner, pointing to an extensive transcriptional rewiring determined by 7q11.23 dosage (**Fig. 6A**, squared TFs). Among them, we prioritized REST for functional interrogation, given its well-established role as key regulator of neuronal differentiation by controlling the expression of neuron-specific genes including those for intrinsic excitability (Aoki et al., 2012; Ballas et al., 2005; Mucha et al., 2010; Pozzi et al., 2013). Despite its transcriptional upregulation in WBS and downregulation in 7Dup iNeurons (**Fig. S6A**), we found no change at the REST protein level between genotypes (**Fig. S6B**). Given its ranking in the master regulator analysis, we thus hypothesized that alterations in the composition of REST-containing transcriptional complexes could be responsible for the transcriptional rewiring we observed, and we thus analyzed the expression of the REST interactome (Lee et al., 2016). We found that nearly all reported REST interactors (Lee et al., 2016) were transcriptionally deregulated in a 7q11.23 dosage-dependent manner (**Fig. S6C**), including HDAC2, which we also previously showed to interact with GTF2I (Adamo et al., 2015). To functionally validate the involvement of REST, we treated isoWBS iNeurons, which show downregulation of ion channels and other REST targets, with the REST inhibitor X5050 (Charbord et al., 2013). Inspection of the genes rescued by the REST inhibition revealed a set of potassium channels, including those that changed with 7q11.23 copy-number (**Fig. 6B** and **6C**, respectively) and known to be critical regulators of membrane potential and intrinsic excitability (Frankenhaeuser and Hodgkin, 1956). Interestingly, the treatment also triggered the downregulation of several important translation initiation factors and ribosomal protein transcripts that we found downregulated in 7Dup, thereby corroborating a REST-dependent recapitulation of their dosage-dependency (**Fig. 6D**). Finally, we confirmed that administration of X5050 rescued WBS iNeurons intrinsic excitability, restoring a physiological firing rate and AP amplitude comparable to control iNeurons (**Fig. 6E-F**). The passive properties of iNeurons were not altered by the treatment, indicating that the rescue of the excitability was not attributable to modulation of neither input resistance nor membrane potential (**Fig. 6 G-I**). Together, these findings uncover dosage-dependent REST dysregulation as a convergent mechanism underlying the transcriptional, translational and electrophysiological endophenotypes of 7q11.23 CNV and pave the way for the preclinical validation of REST inhibition in WBS.

**Fig. 6.**
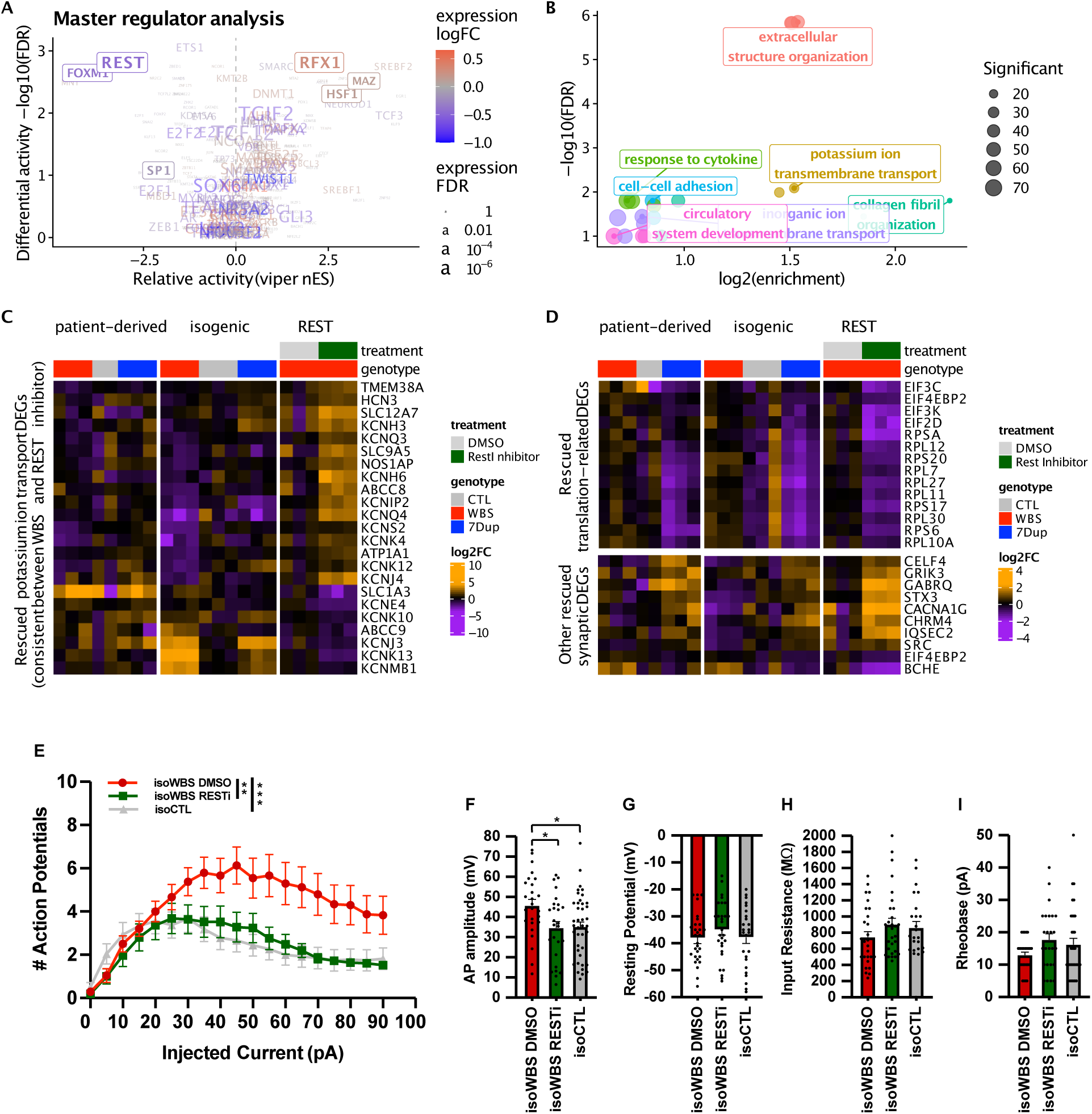
REST mediates WBS pathophysiological phenotypes. **(A)** Master regulator analysis on the 7q11.23 dosage-dependent genes based on transcription factor curated targets. The x and y axes respectively indicate the magnitude and significance of the inferred changes in the activity, while the color and size respectively indicate the magnitude and significance of the change in expression of the factor at the RNA level. Factors in a box are consistent and statistically significant in both activity and expression. **(B)** The genes altered in WBS versus control and rescued by REST inhibition in isoWBS iNeurons are associated with potassium ion transmembrane transport, extracellular structural organization and cation transport. **(C)** Heatmap showing potassium channel genes that were consistently differentially expressed in WBS iNeurons and were rescued by REST inhibition. **(D)** Heatmap showing WBS DEGs related to protein translation and synaptic communication that were rescued by REST inhibition. **(E)** REST inhibition rescues AP frequency in isoWBS iNeurons. Quantitative analysis depicting the number of AP elicited in isoWBS iNeurons treated with REST inhibitor or DMSO controls. Note how REST inhibition normalizes AP frequency to isoCTL iNeurons levels (isoWBS-DMSO, n=24 neurons; isoWBS-RESTi, n=25; isoCTL, n=27). **(F-I)** Passive properties were not affected by the REST inhibitor, AP amplitude instead was lowered. Data are shown as mean ± SEM. All the data is the average of 3 independent experiments. For comparing AP frequency, we used two-way ANOVA followed by Tukey’s multiple comparison test, while for comparing passive properties we used one-way ANOVA followed by Tukey’s multiple comparison test. Significance level was set at p<0.05. *p<0.05; **p<0.01; ***p<0.001.

## Discussion

Reprogramming-based disease modelling designs (Hyman, 2018) that compare cell lines derived from patients and healthy individuals suffer from lower sensitivity and are inherently prone to the confounding effects of spurious endophenotypes arising from differences in individual genetic backgrounds rather than authentic pathogenic mechanisms (Germain and Testa, 2017). Moreover, while patient-specific approaches are a cornerstone of precision disease modelling since they afford the unique opportunity to match the specificity of clinical histories to that of molecular phenotypes, such case-control designs greatly benefit from complementary, isogenic approaches that offer intrinsically a more direct route to establish causality between genetic lesions and endophenotypes. In this study, we integrated patient-derived with isogenic neuronal models engineered to harbor the full 7q11.23 CNVs. This approach enabled the identification of robust molecular, cellular and electrophysiological endophenotypes caused by 7q11.23 genetic dosage imbalances, including symmetrically opposite dynamics of neuronal differentiation, transcriptional alterations and differences in intrinsic excitability of iNeurons, as well as complex and CNV-specific patterns of post-transcriptional dysregulation. In turn, especially for human genetic variations that cause complex cognitive/behavioral phenotypes, this systematic engagement of multiple phenotypic layers (transcriptome, translatome, proteome, differentiation dynamics and excitability) addresses a major question of the disease modelling field, namely how, in actual disease-relevant cell types, phenotyping at the level of the transcriptome (arguably the most proximal and tractable layer) reverberates through more distal endophenotypes. In tracking symmetry from genotypes across phenotypes, we thereby offer the regulatory complexity that emerges from this high-resolution example as a template also for further efforts aimed at dissecting actionable disease mechanisms that specifically emerge from such multi-layered endophenotyping.

We identified largely symmetrically opposite transcriptional alterations in iNeurons, only part of which are buffered at the level of the proteome by a remodeling of translation regulation. The transcriptional changes that were not buffered were enriched for genes related to neuronal transmission, in particular synaptic genes and ion channels, which is consistent with the symmetrically opposite pattern of intrinsic excitability that we observed. Master regulator analysis revealed the REST regulon as key target of 7q11.23 dosage, whose activity was modulated accordingly. Its inhibition rescued the electrophysiological and corresponding transcriptional alterations in WBS, consistent with the crucial role that REST plays in regulating neuronal-specific genes coding for ion channels (McClelland et al., 2011, 2014; Pozzi et al., 2013), which in turn determine the electrical properties of neurons and drive intrinsic excitability. We showed that REST inhibition rescued the expression of several potassium channel genes, notably *KCNQ3*, a key mediator of M-currents that stabilizes membrane potential and suppresses neuronal excitability (Mucha et al., 2010; Wang et al., 1998). Hence, our findings led us to posit a model in which increased REST activity in WBS neurons suppresses the expression of potassium channels, including *KCNQ3*, thereby leading to the abnormally high intrinsic excitability (and the opposite in 7Dup), while REST inhibition rescues potassium channels expression and consequently lowers intrinsic excitability.

The expression of ribosomal proteins decreases during neuronal differentiation (Harnett et al., 2021; Slomnicki et al., 2016). However, the molecular mechanism of how this coordinated regulation occurs remains elusive. Here we show that REST inhibition reduced the expression of several ribosomal protein genes in WBS iNeurons, which is in contrast to the previously reported activation of ribosome biogenesis by knocking out *Rest* in quiescent neural progenitors in adult mice (Mukherjee et al., 2016). A possible explanation for the opposite effect of REST on ribosome biogenesis at different stages of neuronal development may be in its interaction with other TFs to regulate ribosomal biogenesis. For example, the binding sites for transcriptional activator SP1 have been found in most human ribosomal protein promoters (Petibon et al., 2021) and it was shown that SP1 regulates the transcription of ribosomal RNA and some ribosomal proteins (Nosrati et al., 2014; Rajput et al., 2016). In addition, it was also shown that REST fine-tunes SP1 levels to coordinate the expression of common target genes (Plaisance et al., 2005). Thus, given the fact that SP1 also featured prominently in our master regulator analysis, we speculate that fine-tuning of REST-SP1 interaction may regulate ribosome expression during neurodevelopment.

Not all the 7q11.23 dosage-dependent transcriptomic changes were reflected in the proteome, and most of the buffering was genotype-specific. The expression of translation-related genes, in particular the ribosomal genes with 5’ TOP motif, which was anti-correlated with 7q11.23 dosage at the transcriptional level, was translationally buffered in 7Dup. TOP mRNAs translation is regulated by the mTOR pathway (Meyuhas and Kahan, 2015) and mTOR stimulation partially rescued NDDs that are characterized by the deregulation of ribosomal biogenesis (Hetman and Slomnicki, 2019; Ricciardi et al., 2011; Xu et al., 2015, 2016). Therefore, we originally hypothesized that high mTOR activity drives increased translation of TOP mRNAs in 7Dup. Instead, we found a decrease in basal activity of both ERK and mTOR pathways as well as downregulation of mGLUR5 expression and global protein synthesis in both genotypes. Surprisingly, a 20 min treatment with BDNF led to a complete rescue of mGLUR5 expression and activity of both signaling pathways specifically in 7Dup, revealing a novel BDNF-mGLUR5-ERK-mTOR axis that operates in ASD-associated 7q11.23-dependent phenotype. The relevance of this axis can now be further explored also in other mGLUR5-associated ASD phenotypes.

Interestingly, pharmacological intervention on mGluR5 activity has been shown to modulate REST expression and signaling to the nucleus (de Souza et al., 2020) and, importantly, REST inhibition in WBS rescued the expression of key translation genes, suggesting that the transcriptional and translational aspects are heavily intertwined. *REST* expression decreases during neuronal differentiation both in development (Ballas et al., 2005; Kuwabara et al., 2004; Mukherjee et al., 2016) and in Ngn2-driven differentiation (**Fig. S6D**), along with the decreased expression of ribosomal proteins (Harnett et al., 2021; Slomnicki et al., 2016), while synaptic and ion channel genes instead increase (Fig. S6D). It is therefore attractive to relate all these changes to the delay and acceleration in differentiation, respectively observed in WBS and 7Dup, both in the present and in the companion article (Lopez-Tobon et al., same issue). Indeed, an increasing body of evidence from us and others suggests that changes in the dynamics of neuronal differentiation (acceleration and delay) are a major point of convergence across NDDs, despite differences in the underlying molecular mechanisms (Lalli et al., 2020; Paulsen et al., 2022; Villa et al., 2022).

In conclusion, starting from the pair of 7q11.23 CNV featuring a paradigmatic suite of symmetrically opposite and shared manifestations, we uncover several dosage-dependent endophenotypes in domains of dysfunction that are well-established proxies of cognitive-behavioral phenotypes, revealing a multilayered interplay between kinetics of differentiation, transcriptional and translational control, and neuronal function, which can productively inform the study also of other NDDs.

## Materials and Methods

### Human samples

In this study we used patient-derived iPSC lines that we have previously generated and already reported in (Adamo et al., 2015; Cavallo et al., 2020). See also Table 3.

Participation of the patients and healthy control individuals along with skin biopsy donations and informed consent procedures were approved by the ethics committees of the Genomic and Genetic Disorder Biobank (Casa Sollievo della Sofferenza, San Giovanni Rotondo, Italy) and the University of Perugia (Azienda Ospedaliera–Universitaria ‘Santa Maria della Misericordia’, Perugia, Italy).

Briefly, the biological samples from which we derived the iPSC lines used in this study are archived in the Genetic and Genomic Disorders Biobank (GDBank), which is part of the Telethon Network of Genetic Biobanks (TNGB; https://www.telethon.it/en/scientists/biobanks). The primary fibroblasts samples were collected from the following sources:

The Genetic and Genomic Disorders Biobank (GDBank) (Dr. Giuseppe Merla, Casa Sollievo della Sofferenza, San Giovanni Rotondo (Italy) for samples Ctl01C (Female - F), WBS01C (Male -M), WBS02C (F), Dup02K (F), Dup03B (M). Dr. Paolo Prontera, University of Perugia (Azienda Ospedaliera–Universitaria “Santa Maria della Misericordia”, Perugia, Italy) for line Dup01G (M). Dr. Frank Kooy, University of Antwerp, Antwerp (Belgium) for line Dup04A (M). The Wellcome Trust Sanger Institute, Cambridge (UK) provided upon purchase the iPSC line Ctl08A (M).

SNPs reports are available in (Adamo et al., 2015; Cavallo et al., 2020).

### iPSC culture

iPSC lines were cultured on plates coated with human-qualified Matrigel (BD Biosciences; dilution 1:40 in DMEM-F12) in mTeSR (StemCell Technologies) or TeSR-E8 medium (Stemcell Technologies) supplemented with penicillin 100 U/ml and streptomycin (100 μg/ml). Cells were passaged with ReLeSR (StemCell Technologies) or Accutase (Sigma). For single-cell culture, cells were plated in mTeSR or TeSR-E8 medium supplemented with 5 μM Y-27632 (Rock inhibitors (RI), Sigma).

### Generation of isoCTL and isoWBS

The Isogenic lines, isoCTL, and isoWBS, have been generated in two consecutive rounds of CRISPR/Cas9 genome editing by co-transfecting purified Cas9 protein and specific gRNAs. For the generation of isoCTL, we exploited the duplication of WBSCR in 7Dup patient (242K), introducing only one guide RNA (gRNA) that cut both duplicated WBSCRs, and we screened for the combination where the 5’ end of the first WBSCR fused to the 3’ end of the second WBSCR, thus generating one complete WBSCR on the place of initial duplication (Fig.1A). For the generation of the isoWBS instead, we used two gRNAs, centromeric and telomeric, which delineate the whole WBSCR. We designed forward primers for gRNAs containing universal forward primer for T7 promoter (in blue), thus allowing *in vitro* transcription. *In vitro* transcription was performed on a purified PCR template for gRNA. The primers for isoCTL were F1 and R1, while for isoWBS, we designed F_CE and R_CE for centromeric cut and F_TE and R_TE for a telomeric cut. The gRNAs were designed by using the MIT CRISPR design tool.

Primers for gRNAs (blue T7promoter sequence; black: gRNA sequence)

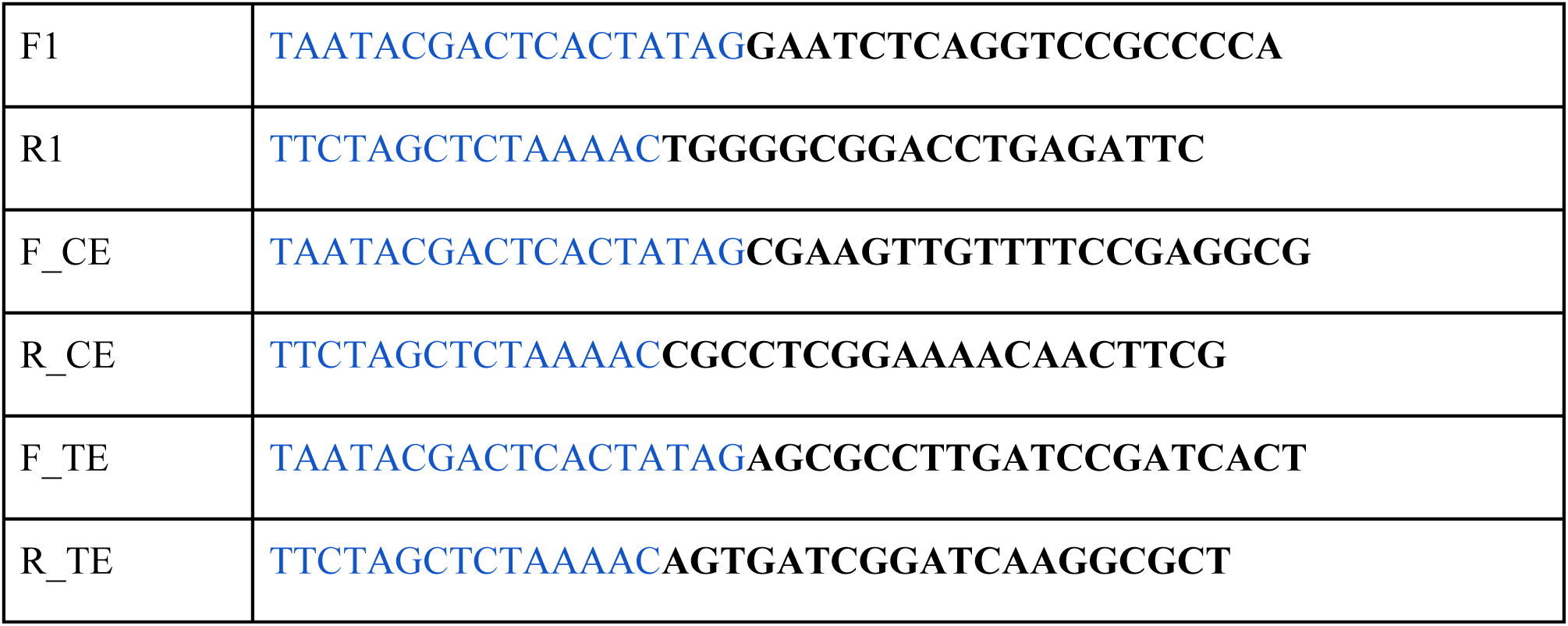

PCR reaction (5 μl Phusion HF buffer 5x, 0.25 μl Phusion Polymerase, 0.5 μl 10 mM dNTPs, 1 μl Tracer Fragment + T7 primer Mix, 1 μl F/R oligonucleotide mix (0.3 μM) and 17,25 μl H2O) was done as follows: initial denaturation 98°C, 10 sec; 32 cycles of denaturation 98 °C, 5 sec and annealing 55°C, 15 sec, followed by final extension 72°C, 60 sec. *In vitro* transcription (8 μl of NTP mix (100 mM ATP, GTP, CTP, UTP), 6 μl PCR template of gRNA, 4 μl TranscripAid Reaction Buffer 5x, 2 v TranscriptAid Enzyme Mix) was performed for 2 hours at 37 °C, followed by DNAase treatment (1 μl of DNase I, 1 U/μl) for 15 min at 37 °C. The product was visualized on agarose gel and purified with 5M ammonium acetate. Briefly, add ½ Volume (V) of 5 M ammonium acetate, 3V 100% EtOH, precipitate 1h at −80 °C, and spin 30 min at 13000 rpm, +4°C. The RNA pellet was dried for 25 min and resuspended in 200 μl RNase-free H2O.

We co-transfected 4×10^5 cells (seeded in a well of MW6) with 20 μg Cas9 and 10 μg gRNA. Cas9 was resuspended in Cas9 Transduction buffer 5x (500 mM NaCl, 25 mM NaH2PO4, 250 mM NDSB-201, 150 mM glycerol, 75 mM glycine, 1.25 mM MgCl2 and 1 mM 2-mercaptoethanol, pH8.0) to obtain final concentration of 3.88 μg/μl. The Cas9/gRNA mix was incubated at 37°C for a maximum of 10 min and then electroporated. The electroporation was performed using the Neon Transfection System (MPK10096, Thermo Fisher Scientific). The iPSC were pretreated for 2h with 5 μM RI, detached with accutase to obtained single-cell suspension, and resuspended in buffer R supplemented with Cas9/gRNA mix for a total volume of 120 μl for 4×10^5 cells. The electroporation was performed at 1300 volts for 20 ms and 2 pulses. The cells were seeded in mTeSR supplemented with RI. Electroporated cells were single-cell sorted in 96 well plates by using DAPI, 24h after electroporation. The medium was not changed for the first 5 days, and from the 6th day on, only ½ of the medium was changed every second day. We started to change the whole medium only when the colonies started to be visible. When confluent, cells were split first into 48 well plates, then 12 well plates, and screened when they reached confluency in 6 well plates. We obtained 17 clones for isoCTL and 8 for isoWBS, which were further analyzed by digital PCR and FISH for copy number variation of WBSCR.

### Fluorescence *in situ* hybridization (FISH) analysis

A co-hybridization using a specific clone covering the ELN locus at 7q11.23 and a specific alpha-satellite as a control probe for ploidy status was performed to investigate the ELN gene pattern iPSCs. Specifically, CTD-2510G2 BAC clone for the ELN gene (red signal) and 7 Alpha Satellite probes (green signal) were used for the identification of the gene copy number. The BAC clone was selected using the University of California Santa Cruz Genome Browser Database (http://genome.ucsc.edu/) and was tested on normal human metaphase cells to verify the absence of cross-hybridization while the alpha-satellite probes were kindly provided by Dr. M. Rocchi, University of Bari, Italy. FISH experiments were performed as previously described, with minor modifications (Fabris et al., 2005).

### Digital PCR

Eighty ng of DNA was amplified in a reaction volume of 15 μl containing the following reagents: 7.5 μl of “QuantStudio™ 3D Digital PCR Master Mix v2, Thermofisher”, 0.75 μl of “TaqMan Copy Number Assays, SM 20x, Thermofisher” FAM labeled and 0.75 μl of “TaqMan Copy Number Reference Assay 20x, Thermofisher” VIC labeled (for details see below). The mix was loaded on a chip using the QuantStudio 3D Digital PCR Chip Loader. The chips were then loaded on the Proflex PCR System (Thermofisher), and the PCR was carried out using a pre-PCR step of 10 min at 96° C, followed by 39 cycles of 2 min at 60 °C and 30s at 98 °C, followed by 2 min at 60 °C.

Data were analyzed with “QuantStudio 3D Analysis Suite Cloud Software”. The entire process was performed by the qPCR-Service at Cogentech-Milan, Italy. TaqMan Copy Number Assays (FAM): CLIP2 Hs03650293_cn, BAZ1B Hs04952131_cn andEIF4H Hs01632079_cn; TaqMan Copy Number Reference Assay: TERT (VIC) catalog number 4403316.

### Short Tandem Repeat (STR) analysis

STR analysis was done with GenePrint 10 System (Promega) according to the manufacturer’s instructions. STR loci consist of short (3-7 nt), repetitive sequence elements. They are well distributed with the human genome and represent a rich source of highly polymorphic markers which can be detected by PCR. GenePrint 10 System allows co-amplification and 3-color detection of nine human loci, providing a genetic profile with a random match probability of 1 in 2.92 x 10^9. PCR master mix contained 5x master mix, 5x primer master mix, and 2 ng of genomic DNA. The thermal cycling protocol was one recommended by the manufacturer (96 °C −1 min; 30 cycles of 94 °C −10 sec, 59 °C −1 min, 72 °C −30 sec; 60 °C −10 min; 4 °C).

### Karyostat

The KaryoStat™ (Thermo Fisher) assay allows for digital visualization of chromosome aberrations with a resolution similar to g-banding karyotyping by relying on 150k SNP probes across the human genome. The size of structural aberration that can be detected is > 2 Mb for chromosomal gains and > 1 Mb for chromosomal losses. Genomic DNA was extracted from pellets corresponding to 2 x 10^6 cell by using Genomic DNA Purification Kit (Catalog #: K0512) and quantified using the Qubit™ dsDNA BR Assay Kit (Catalog #: Q32850). 250 ng of total gDNA was used to prepare the GeneChip® for KaryoStat™ according to the manual, and the array was used to look for SNPs, copy number variants and single nucleotide polymorphisms across the genome.

### PiggyBac transposon system

For robust and rapid glutamatergic neuron differentiation, we used PiggyBac transposon system for the Neurogenin-2 (NgNn) delivery to the cells, as previously described (Cavallo et al., 2020). Briefly, mouse Ngn2 cDNA, under tetracycline-inducible promoter (tetO), was transfected into iPSCs by enhanced PiggyBac (ePB) transposon system (Kim et al., 2016). For each iPSC line, 4×10^5 cells were electroporated with 2,25 μg of the ePB construct carrying the inducible Ngn2 cassette and 250 ng of the plasmid encoding transposase for the genomic integration of the inducible cassette. Electroporations were performed using the Neon Transfection System (MPK10096, Thermo Fisher Scientific). iPSCs were selected using blasticidin 5 μg/ml (R21001, Gibco) for 5 days, and stable iPSC lines were stocked.

### Ngn2 neuronal differentiation

In order to obtain cortical glutamatergic neurons (iNeurons), iPSCs were dissociated with Accutase (GIBCO, Thermo Fisher Scientific) and plated on Matrigel-coated plates (final 2% v/v, Corning) in mTeSR™ or TeSR-E8 (Stemcell Technologies) supplemented with 5 μM Y-27632 (Sigma). iPSCs were then cultured in MEM1 (DMEM/F12 1:1 (Euroclone/Gibco) supplemented with NEAA 1%, N2 1%, BDNF 10 ng/ml, NT-3 10 ng/ml, Laminin 0,2 μg/ml and 2 μg/ml doxycycline hydrochloride, penicillin 100 U/ml and streptomycin 100 μg/ml) for 2 days. On the second day of MEM1, cells were selected with 1 μg/ml puromycin, to reassure that only the cells with Ngn2-inducible cassette will survive. For patient-derived iNeurons, after two days of MEM1, media was change to Neurobasal medium (NB, Thermo Fisher Scientific) supplemented with BDNF 10 ng/ml, NT-3 10 ng/ml, B27 (1:50), GlutaMax (Gibco, 1:100), penicillin 100 U/ml, streptomycin (100 μg/ml), which was previously conditioned on mouse astrocytes for 24 h. Half of media was changed every other day. Instead, for isogenic iNeurons after two days of MEM1, media was change to Neurobasal Plus medium (NB-Plus) composed of B-27™ Plus Neuronal Culture System (Thermo Fisher Scientific) supplemented with Glutamax 0.25% (Thermo Fisher Scientific), 2 μg/ml, doxycycline hydrochloride and penicillin 100 U/ml, streptomycin (100 μg/ml). The media was changed twice a week. On day 7-8, cells that already acquired neuron-like shape were dissociated with Accutase, counted and seeded into plates coated with 15 μg/ml of poly-D-lysine at a density of 1 x 10^6 cells/well of multiwell 6, 2 x10^6 in 6 cm dishes or 4 x10^6 in 10 cm dishes (Nunc Edge plates, Thermo Fisher Scientific) in conditioned NB or NB-Plus. Half of media was changed twice a week until day 30-35.

For electrophysiological recordings (i.e. intrinsic excitability), iPSC differentiation was performed under normal incubator environment (20% oxygen, 5% carbon dioxide) in the presence of mouse astrocytes (1:1) from day 7 onwards, on day 2 media was changed to Neurobasal A and DMEM/F12 1:1 (Stemcell technologies) supplemented with N2 0.5%, NEAA 1%, BDNF 10 ng/ml, NT-3 10 ng/ml (Peprotech), Laminin 0,2 μg/ml (Roche), Culture 1 1% (Gibco), fetal bovine serum 2.5% and 2 μg/ml doxycycline hydrochloride.

For sEPSCs recording instead, we added 1:1 rat astrocytes at day 2, and medium from day 3-7 is Neurobasal Plus medium (NB-Plus) supplemented with B-27 μg/mL (Thermo Fisher Scientific), GlutaMax 10 μg/mL (Thermo Fisher Scientific), primocin 0.1 µg/ml, BDNF 10 ng/ml, NT-3 10 ng/ml (Peprotech) and 2 μg/ml doxycycline hydrochloride. From day 10 onwards, the medium is supplemented with fetal bovine serum 2.5% (Sigma).

### Generation of neural progenitor cells with STEMdiff™ Neural Induction Medium

iPSCs were dissociated with Accutase (GIBCO, Thermo Fisher Scientific) and plated on Matrigel coated plates (final 2% v/v, Corning) directly in STEMdiff™ Neural Induction Medium (#05839, Stem cell technologies) supplemented with 5 μM Y-27632 (Sigma). Cell medium was changed daily till day 5, when the neural progenitor cells were analyzed.

### Cortical organoids generation

Cortical brain organoids were differentiated until day 8 or 50, following the protocol described in (Paşca et al., 2015), with minor modifications to improve its efficiency as we showed previously (López-Tobón et al., 2019). Since we optimized the protocol to avoid the use of mouse embryonic fibroblast (MEF) for stem cell culture, the following procedures were followed: when the hiPSC line reached 80% confluency in a 10 cm dish, colonies were dissociated with Accutase and centrifuged to remove the enzymatic suspension. After resuspension in TeSR/E8 medium supplemented with 5 μM ROCK inhibitor cells were counted with a TC20 automated cell counter (Biorad) and resuspended to get a final concentration of 2 x 10^5 cells/ml. 100 μl/well of cell suspension were seeded into ultra-low attachment PrimeSurface 96-well plates (SystemBio) and then the plates were centrifuged at 850 rpm for 3 minutes to promote the formation of embryoid bodies (EB). The day of the EB generation is referred to as day −2. On day 0 neuronal induction media was added, consisting of of 80% DMEM/F12 medium (1:1), 20% Knockout serum (Gibco), 1 mM non-essential amino acids (Sigma), 0.1 mM cell culture grade 2-mercaptoethanol solution (Gibco), GlutaMax (Gibco, 1:100), penicillin 100 U/ml, streptomycin (100 μg/ml), 7 μM Dorsomorphin (Sigma) and 10 μM TGFβ inhibitor SB431542 (MedChem express) for promoting the induction of neuroectoderm. From day 0 to day 4, medium change was performed every day, while on day 5 neuronal differentiation medium was added, consisting of neurobasal medium (Gibco) supplemented with B-27 supplement without vitamin A (Gibco, 1:50), GlutaMax (1:100), penicillin 100 U/ml, streptomycin (100 μg/ml), 20 ng/ml FGF2 (Thermo) and 20 ng/ml EGF (Thermo) until day 25. From day 25-43 the neuronal differentiation medium was supplemented with 20 ng/ml BDNF and NT3 (Peprotech). From day 43 onwards no growth or neuronal maturation factors were added to the medium.

### Cortical organoids processing and immunostaining

Cortical organoids were harvested on day 18 or 50, fixed overnight at 4 °C in paraformaldehyde 4% PBS solution (SantaCruz). After rinsing with PBS, organoids were embedded in 2% low melting agarose dissolved in PBS; upon agarose solidification, blocks were put in 70% ethanol and kept at 4 °C before paraffin embedding, sectioning, and routine hematoxylin/eosin staining. Deparaffinization and rehydration were achieved by consecutive passages of 5 minutes each in the following solutions: 2 x histolemon (Carlo Erba), 100% ethanol, 95% ethanol, 80% ethanol and water. Sections were then incubated for 45 min at 95 °C in 10 mM Sodium citrate (Normapur)/ 0,05% Tween 20 (Sigma) buffer for simultaneous antigen retrieval and permeabilization; then left to cool for at least 2 hours at RT. To immunolabel the markers of interest, a blocking solution made of 5% donkey serum (ImmunoResearch) in PBS was applied for 30 minutes to the slides, while primary antibodies diluted in blocking solution were subsequently added, performing overnight incubation at 4 °C. Secondary antibodies and DAPI were diluted in PBS and applied to the sections for 1 hour and 5 minutes respectively. After each incubation step, 3 x 5 minutes of washing steps with PBS buffer were performed. After a final rinse in deionized water, slides were air-dried and mounted using Mowiol mounting medium. The following primary antibodies and dilutions were used; KI67 (1:500 Abcam) and PAX6 (1:250 Abcam), CTIP2 (1:500 Abcam).

### Cortical organoids image acquisition and processing

For organoids’ size and circularity assessment; each well was recorded every day for 8 days in the IEO imaging facility with the TIRF microscope. For each well, the ROI corresponding to the area of the organoid was identified and analyzed using a customized pipeline (all code is available as supporting material: TL_analysis_v1.ijm) developed in collaboration with the IEO imaging facility in FIJI (v.1.49 NIH-USA). In brief, ROI were defined with masks using a minimum threshold of signal to noise (Otsu’s threshold), and the “Analyze Particles” function was used to calculate the circularity metric for each ROI.

R software was used to plot the values of organoids’ area and circularity grouping them by days and lines (all code is available as supporting material: OrganoidsGrowthFinal.html).

Image acquisition of 5 μm organoids paraffin sections was done in Leica DM6 B widefield microscope. KI67, PAX6 and CTIP2 quantification was done by measuring markers positive area normalized against DAPI staining using Fiji (v. 2.0.0) software.

### Immunofluorescence in iNeurons

For MAP2B quantification, induced neurons were differentiated in glass coverslips until day 10, 20 and 30. iNeurons were fixed in 4% paraformaldehyde in PBS for 15 min at room temperature immediately after removal of culture medium. iNeurons were then washed 3 times for 5 min with PBS, permeabilized with 0.1% Triton X-100 in PBS for 15 min, and blocked in 5% donkey serum in PBS for 30 min. After blocking, the cells were incubated with primary antibodies diluted in blocking solution overnight at + 4 °C. The cells were washed three times with PBS for 5 min and incubated with secondary antibodies at room temperature for 1 h. Neuronal nuclei were then stained with DAPI solution at room temperature for 5 min. Coverslips were rinsed in sterile water and mounted on a glass slide with 5 μl of Mowiol mounting medium. Primary antibodies used: MAP2B 1:500 (BD Biosciences). Images were acquired with Leica SP8 confocal microscope and processed with Fiji (v. 2.0.0)

#### Sholl analysis

##### Virus preparation

Five million HEK 293T cells were plated in 10 cm plates and grown in 10% fetal bovine serum in DMEM. On the next day, cells were transfected with plasmids for gag-pol (10 μg), rev (10 μg), VSV-G (5 μg) and the target construct (15 μg) CaMKIIα-mKO2, using the calcium phosphate method. On the next day, the medium was changed. The day after, the medium was spun down in a high-speed centrifuge at 30,000g, at 4 °C for 2 h. The supernatant was discarded and 100 μl of PBS were added to the pellet and left overnight at 4 °C. On the next day, the solution was distributed into 10-μl aliquots and frozen at −80 °C.

##### Neuronal infection

At day 4 of the differentiation protocol, neurons were infected with virus bearing CaMKIIα-mKO2; specifically, 10 μl virus per 6cm plate from a standard preparation (see Virus preparation). At this time, an appropriate number of vials of mouse astrocytes were thawed into a 10 cm plate, in order to obtain at least 1.25 millions of astrocytes at day 8. At day 8, infected neurons were digested with accutase for 5 min., washed with PBS, counted, and seeded at a total density of at least 30.000 cells/cm^2^ (300.000 cells/well in a 6-well plate) in a 1:50 ratio with not infected neurons and in a 1:1 ratio with mouse astrocytes, in poly-D-lysine-coated coverslips. Over the following weeks, the coverslips were monitored and those with at least 10 visible individual neurons were kept for image acquisition. Sholl analysis was performed at day 35.

##### Image acquisition

For morphometric analysis, images of neurons were acquired at 10x magnification using the Leica DM6 Multifluo Fluorescence Microscope. Image acquisition was done in a semi-automated manner, with manual picking of individual neurons and batch acquisition. Two channels were acquired per batch for GFP and mKO2.

##### Morphological analysis

All analyses were completed in Fiji (ImageJ). Dendrite length was characterized using the Simple Neurite Tracer ImageJ plugin. For Sholl analysis, the center of concentric spheres was defined as the center of the soma, and a 10 mm radius interval was used. In order to compare the Sholl analysis curves between genotypes, a two-way ANOVA test was performed. P-values < 0.05 were considered statistically significant.

### Western Blot

Proteins were extracted using SDS-buffer (4.8% SDS, 20% glycerol, 0.1 M Tris-HCl, pH 7.5). Cell-pellets were lysed with 5 volumes of boiling SDS-buffer, denaturated for 10 min at 95 °C and sonicated. Protein lysates were centrifuged for 10 min at 13000 rpm, RT and supernatants were quantified using BCA (Pierce BCA Protein assay kit, 23225). Proteins were resolved by and blotted on Transfer membrane (Immobilon-P, Merck Millipore IPVH00010). Membrane blocking (5% BSA/TBS, 0.2% Tween-20 for 1h) was followed by incubation with the specific primary antibody and HRP-conjugated secondary antibodies. Proteins were detected by ECL (Bio-Rad). The antibodies used in the pape are listed in the key resource table.

### RNA extraction, RT-PCR and qPCR

RNA was isolated using the RNeasy mini Kit (QIAGEN) and genomic DNA was removed using the RNase-Free DNase Set (QIAGEN). Retrotranscribed cDNA was obtained from 1 μg of total RNA using the SuperScript VILO Retrotranscription kit from Life Technologies according to the manufacturer’s instructions.

For RT-qPCR analysis, a total cDNA amount corresponding to 5 ng of starting RNA was used for each reaction. TaqMan Fast Advanced Master Mix from Life Technologies and 900 nM TaqMan Gene Expression Assay for each gene analyzed were used in a 10μl volume reaction.

A QuantStudio 6 Flex Real-Time PCR system (Applied Biosystems) was utilized to determine the Ct values. Relative mRNA expression levels were normalized to housekeeping genes and analyzed through the comparative delta-delta Ct method using the QBase Biogazelle software.

### RNA-seq libraries preparation

Library preparation for RNA-seq was done with the Ribo-Zero Total RNA sample preparation kit (Illumina), starting from 250 ng to 1 g of total RNA. The quality of cDNA libraries was assessed by Agilent 2100 Bioanalyzer using the High Sensitivity DNA Kit. Libraries were sequenced with the Illumina HiSeq machine, 50–base pair (bp) paired end with a coverage of 35 millions of reads per sample

### Ribosomal profiling

Libraries of ribosome-protected fragments (RPFs) was generated using the TruSeq Ribo Profile (Mammalian) (Illumina). The experiment was done in three biological replicates for each genotype. Briefly, iNeurons were treated with cycloheximide at 0.1mg/mL for 5 min and lysed according to the manufacturer’s instructions. To generate Ribosome Footprints, cell lysates were digested with TruSeq Ribo Profile Nuclease. The Ribosome Protected fragments were first purified using MicroSpin S-400 columns (GE) and then size-selected (28-30 nt) from 15% polyacrylamide/7-8 M urea/TBE gel. rRNA was removed using the Ribo-Zero Gold Kit (Illumina). After 3’ adaptor ligation and reverse transcription of the library, cDNA (70-80 nt) was purified from 10% polyacrylamide/7-8 M urea/TBE gel. After cDNA circularization and limited amplification (9 cycles), the RPF libraries were purified using an 8% native polyacrylamide gel. The RPF libraries were sequenced on an Illumina HiSeq 2500 sequencer with single-end 50 cycles (SE50) run type.

### MS-based Proteomics

Cell pellets were processed and digested using the PreOmics iST sample preparation kit, following the manufacturer’s guidelines. Peptide mixtures were separated by reversed-phase chromatography on an EASY-nLC 1200 ultra high-performance liquid chromatography (UHPLC) system through an EASY-Spray column (Thermo Fisher Scientific), 25-cm long (inner diameter 75 µm, PepMap C18, 2 µm particles), which was connected online to a Q Exactive HF (Thermo Fisher Scientific) instrument through an EASY-Spray™ Ion Source (Thermo Fisher Scientific). Peptides were loaded in buffer A (0.1% formic acid) and eluted with a linear 135 min gradient of 3–30% of buffer B (0.1% formic acid, 80% (v/v) acetonitrile), followed by a 10 min increase to 40% of buffer B, 5 min to 60% buffer B and 2 min to 95% buffer B, at a constant flow rate of 250 nl/min. The column temperature was kept at 45°C under EASY-Spray oven control. The mass spectrometer was operated in a top-10 data-dependent acquisition (DDA) mode. MS spectra were collected in the Orbitrap mass analyzer at a 60,000 resolution (200 m/z) within a range of 375–1650 m/z with an automatic gain control target of 3e6 and a maximum ion injection time of 20 ms. The 10 most intense ions from the full scan were sequentially fragmented with an isolation width of 2 m/z, following higher-energy collisional dissociation with a normalized collision energy of 28%. The resolution used for MS/MS spectra collection in the Orbitrap was 15,000 at 200 m/z with an AGC target of 1e5 and a maximum ion injection time of 80 ms. Precursor dynamic exclusion was enabled with a duration of 20s. Each sample was injected twice, to increase protein identifications.

The MS data were processed with MaxQuant (Tyanova et al., 2016), using the Uniprot HUMAN (181029) database. Enzyme specificity was set to trypsin and two missed cleavages were allowed. Methionine oxidation and N-terminal acetylation were included as variable modifications and the FDR was set to 1%, both at the protein and peptide level. The label-free software MaxLFQ (Cox et al., 2014) was activated, as well as the “match between runs” feature (match from and to, matching time window=2 min). The mass spectrometry data have been deposited to the ProteomeXchange Consortium (Vizcaíno et al., 2014) via the PRIDE partner repository with the dataset identifier PXD035276.

### Electrophysiological recordings

Patient-derived neurons were differentiated until day 35. Intrinsic neuronal excitability was measured using whole-cell patch-clamp recording in the current-clamp configuration by injecting 500 msec 5 pA current steps from −60 mV as previously described (Pozzi et al., 2013). Patch-clamp recordings were performed in an extracellular solution with the following composition: 130 mM NaCl, 5 mM KCl, 1.2 mM KH2PO4, 1.2 mM MgSO4, 2 mM CaCl2, 25 mM HEPES, and 6 mM Glucose, pH 7.4, in the presence of synaptic transmission blockers, CNQX 10 mM and APV 20 mM. Borosilicate glass pipettes of 4-6 MΩ were filled with the following internal solution (in mM): 135 K-gluconate, 5 KCl, MgCl2, 10 HEPES, 1 EGTA, 2 ATP, 0.5 GTP, pH 7.4. Electrical signals were amplified by a Multiclamp 200 B (Axon instruments), filtered at 5 kHz, digitized at 20 kHz with a digidata 1440 and stored with pClamp 10 (Axon instruments). Passive properties including capacitance and membrane resistance were calculated in voltage-clamp mode using a pulse test of 10 mV. Only neurons with a stable (max deviation 10%) access resistance <15 mW and with a holding current < 100 pA were considered. Intrinsic neuronal excitability was calculated as the total number of action potentials for each current step, whereas the current threshold density was calculated as the minimum depolarizing current needed to elicit at least one action potential. The parameters describing the action potential shape (AP peak and Max Rising Slope) were analyzed using the pClamp software (Molecular Devices) and data were analyzed using Prism software (GraphPad Software, Inc.).

Sodium (I_Na_) and potassium (I_k_) currents were recorded in voltage-clamp mode using a step protocol from −120 mV to + 50 mV at 10 mV increments from a holding potential of −70 mV. Only neurons with a stable access resistance < 5mW (uncompensated) were considered.

Spontaneous excitatory post-synaptic currents (sEPSCs) were measured by 10 min continuous recording at a holding potential (Vh) of 60 mV. sEPSCs were analyzed using MiniAnalysis 6.0.2 (Synaptosoft Inc, GA, USA) as previously described (Mossink et al., 2022).

For the experiments with REST, inhibitor X5050 (Millipore) was dissolved in DMSO and 100uM was added in the neuronal medium for 24 hours before recordings or harvesting neurons for RNA extraction and RNA sequencing.

### OMICs analysis

The code underlying the -omics analyses and related figures, as well as re-usable data objects, are available at https://github.com/plger/7q11ngn2. Unless specified otherwise, the expression heatmaps show log2-foldchanges with respect to the controls of each dataset, with the color scale based on the central 98 percentiles to avoid distortion by outliers, as implemented in the *sechm* package. All data is available at 7q11.23 Explorer, a web server that allows browsing the data: https://ethz-ins.org/7q/

### RNAseq analysis

Quantification was done using Salmon 0.9.1 (Patro et al., 2017) on the ensembl 92 transcriptome. Only protein-coding transcripts were retained, and counts were aggregated at the level of gene symbols. A line from a WBS patient with an atypical deletion was profiled alongside other patient-derived lines (and is deposited), and was included for normalization and dispersion estimates, but was excluded from differential expression and downstream analysis, and not presented in this paper for the sake of simplicity. Only genes with at least 20 reads in at least a number of samples corresponding to 75% of the smallest experimental group were included in the analysis. Differential expression analysis was done using *DESeq2* (Love et al., 2014), and for pairwise comparisons between groups foldchanges were shrunken using the apeglm method (Zhu et al., 2019). In addition, a regression on 7q11.23 copy-numbers was performed. Unless the analysis is specified, DEGs include the union of genes significant across these analyses.

For the merged analysis of isogenic and patient-derived datasets, we first performed surrogate variable analysis to account for technical vectors of variation using the *sva* package (Leek et al., 2012) on variance-stabilized data (as implemented in the *SEtools* package and benchmarked in (Germain et al., 2020), and included the 2 variables in the differential expression model. Unless specified otherwise, genes were considered as differentially-expressed in the merged analysis if they showed at least 30% difference and had an FDR<0.01 (in any of the comparisons), and were additionally differentially-expressed in both datasets with a FDR<0.5.

### Proteomics analysis

Analyses were performed at the level of protein groups, using the median of top 3 peptides. The intensity signals across technical replicates were averaged. Potential contaminants and protein groups with more than 4 missing values per dataset were excluded. Variance-stabilizing normalization and imputation using the *minProb* method were used, as implemented in the DEP package (Zhang et al., 2018). Surrogate-variable analysis was performed before running differential expression analysis via limma/eBayes (Ritchie et al., 2015), using again pairwise comparisons between groups or a regression on copy-numbers.

### Ribosome footprinting analysis

Trimmed reads were mapped with STAR v2.5.2b (Dobin et al., 2013) using the ensembl 92 transcriptome as splice junction guide. One sample (7Dup) was excluded due to poor mapping rate and codon periodicity. Reads mapping to coding sequences were quantified at the gene level using featureCounts v1.5.1 (Liao et al., 2014). Differential expression was performed as described for RNAseq. In addition, differential translation efficiency was assessed by fitting a ∼Replicate+Genotype+SeqType+SeqType:Genotype model using DESeq2, and testing for the interaction terms.

### Integrative analysis across layers

For the integrative analysis of translation, we first restricted ourselves to genes that had an FDR<0.05 and an absolute logFC>0.25 at the transcriptome or proteome, or had aggregated significance (Fisher’s method) across the 3 layers. We then took genes that were significant and in the same direction in both the transcriptome and the proteome, in order to establish the normal relationship between RNA and protein logFC (fitting a robust linear model without intercept) for forwarded genes. The genes were fairly well distributed along the diagonal (r=0.91), and their spread was used to establish a 2-SD interval in which genes were classified as forwarded. For other genes, the residuals were considered as evidence of gene-specific post-transcriptional changes. We next repeated this procedure fitting the residuals on the translation efficiency (TE) logFC for genes with TE p-value<0.01, and again observed a fair correlation (r=0.8). Other genes within the 95% confidence interval around the diagonal were then considered regulated at the level of translation, as their TE pattern explained the RNA-Protein discrepancies.

For the clustering across layers, for each layer and condition, we first scaled the genes’ logFCs by unit-variance (without centering), and assigned a 1 if the median was above 0.2 and all individual logFC were in the same direction, a 0.5 if the median was above 0.2 but individual scores were not in the same direction, and a 0 if below threshold (the same for downregulated genes). We then concatenated the RNA and protein scores, and performed k-means clustering with 8 centers. Similar clusters were then grouped manually.

### Enrichment analysis

Gene Ontology (GO) over-representation analysis on genes found differentially expressed via sequencing was performed using the *goseq* package (Young et al., 2010) to account for length biases, using the genes passing the aforementioned filtering (and with some GO annotation) as background and restricting to terms of annotated to at least 10 and not over 1000 genes. To minimize redundancy in visual representations, significant terms were clustered using k-means (using the elbow of the variance explained plot to choose k) on the binary matrix of gene-term membership. Individual terms were then colored by cluster, and the most significant term of each cluster was used to label the clusters.

### Transcriptome-based staging

Since staging against a reference dataset requires relative rather than absolute transcriptome similarity, and technical differences between profiling methods can often distort similarity estimates, we employed a correction method before assessing similarity. For each gene, the minimum distance between the (edgeR-normalized) logCPM of any sample of the query dataset and that of any sample of the reference dataset was removed from the query dataset before similarity calculation. Similarity was then calculated using Pearson correlation.

For the reference datasets, we used the original quantification and clustering from the respective authors. For the comparison to the Lin et al. Ngn2 dataset, we downloaded the single-cell expression and annotation data from ArrayExpress, and excluded clusters 5 and 8 as astrocytes, as well as cells in timepoints other than the first from clusters displaying high expression of pluripotency genes. Pseudo-bulk profiles were then created by summing the counts of remaining cells from each timepoint, and similarity was calculated as described above.

### Master regulator analysis

The master regulator analysis was based on the DoRothEA v0.0.25 regulons, assigning interaction likelihoods weighted according to the approximative AUC of the different interaction categories in their benchmark (Garcia-Alonso et al., 2019). Per-sample transcription factor (TF) activity was then estimated using the *viper* package (Alvarez et al., 2016) based on the sva-corrected data (if applicable), and differential activity assessed using *limma*, blocking for dataset effects. In addition, a differential TF activity was performed on the RNAseq differential expression statistic using *msviper*, and only TFs that were consistent between the two analyses were considered.

## Acknowledgments

This work received funding from the EPIGEN Flagship Project of the Italian National Research Council (CNR) (to G.T); the European Research Council (ERC DISEASEAVATARS no.616441 to G.T. and R.A.); ERC PoC 713652–LSDiASD to G.T.); EDC-MixRisk, European Union’s Horizon 2020 research and innovation programme (Grant No 634880. to G.T.); ENDpoiNTs, European Union’s Horizon 2020 research and innovation programme (Grant No 825759. to G.T.); Telethon (GGP14265 and GGP19226 to G.T.); Fondazione Umberto Veronesi fellowships (to P.L.G., R.S. and G.D.A.); Ministero della Salute, RC 2019 – ERANET NEURON RRC-2019-2366750 - ALTRUISM (to M.M.); ERANET Neuron - AUTISYN (to P.L.G.); R01GM137031 from National Institutes of Health (NIH) (to Y.L); AIRC IG 16722 (to A.N). AIRC IG No. 21834 and EPIC-XS project No. 823839, funded by the Horizon 2020 programme of the European Union (to T. B.); The Leverhulme Trust and The Horizon 2020 Innovative Training Network EpiSyStem (Marie Skłodowska-Curie Actions; to A.S.); PRIN (Ministero dell’Istruzione, dell’Università e della Ricerca, 2017A9MK4R to M.Matteoli). We are grateful to Federica Pisati from IFOM tissue processing facility, IEO Genomic Unit team, IEO Imaging Facility team, Filippo Mirabella (Humanitas University) for technical assistance with the electrophysiological recordings. We are grateful to Elena Signaroldi for the laboratory management.

We wish to thank the University of Perugia, the University of Antwerp, the University of Sheffield, the Wellcome Trust Sanger Institute, the Telethon Biobank and the Genomic and Genetic Disorders Biobank at the IRCCS Casa Sollievo della Sofferenza for providing the patient-derived fibroblasts or iPSC lines. The authors gratefully acknowledge the research participants who donated biopsies and/or tissue samples from themselves and their family members.

## Author contributions

Conceptualization and study design, M. Mihailovich, P-L.G., R.S. and G.T.; generation of isogenic lines, M. Mihailovich, E.T., G.D.A., A.F. and S.T..; FISH analysis, S.F. and A.N.; PB lines derivation, M. Mihailovich, R.S. and S.T.; generation of iNeurons, M. Mihailovich, R.S, E.T., F.T., S.T. and A.S.; aCGH, I.B.; genotype analysis, A.V. and D.C.; proteomics, Y.L. and R.A (patient-derived iNeurons), R.N. and T.B. (isogenic iNeurons); organoids differentiation and imaging, M.T.R., N.C., R.S.; Taqman assays, Sholl analysis, T.F.; MAP2B staining in iNeurons, D.A.; experimental work, S.R.; adaptation of iNeurons differentiation protocol for electrophysiology, S.T., R.S; electrophysiology experiments (intrinsic excitability), D.P. and M. Matteoli; electrophysiology experiments (sEPSC), U.C. and N.N.K.; RPF and protein synthesis related experiments, M. Mihailovich; REST-experiments, R.S; data analysis, P-L.G., G.D.A. and R.S. (electrophysiology and organoids); manuscript writing, M. Mihailovich, R.S., P.L.G. and G.T.; Supervision, G.T.

## Data and code availability

The mass spectrometry proteomics data have been deposited to the ProteomeXchange Consortium via the PRIDE partner repository with the dataset identifier PXD035276.

## Supplemental Figures

**Fig. S1.**
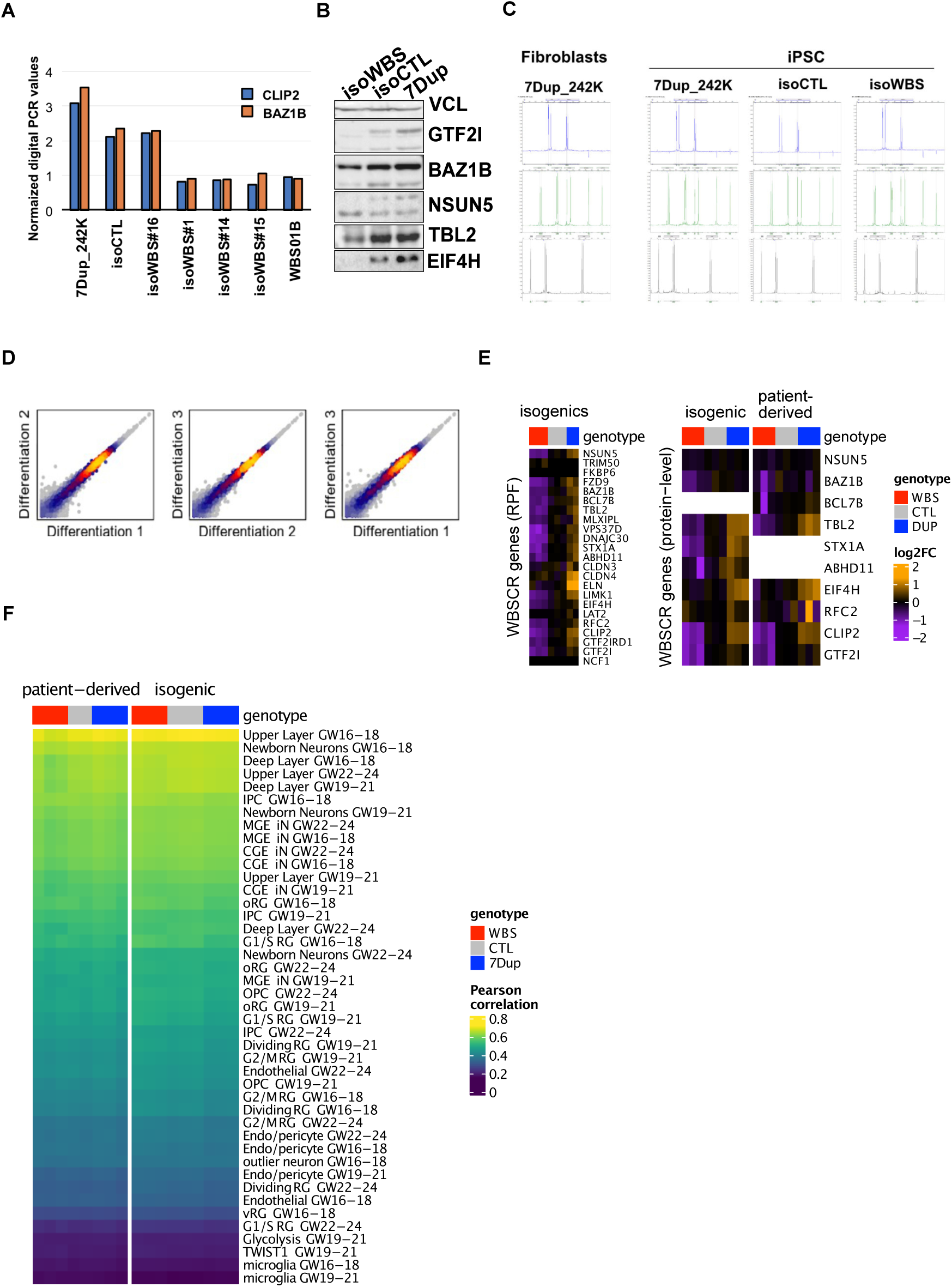
7q11.23 isogenic lines contain a large duplication on Chr14. **(A)** Digital PCR validation of isogenic lines. The analysis was done on DNA from the original patient-derived 7Dup line (242K), isoCTL, several isoWBS clones and one patient-derived WBS line (01B) as a positive control. The probes were designed for two genes from WBSCR, *CLIP2* and *BAZ1B*. **(B)** Western blot analysis for several WBSCR genes in iPSC showed that the dosage is maintained at the protein level. **(C)** Short tandem repeats (STR) analysis confirmed the identity of generated isogenic lines. In addition to the original iPSC 7Dup patient line (242K), we also used the fibroblast from the patient as a control. **(D)** PiggyBac system allows for excellent reproducibility across different rounds of neuronal differentiation (mean logCPM correlation of 0.997). **(E)** Relative expression of WBSCR genes in iNeurons, at the level of Ribosome-protected fragments (RPF, left) and protein (right). All detected genes are shown in genomic order. **(F)** Correlation of iNeurons log(CPM) values with cluster averages from single-cell RNA-seq of human fetal neocortex development (Mayer et al., 2019), after correcting for technical differences.

**Fig. S2.**
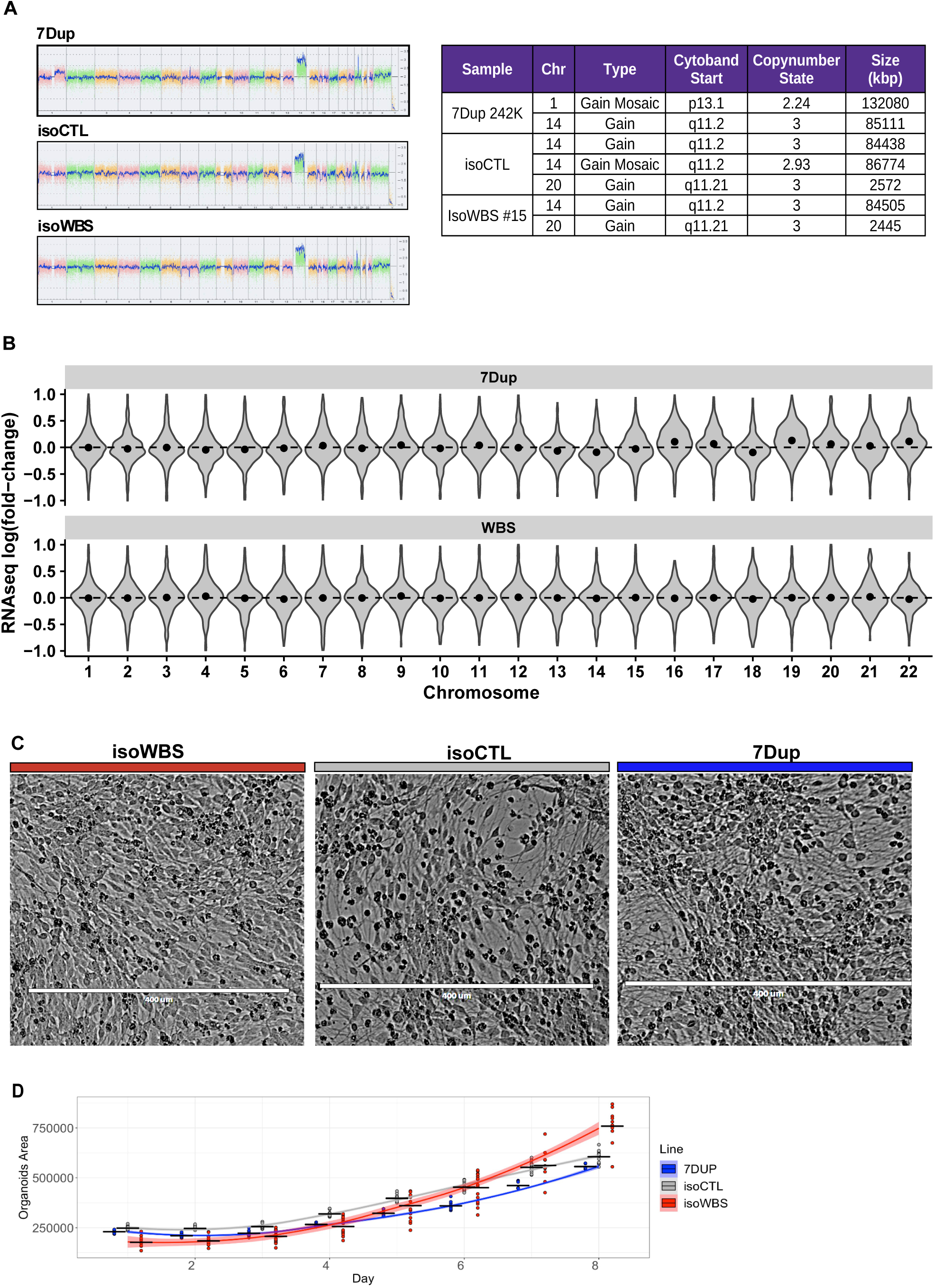
Structural characterization of isogenic cortical organoids. **(A)** Karyostat analysis of DNA isolated from 30 days old induced neurons from the isogenic lines show duplication of Chr14 in all three genotypes (left). The table with genomic alterations identified in the karyostat analysis (right). **(B)** Fold-change distributions by chromosome in 7Dup vs isoCTL and WBS vs isoCTL do not show appreciable differences in the expression of Chr14. **(C)** Brightfield images of isogenic iNeurons on day 14. While isoWBS iNeurons show round cell bodies, isoCTL and 7Dup already have thinner and elongated neurites. **(D)** The plot of the region of interest (ROI) was measured for each organoid across the 8 consecutive days of acquisition. ROI measurements are grouped and colored by line. Black cross bars indicate the mean area value for each line on a corresponding day. Smoothed conditional means are plotted for each line.

**Fig. S3.**
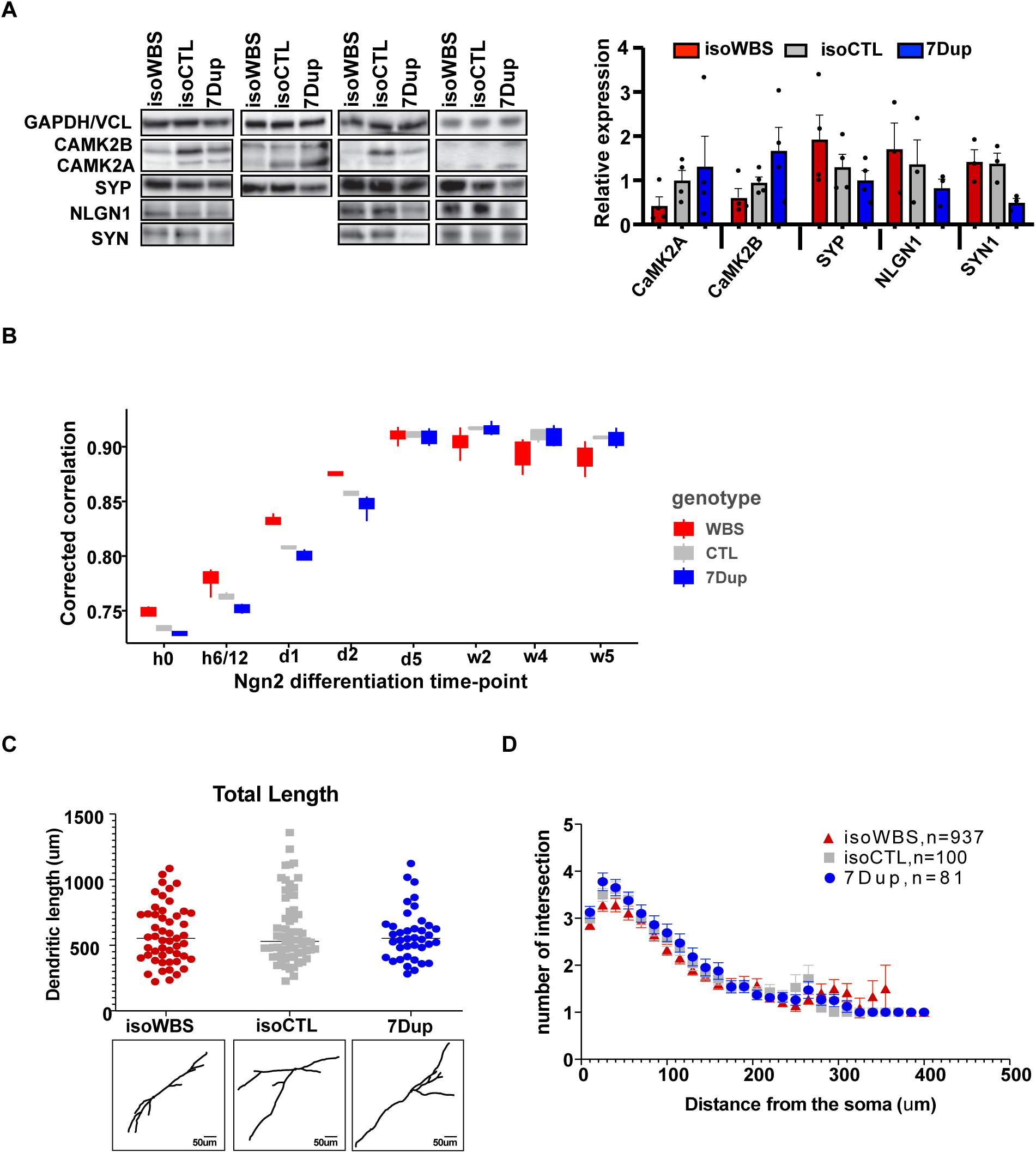
iNeurons resemble best upper layer neurons GW16-18. **(A)** Western blot analysis of selected synaptic proteins done on different rounds of neuronal differentiation (upper panel). The quantification is shown in the right panel. **(B)** Correlation of the iNeurons transcriptomes with a time-course of Ngn2 differentiation (Lin et al., 2021). WBS lines are more correlated than controls to the earlier timepoints, and less correlated with later timepoints, suggesting a differentiation delay. In contrast, 7Dup lines are slightly less correlated with earlier time points, but not significantly more correlated with later time points. **(C)** Quantification of the total dendritic length in iNeurons (mean ± S.D, one-way ANOVA) on the left, shows no statistically significant differences. On the right, representative images of tracings from isoWBS, isoCTL and 7Dup iPSC-derived neurons. **(D)** Quantification of dendritic complexity by Sholl analysis of all orders of branches in isoWBS, isoCTL and 7Dup neurons revealed no significant alterations in dendritic morphology and branching (mean ± S.D., two-way ANOVA followed by Bonferroni multiple comparisons test).

**Fig. S4.**
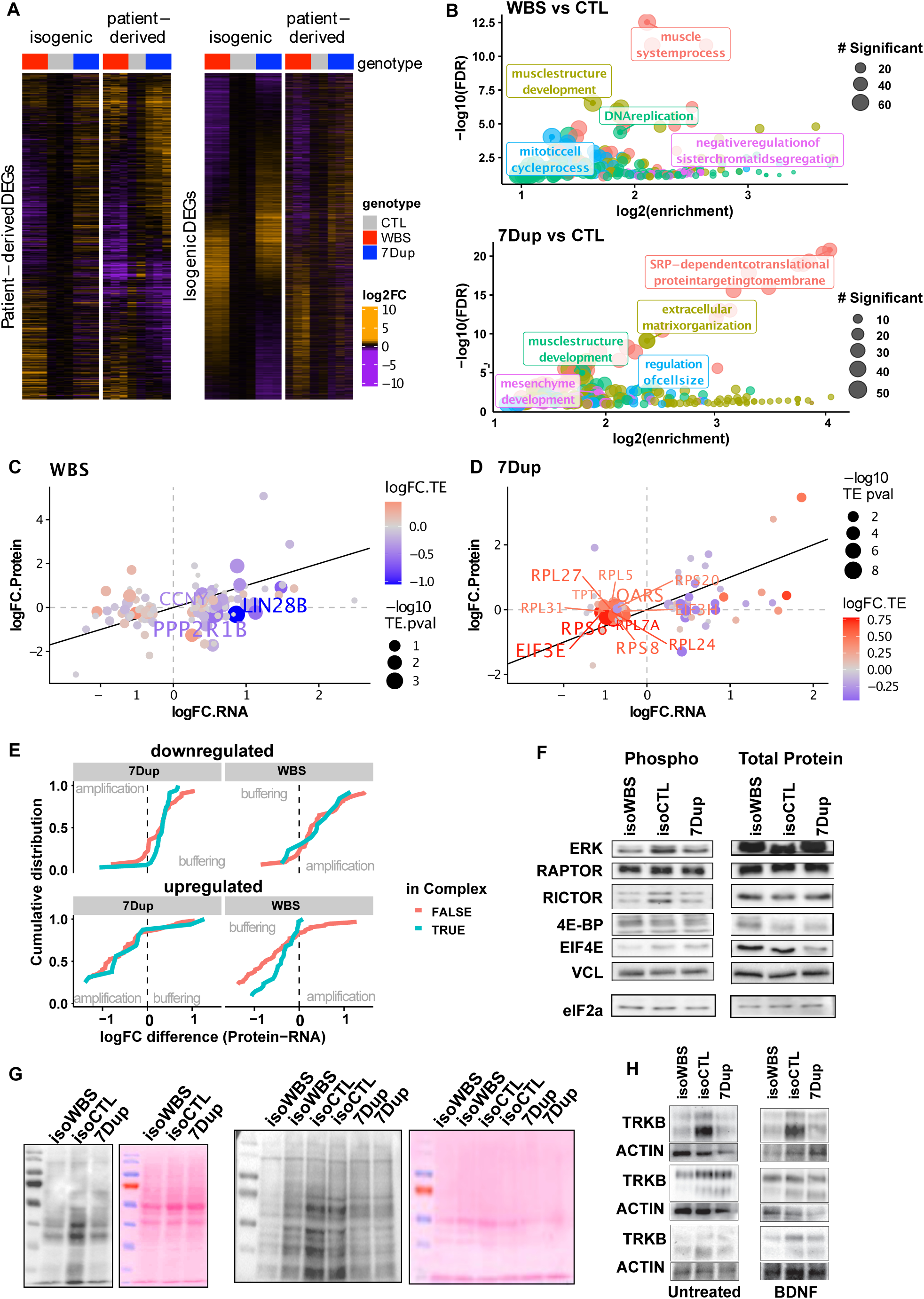
Shared downregulation of protein synthesis and ERK/mTOR activity in WBS and 7Dup. **(A)** Heatmaps showing the fold-changes, in each system, of the DEGs identified in patient-derived lines (left) or isogenic lines (right) alone. **(B)** Enriched Gene Ontology (GO) terms in the WBS vs CTL (upper panel) and 7Dup vs CTL (lower panel). Similar terms are clustered (denoted by colors) and only the top term per cluster is shown. **(C)** Significant differential translation efficiency of specific genes in WBS, using the integrated analysis across the three layers. **(D)** Significant differential translation efficiency of specific genes in 7Dup. **(E)** Cumulative distribution plots of the differences between RNA and Protein log-fold changes split by genotype and direction of the transcriptomic change. The fact that most of the distribution show a positive difference for downregulated genes and a negative one for upregulated genes is consistent with an overall mild buffering. Genes known to form complexes, which have been reported in other contexts to undergo buffering through complex-mediated stabilization and excess degradation, showed only a slightly stronger effect, especially among genes downregulated in 7Dup or upregulated in WBS. **(F)** Replicates of western blot analysis of total and phosphorylated, active, forms of major components of the mTOR pathway, shown in Fig. 3F. Vinculin (VCL) was used as a loading control. **(G)** Additional replicates of western blot analysis of puromycin incorporation assay used for the quantification shown in Fig. 4A. **(H)** Western blot profiling of the TRKB receptor in untreated conditions and after 20 min stimulation with BDNF are shown. Actin was used as a loading control.

**Fig. S5.**
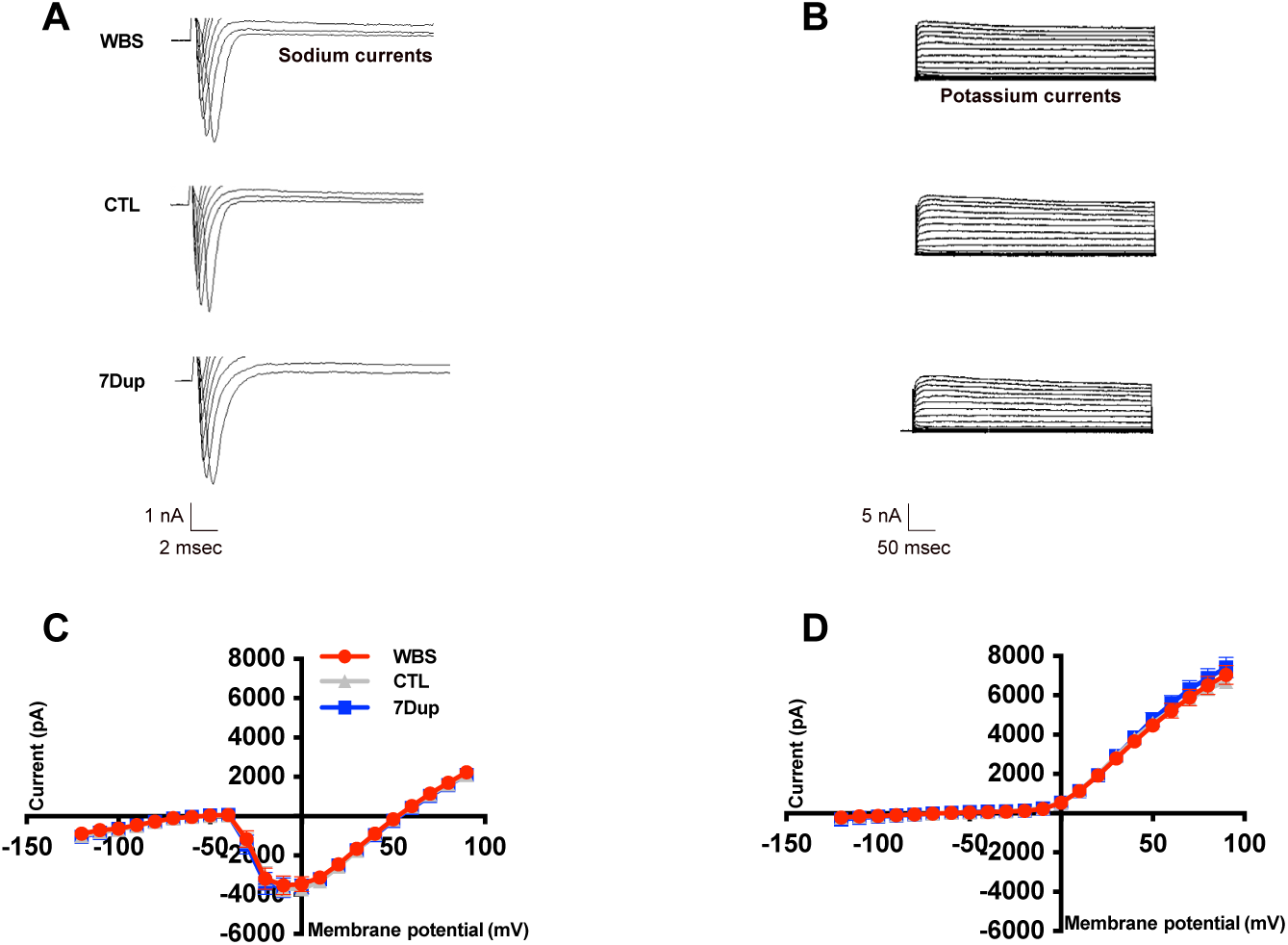
Sodium and potassium currents in patient-derived iNeurons. **(A-B)** Representative sodium and potassium current traces in patient-derived iNeurons from the 3 genotypes recorded in voltage-clamp mode using a step protocol from −120 mV to + 50 mV at 10 mV increments from a holding potential of −70 mV. **(C-D)** Line diagrams of the sodium and potassium currents, respectively. WBS n=45 neurons, CTL n=41, 7Dup n=39. Data are shown as mean ± SEM. All the data is the average of 3 independent experiments. Comparisons were done using two-way ANOVA followed by Tukey’s multiple comparison test. Significance level was set at p<0.05. *p<0.05; **p<0.01.

**Fig. S6.**
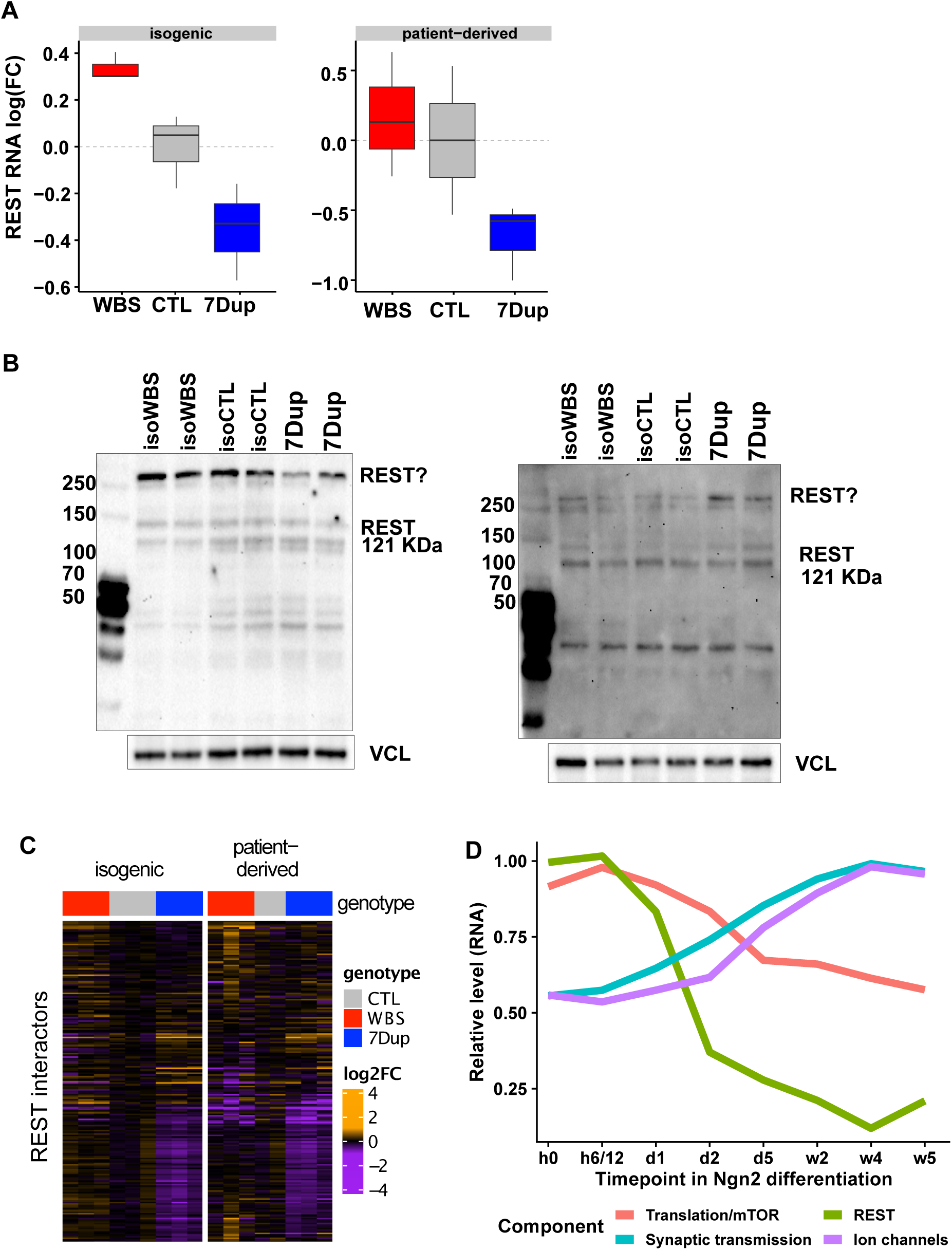
Large transcriptional changes in 7q11.23-dependent manner of the REST-interactome. **(A)** RNAseq analysis of REST mRNA expression in isogenic (left) and patient-derived iNeurons (right). **(B)** REST is buffered at the level of protein in all three genotypes. Predicted REST MW is 121 KDa, however, it is sometimes detected at 200 KDa. In iNeurons we reproducibly detected bands above 250 KDa (marked with REST?), around 120 KDa (marked with REST), and around 40 KDa. one of identified bands consistently was changed between genotypes. Vinculin (VCL) was used as loading control. **(C)** Heatmap showing changes in expression of the REST-interactome in a 7q11.23-dependent manner. **(D)** Relative expression of *REST* and relevant sets of genes during Ngn2-driven differentiation (based on the single-cell sequencing from (Lin et al., 2021).

**Supplementary Table S1 related to Fig.S2.**
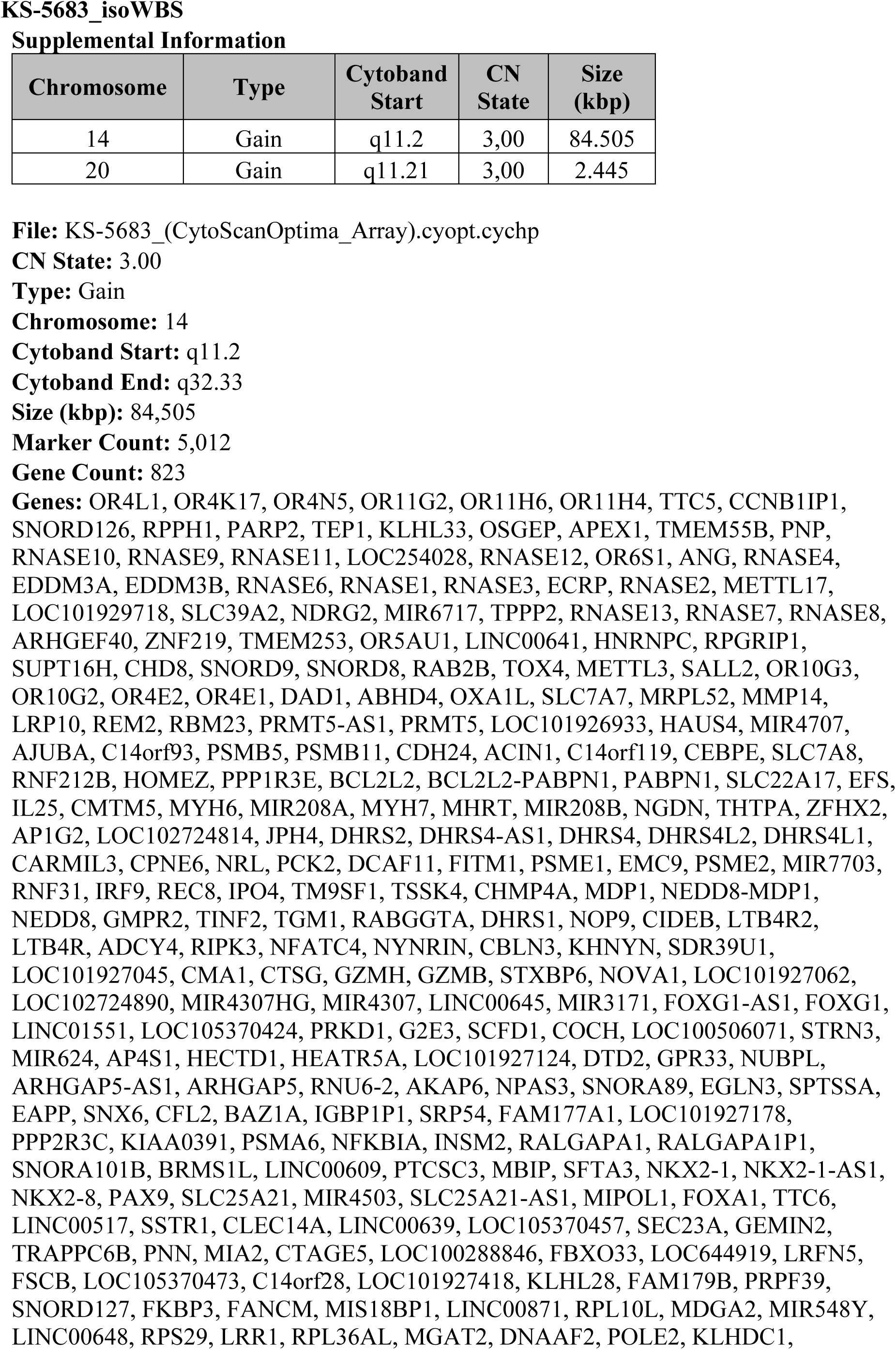
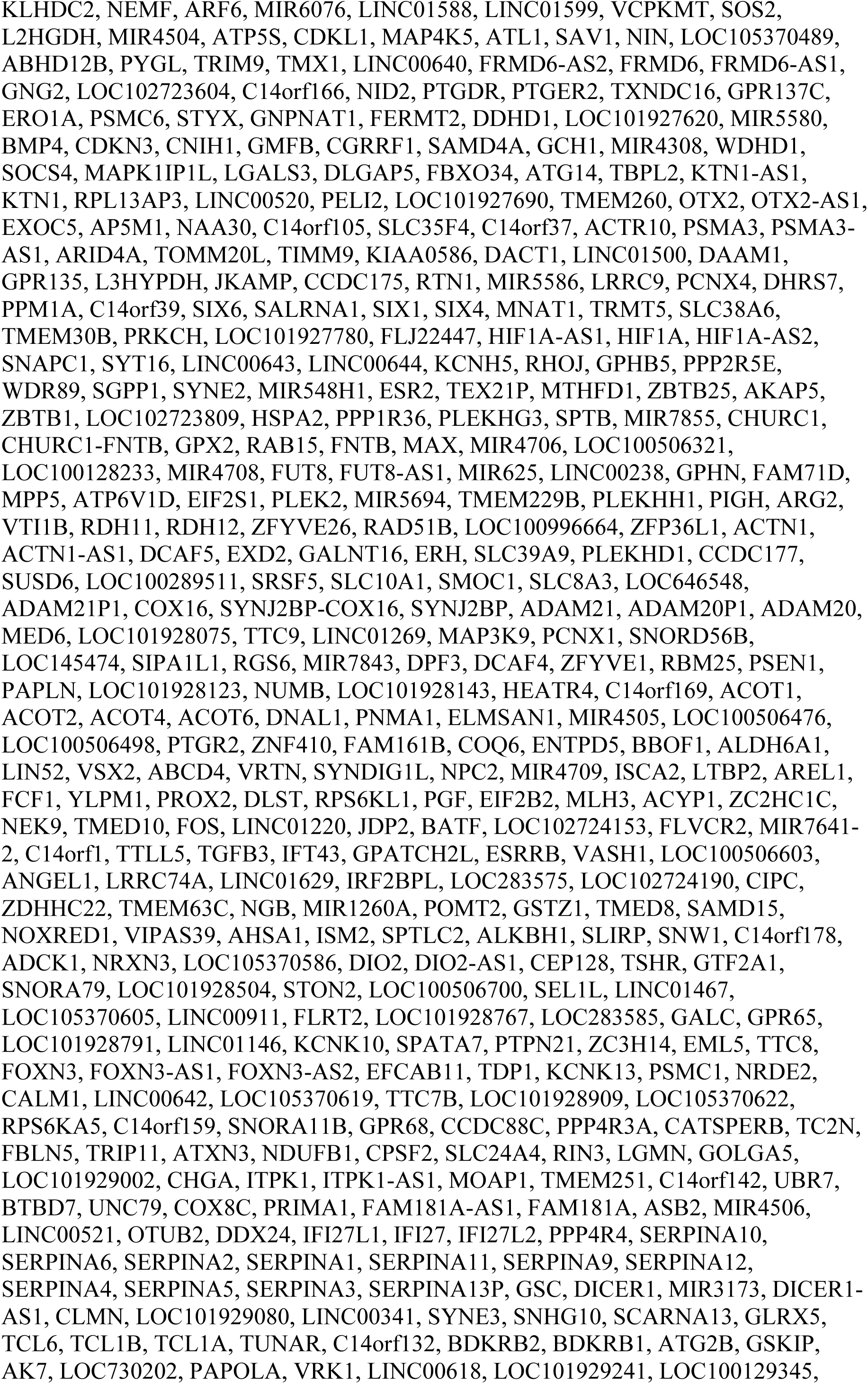
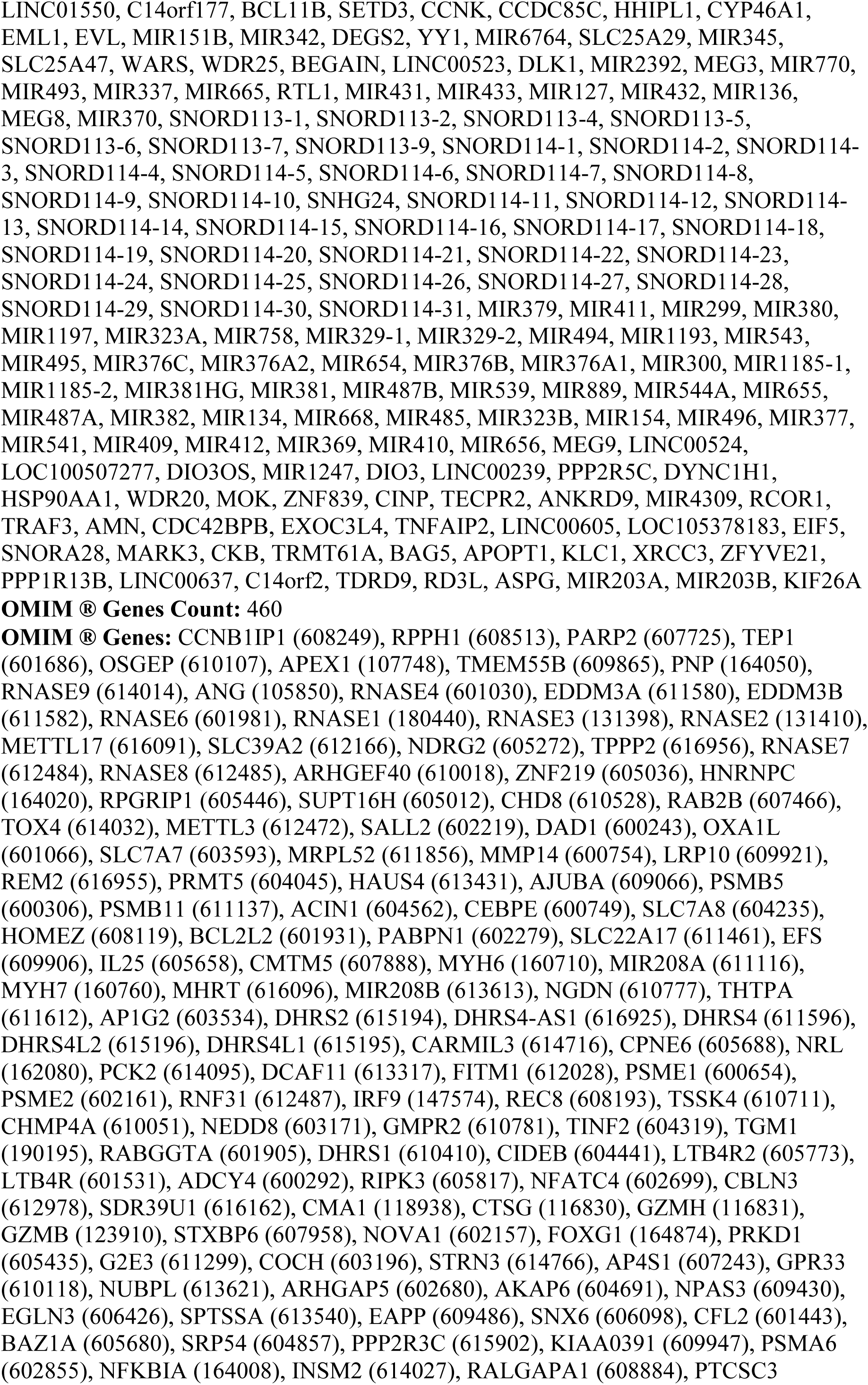
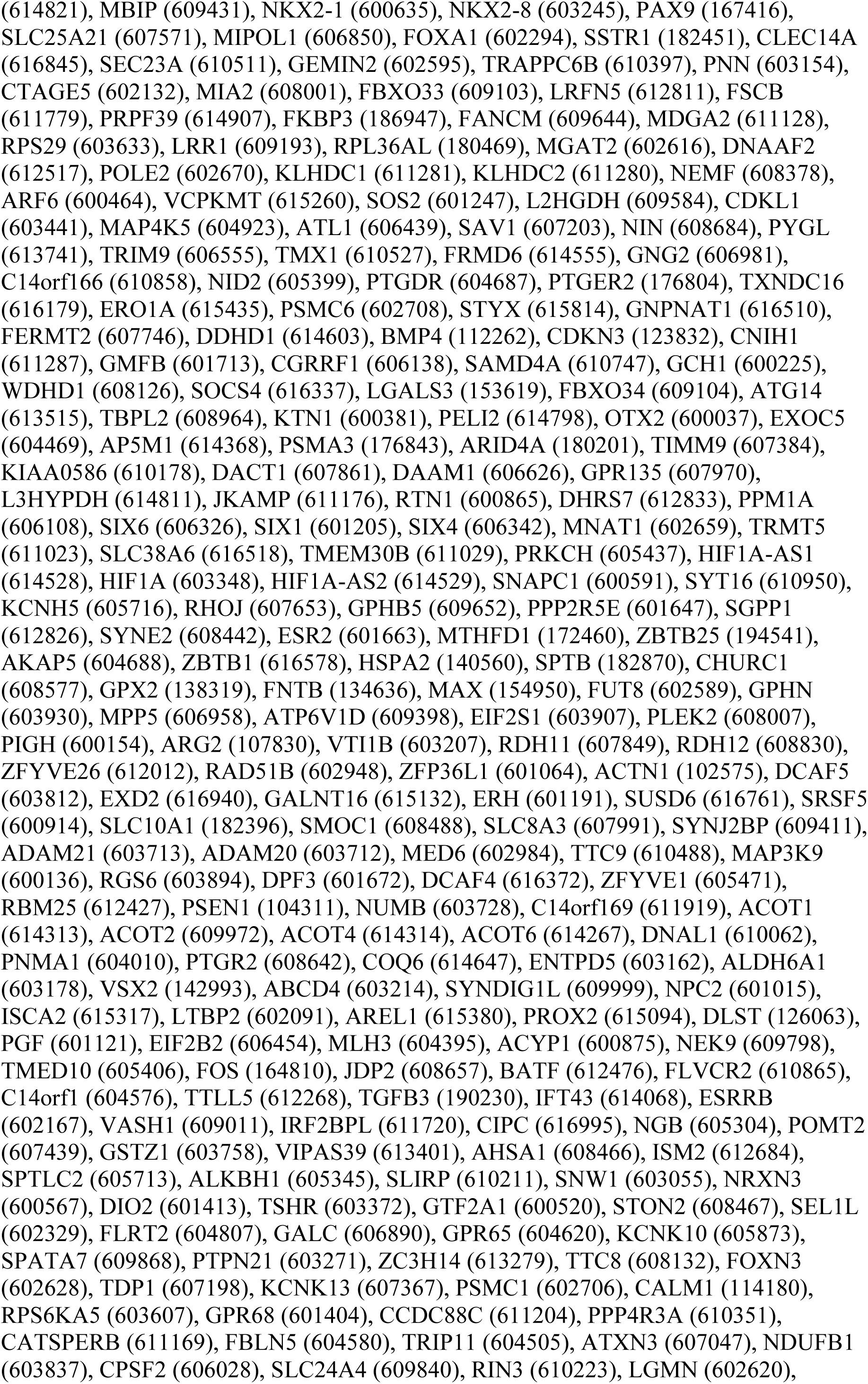
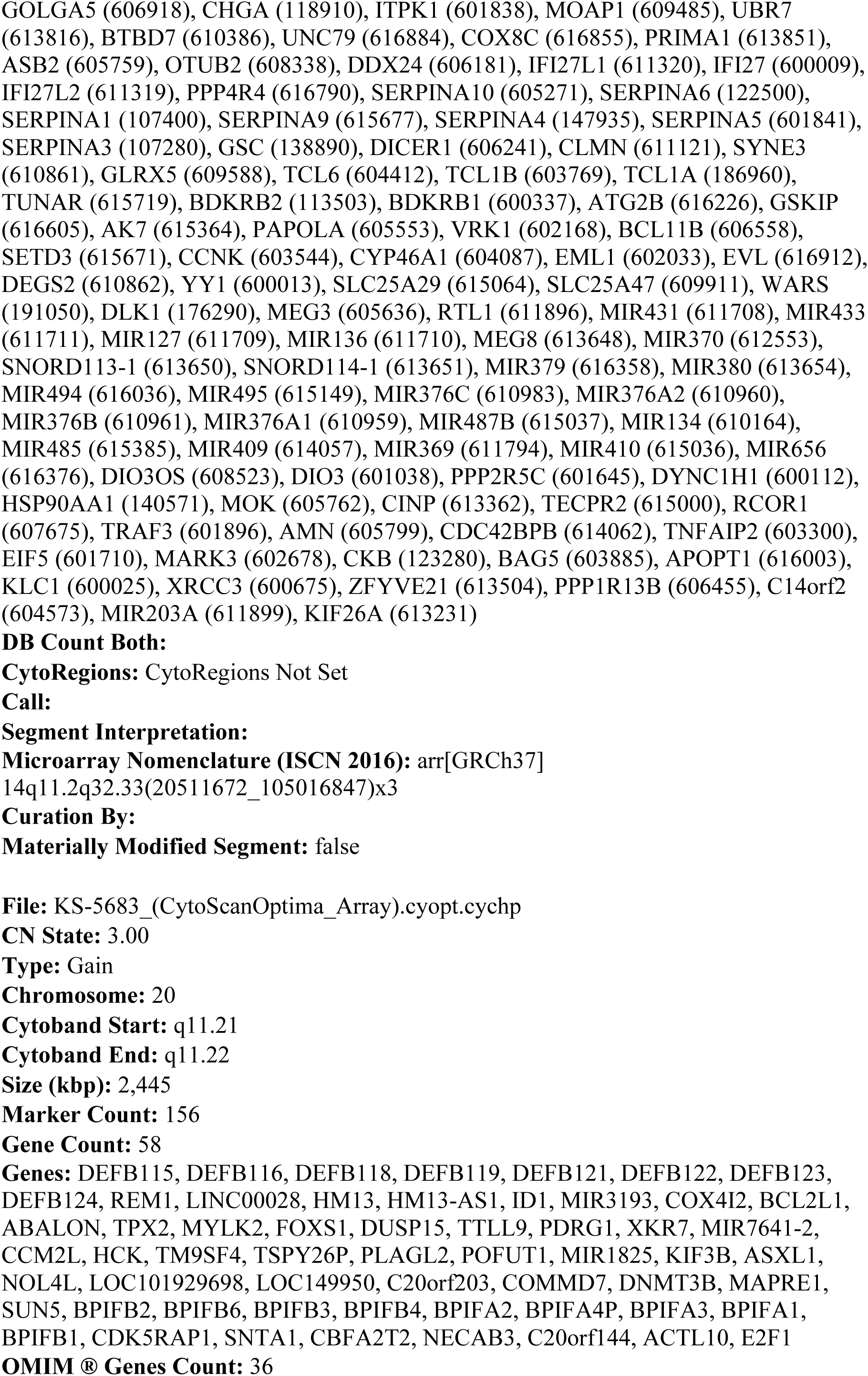
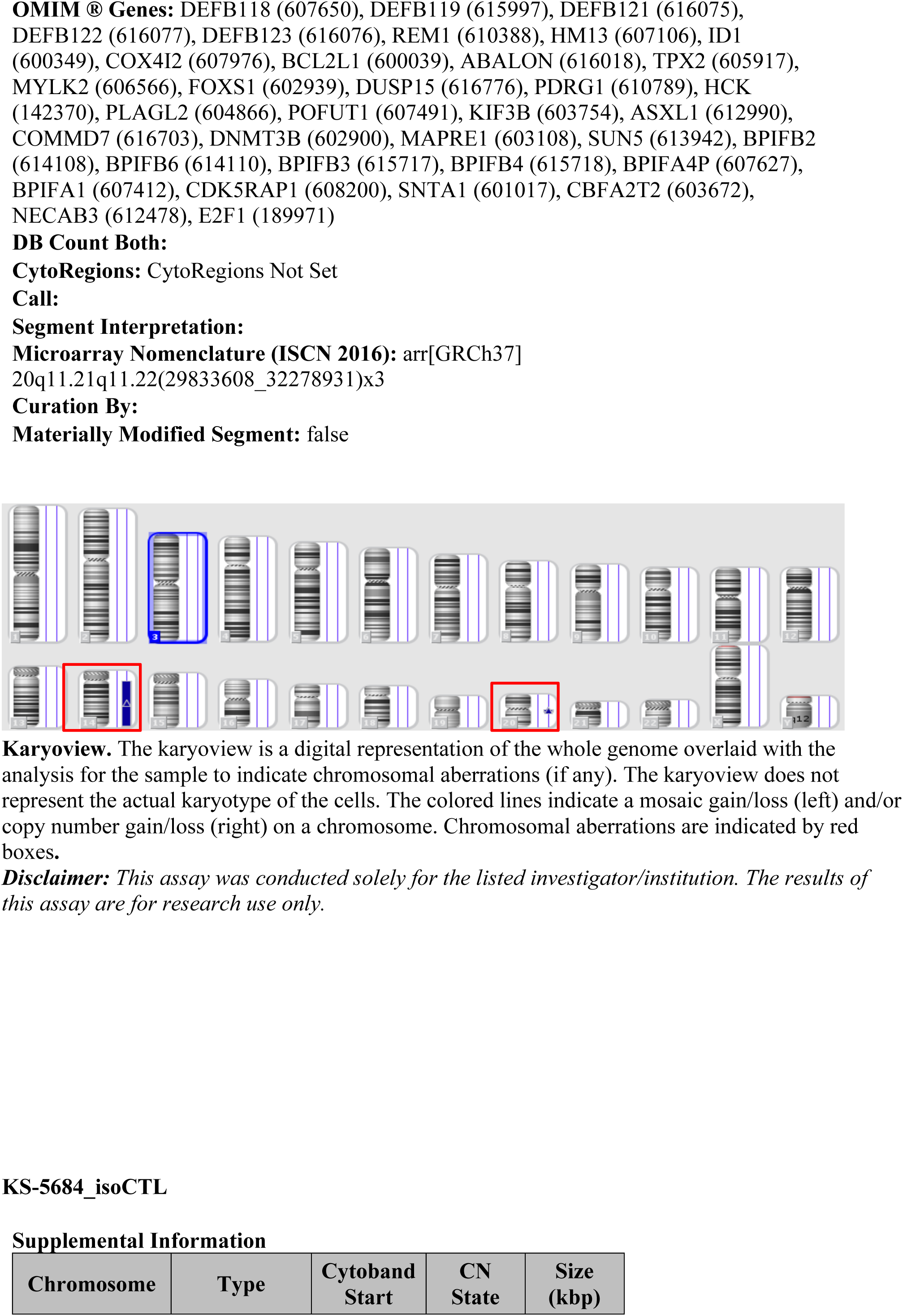
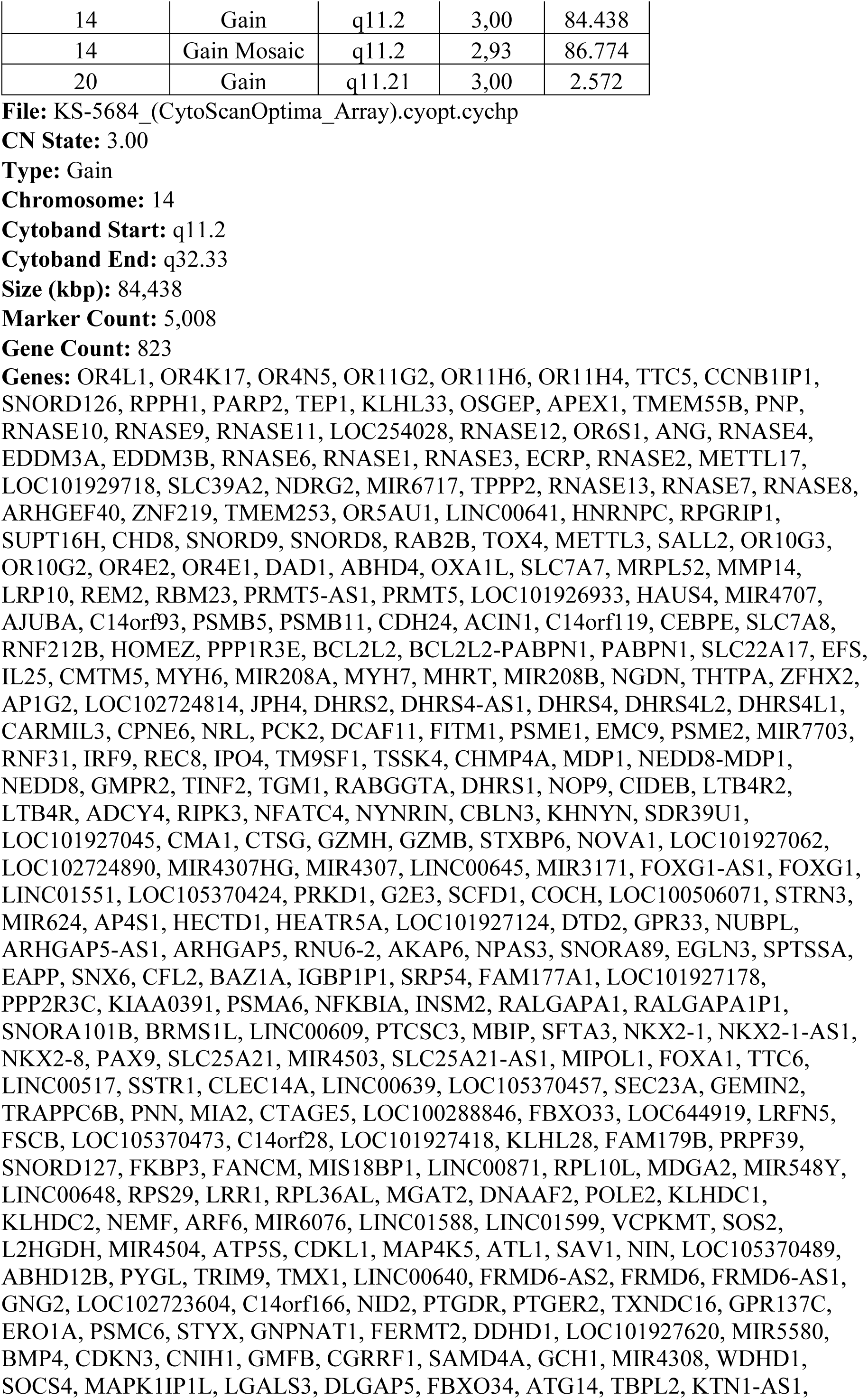
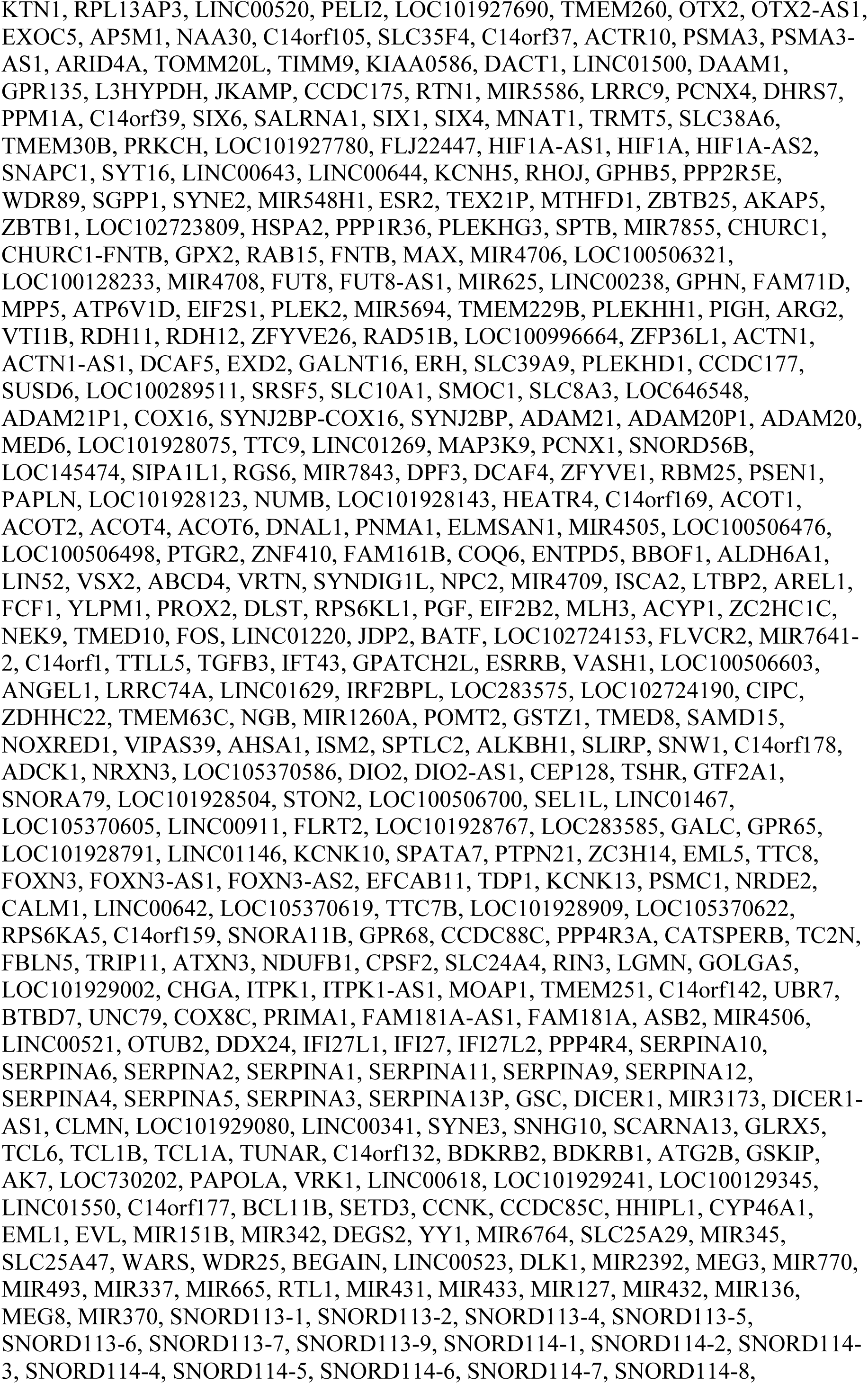
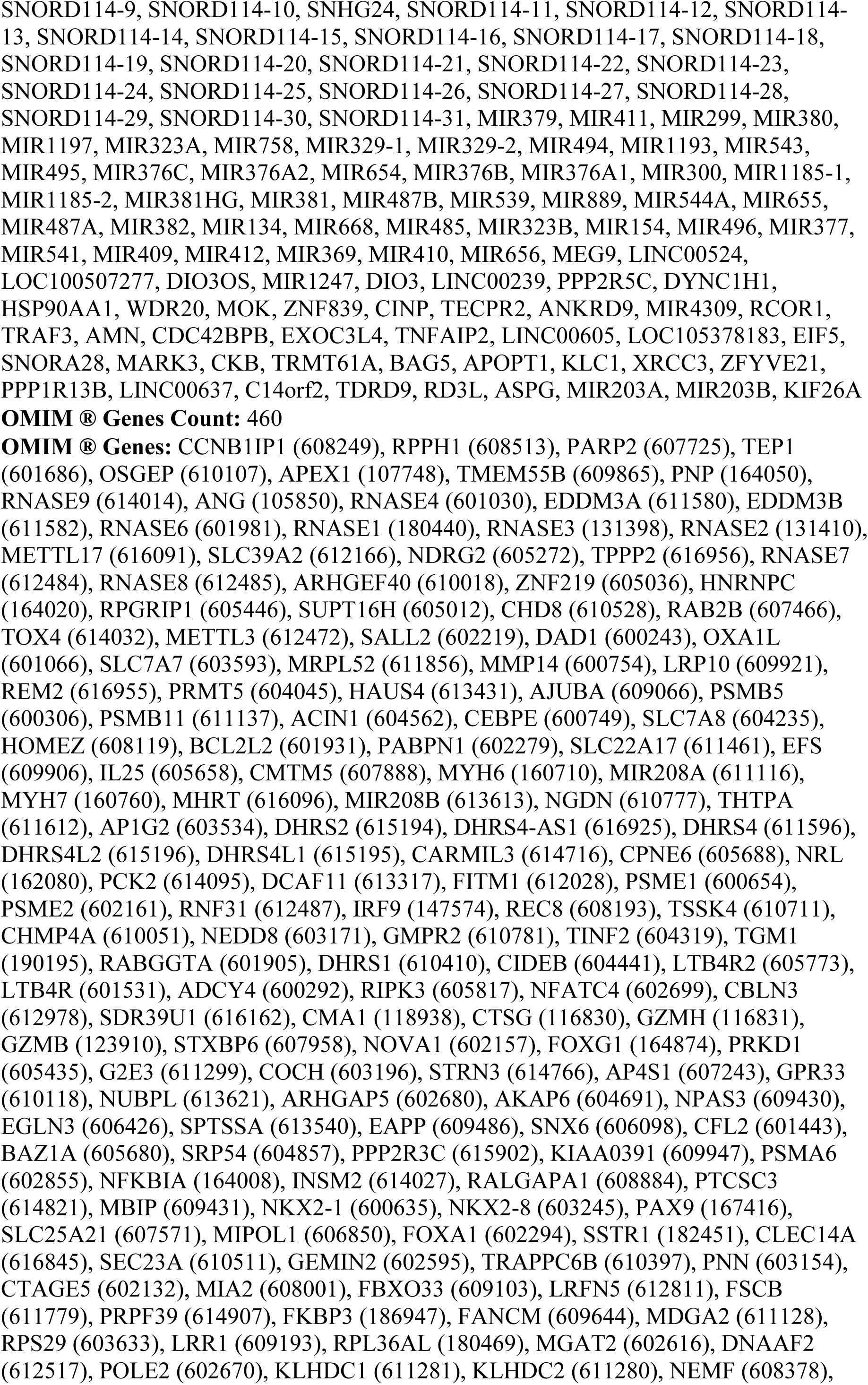
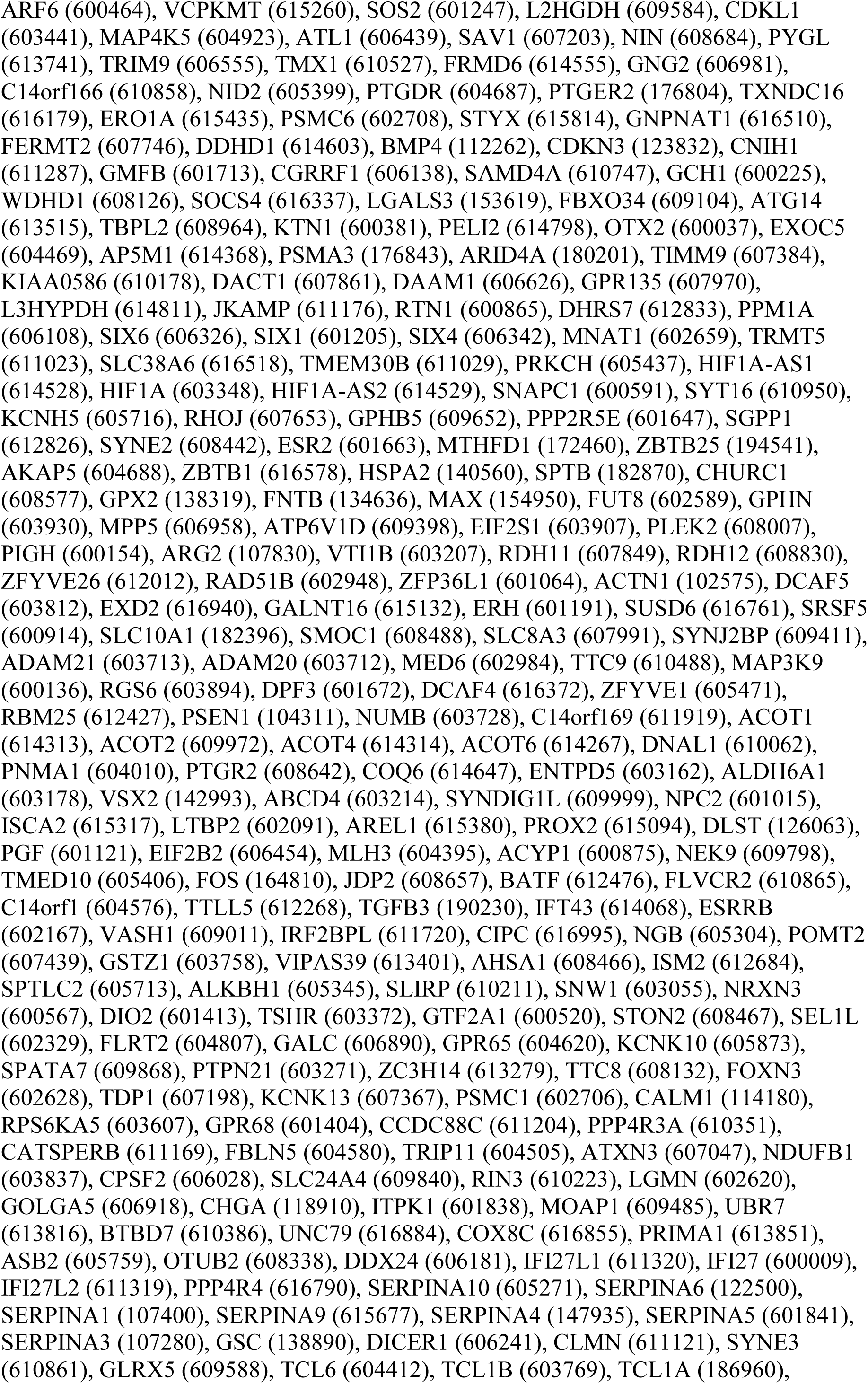
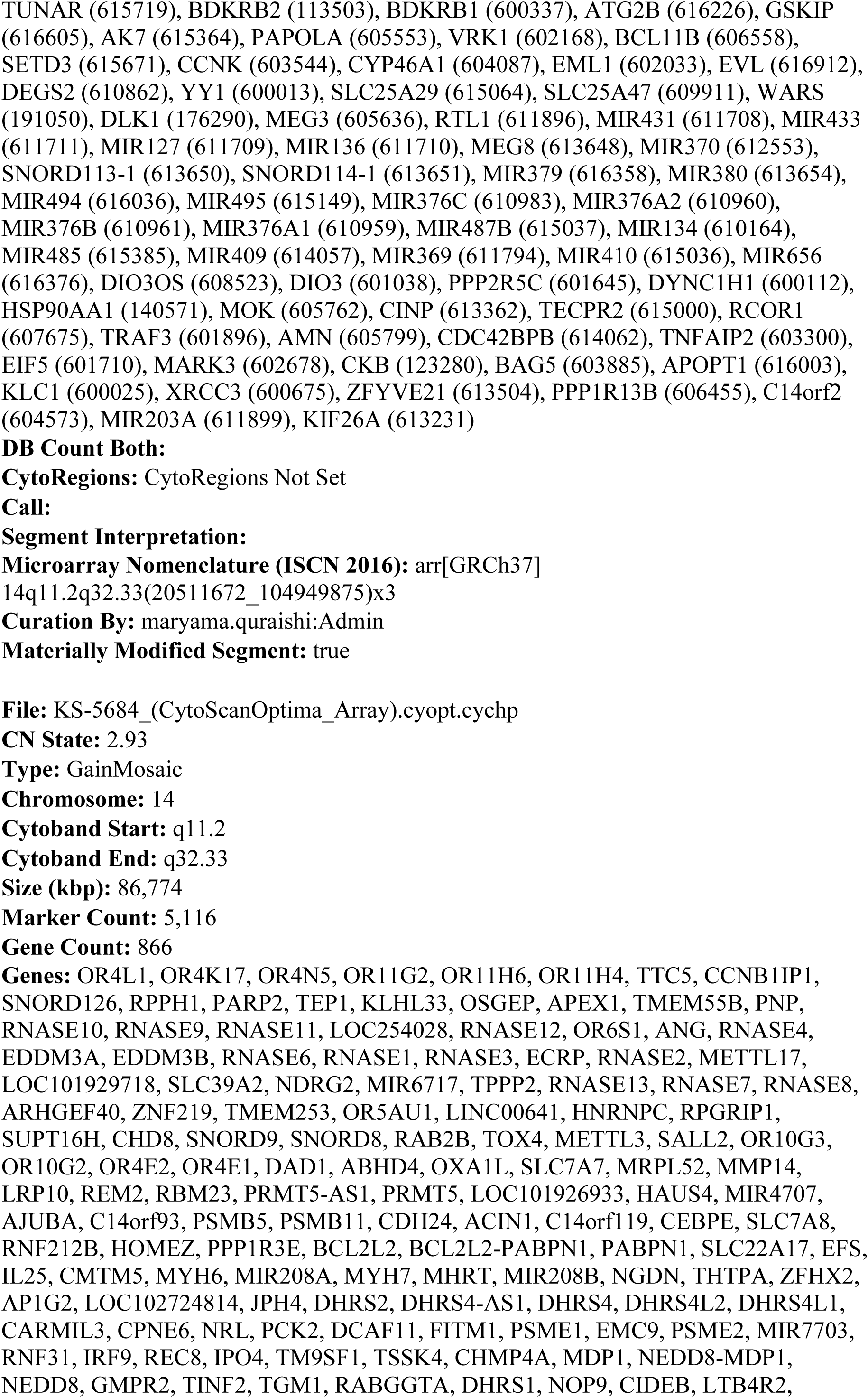
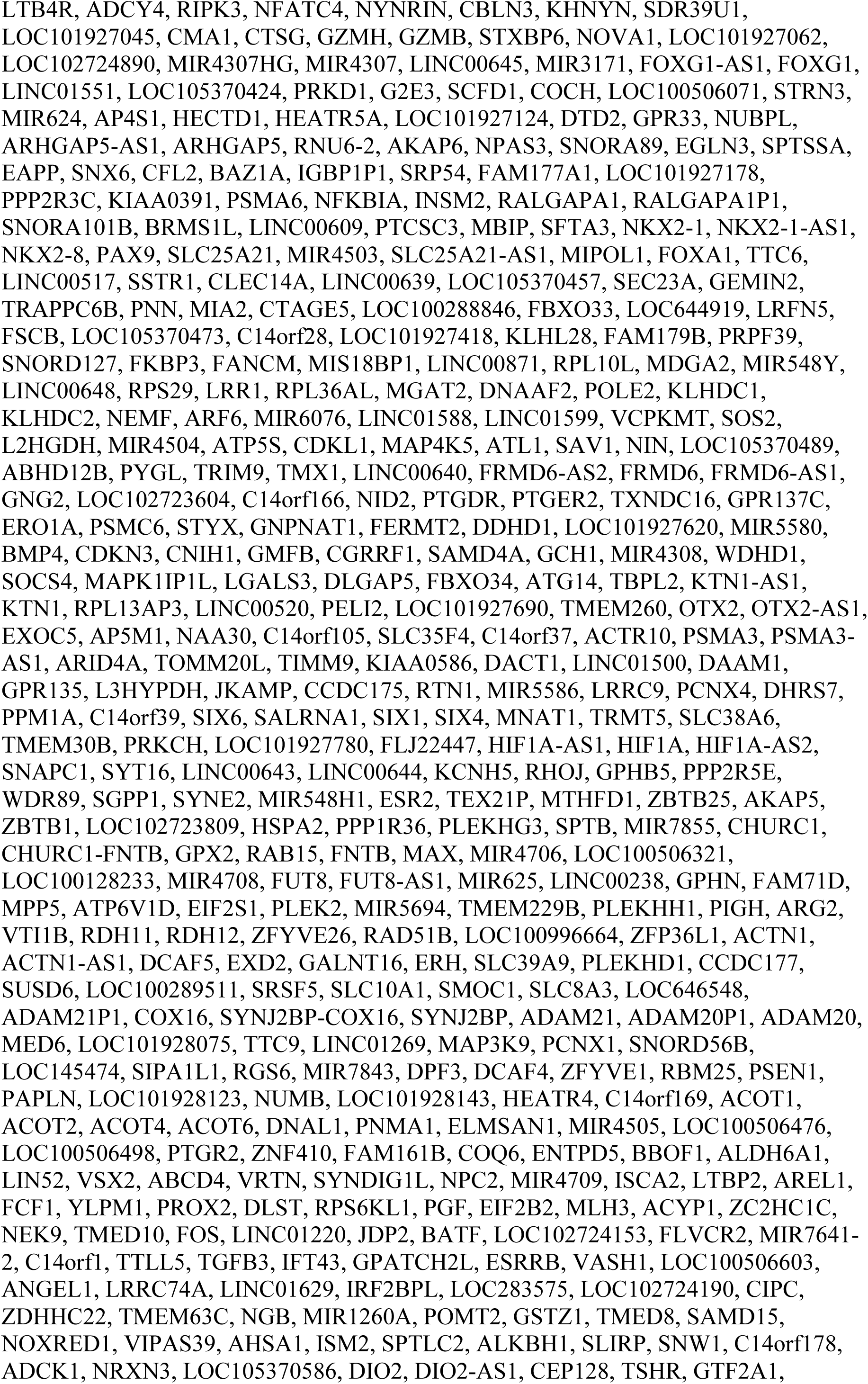
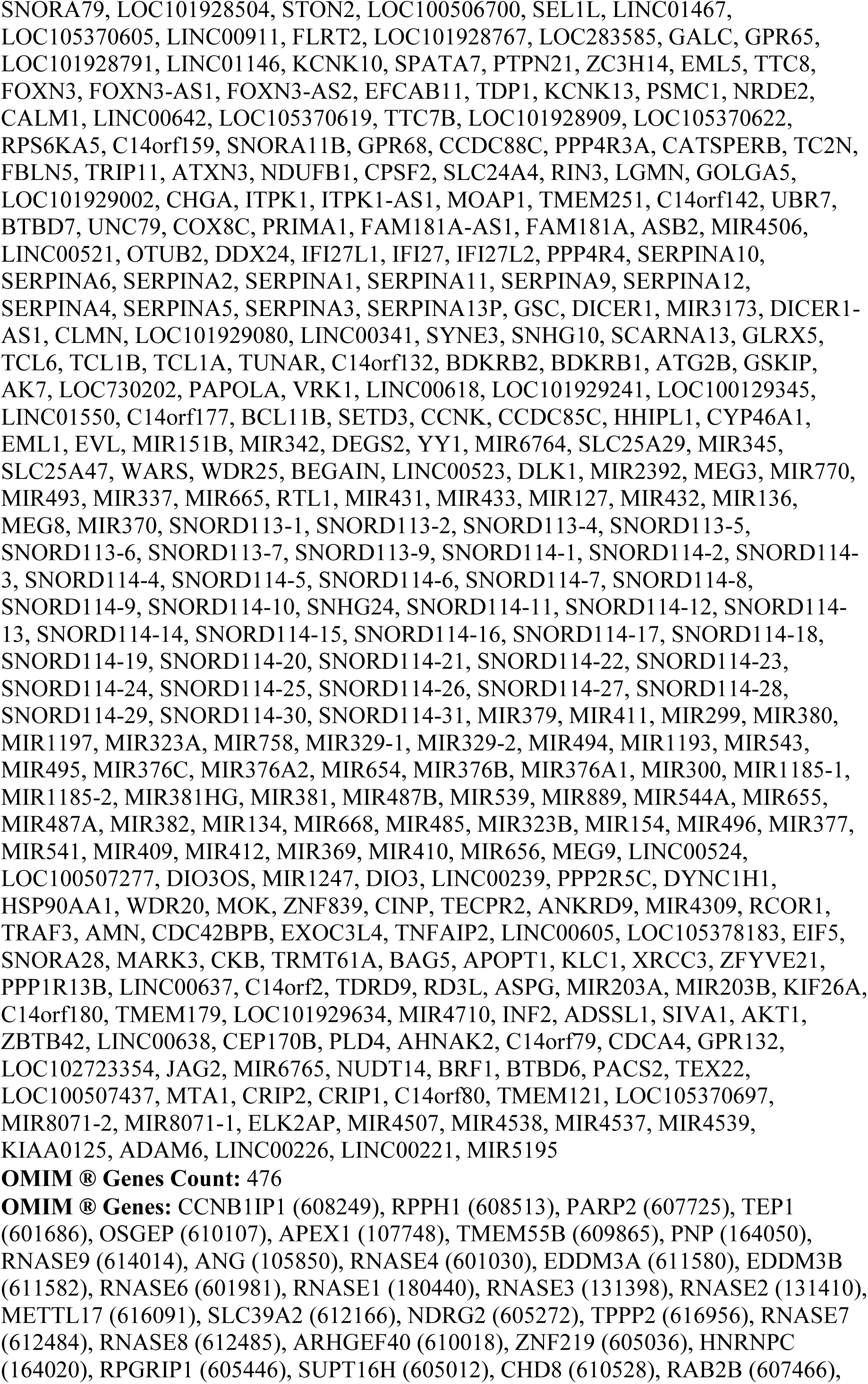
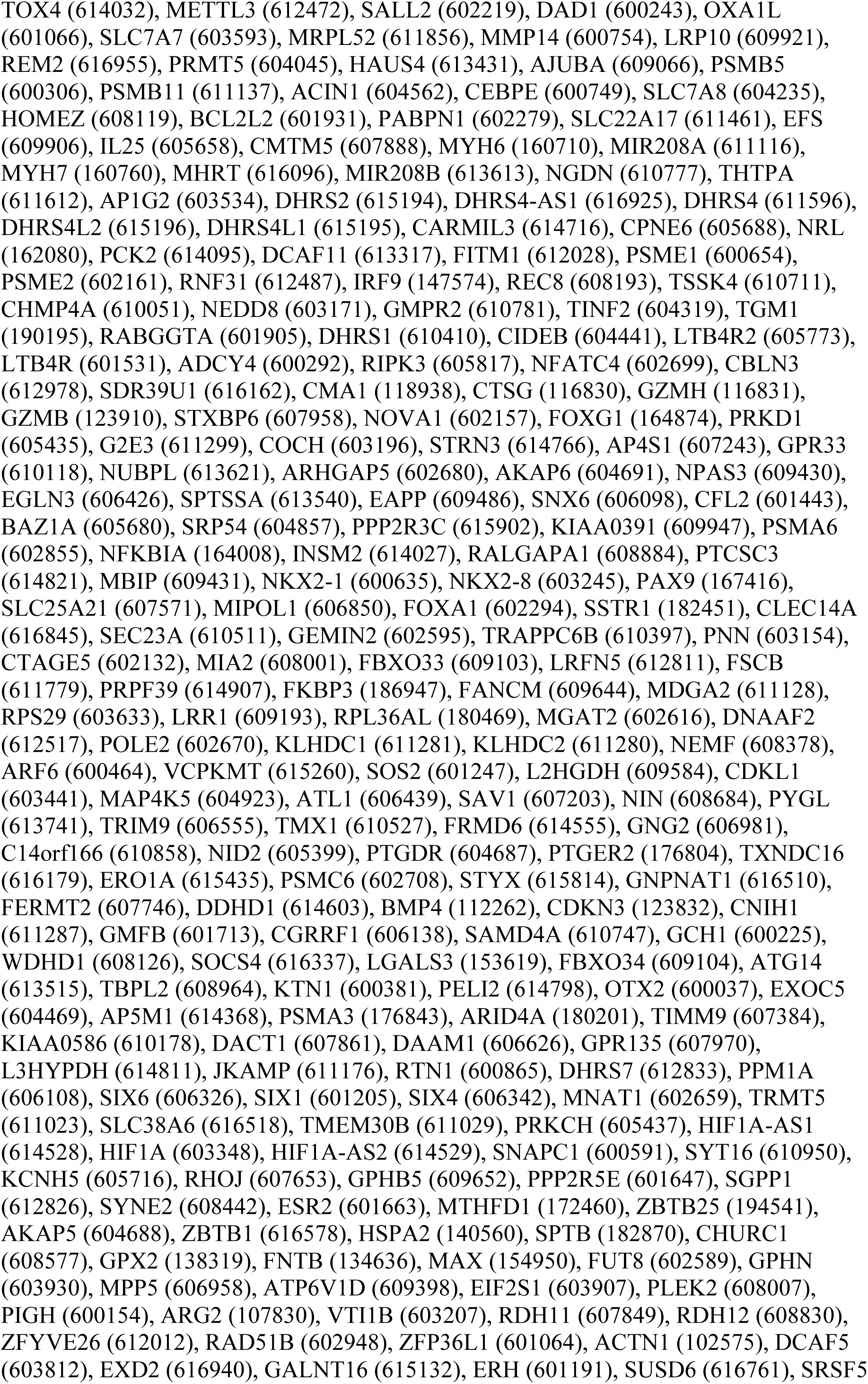
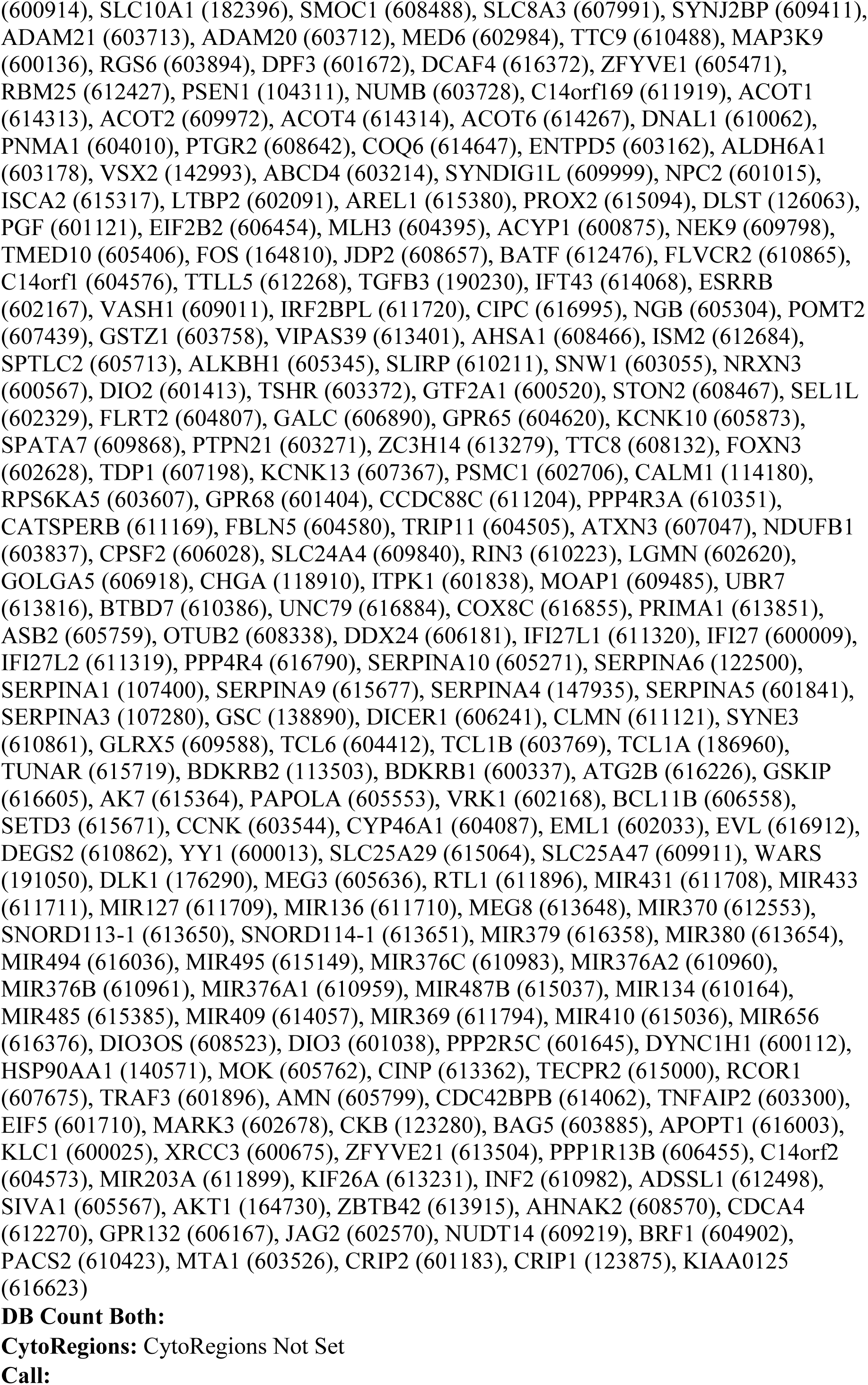
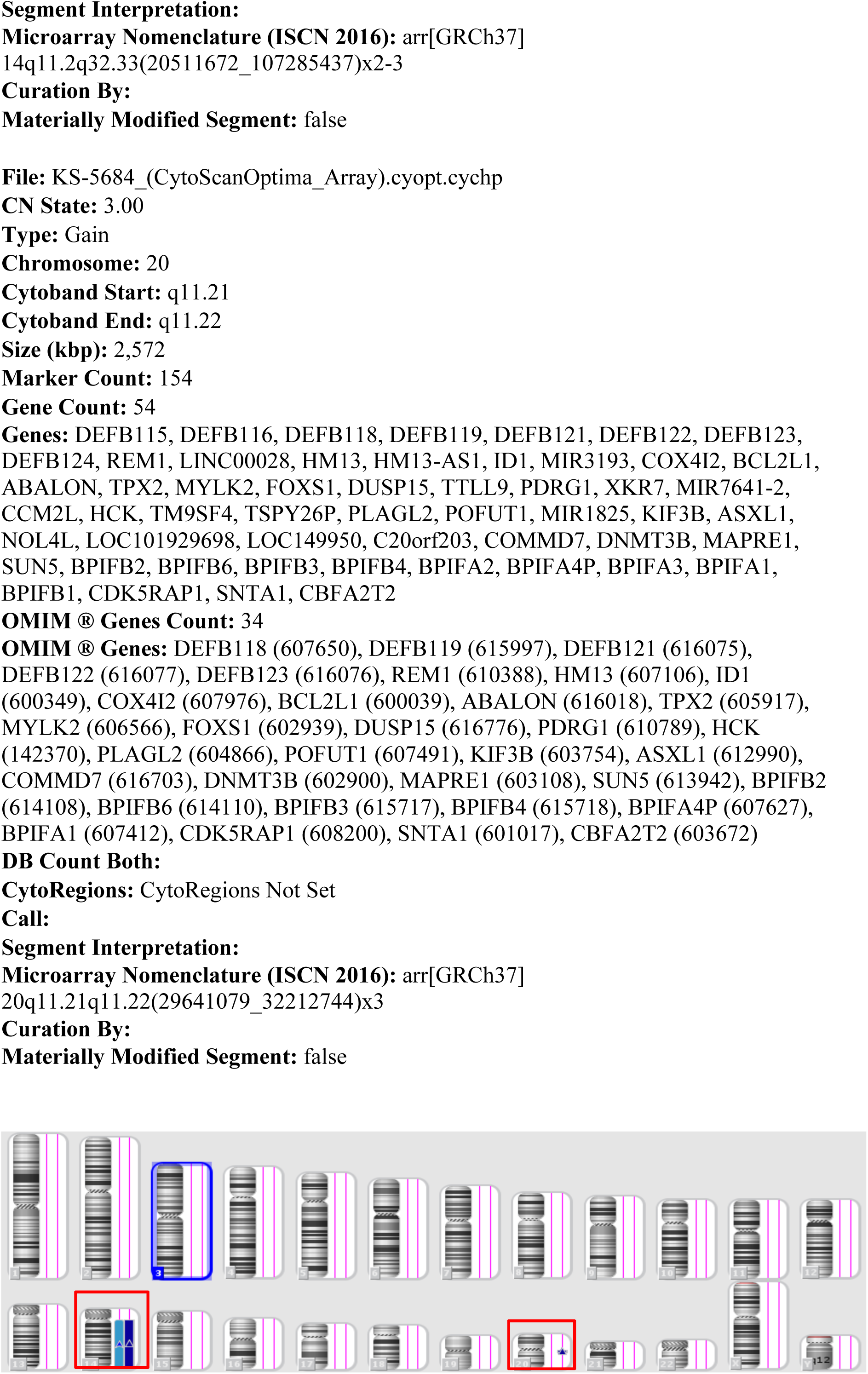
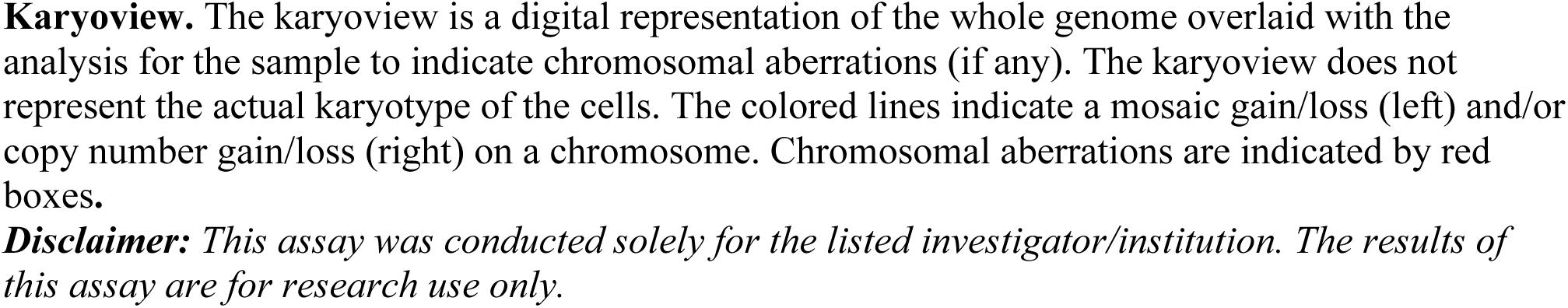
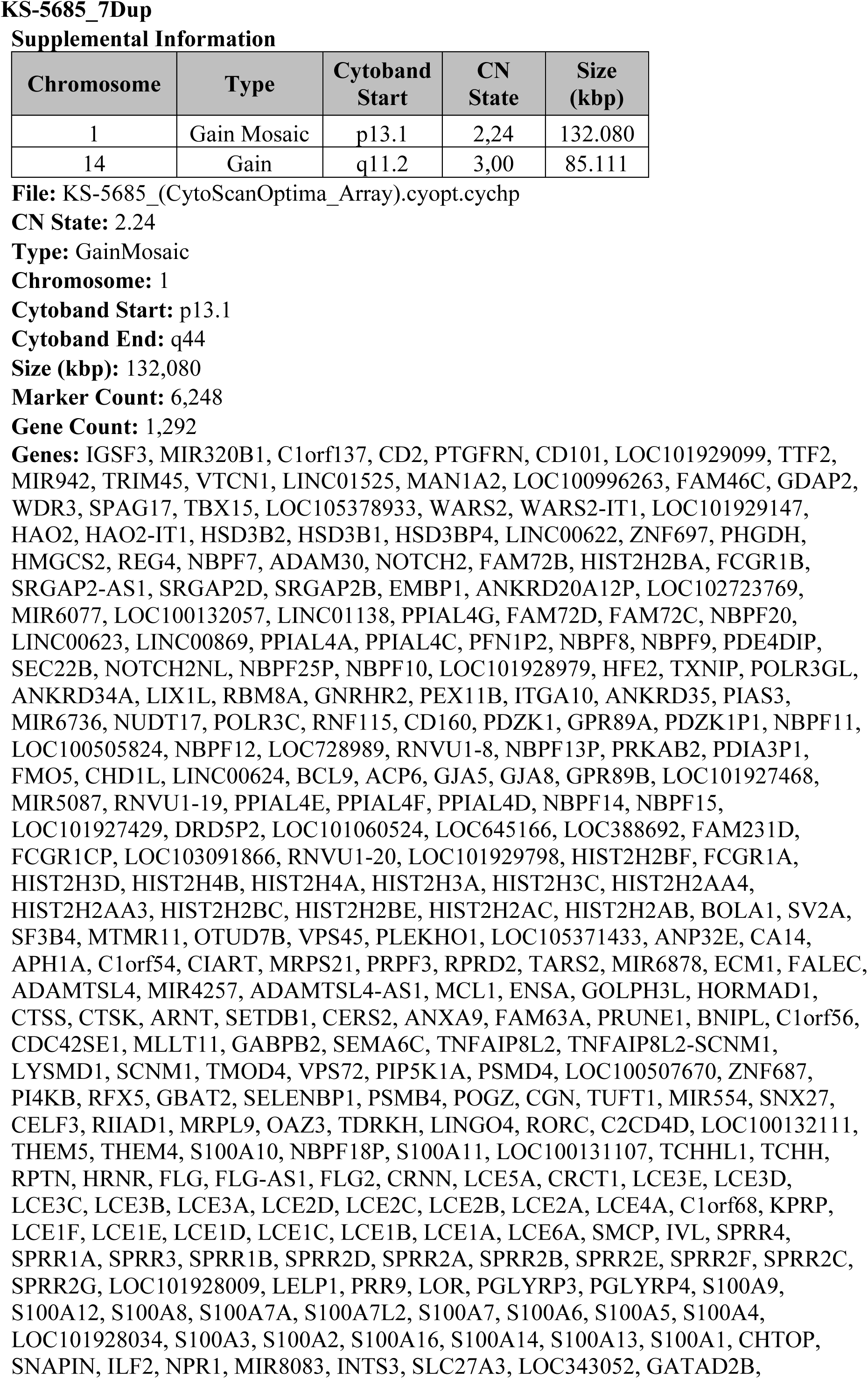
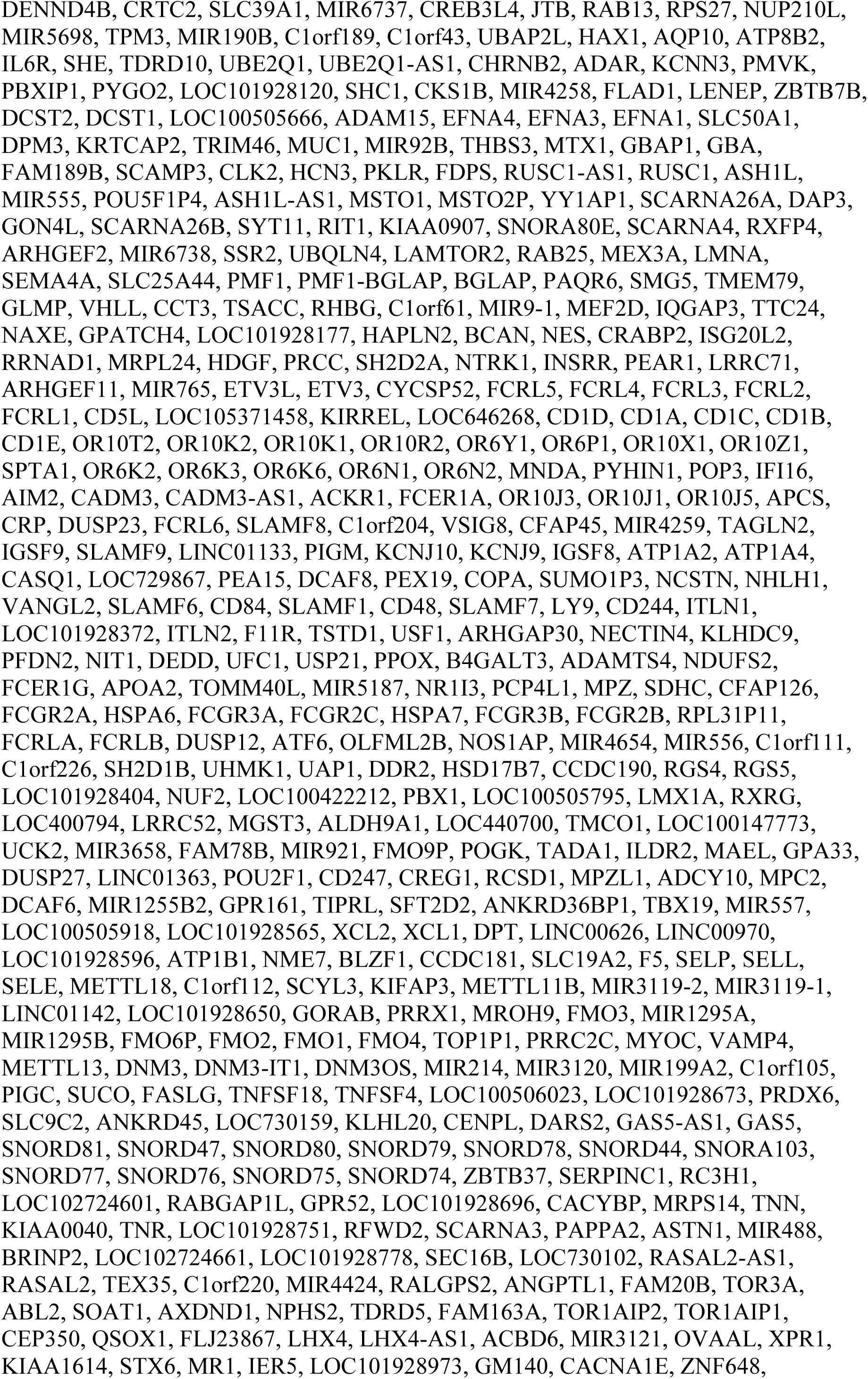
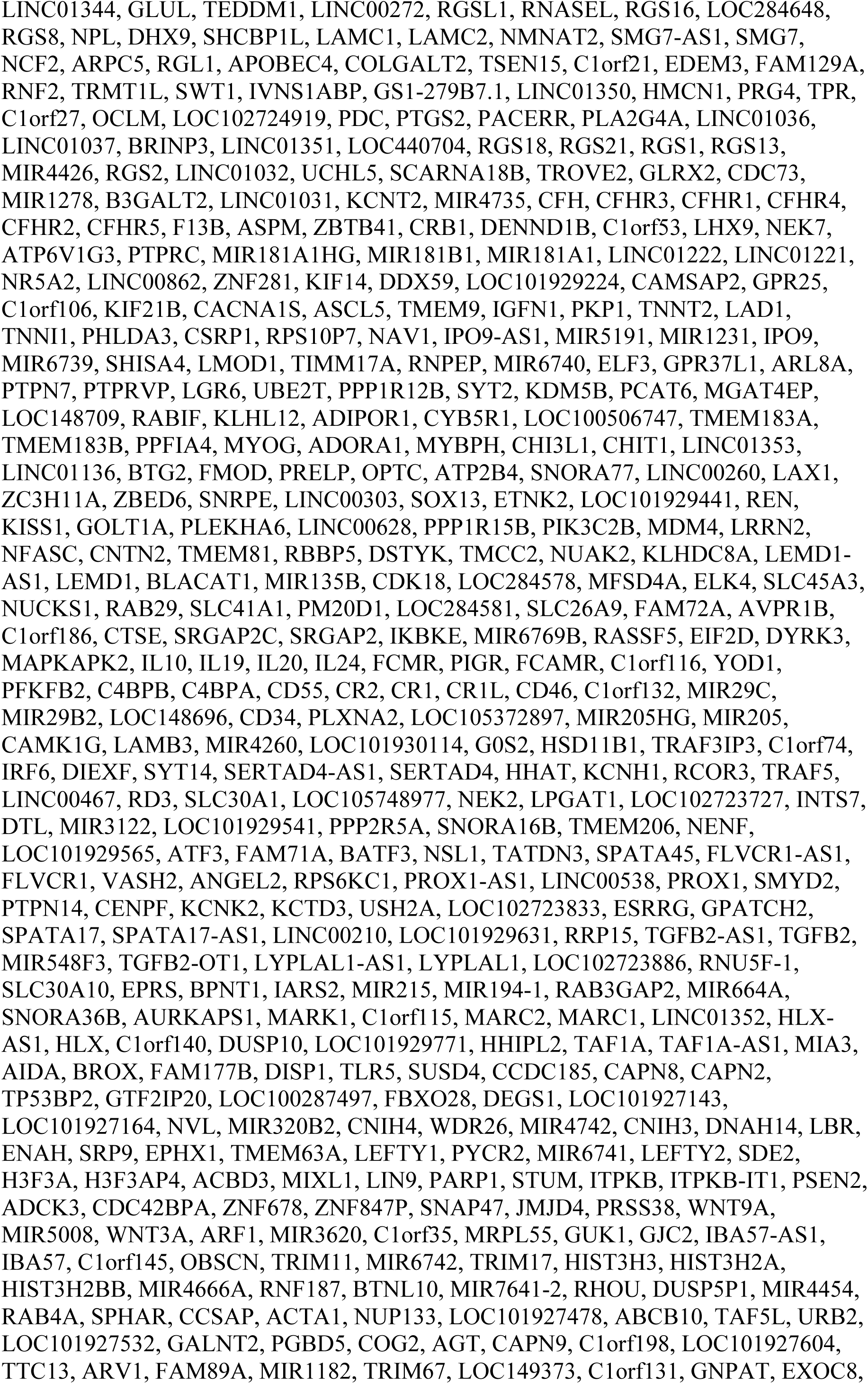
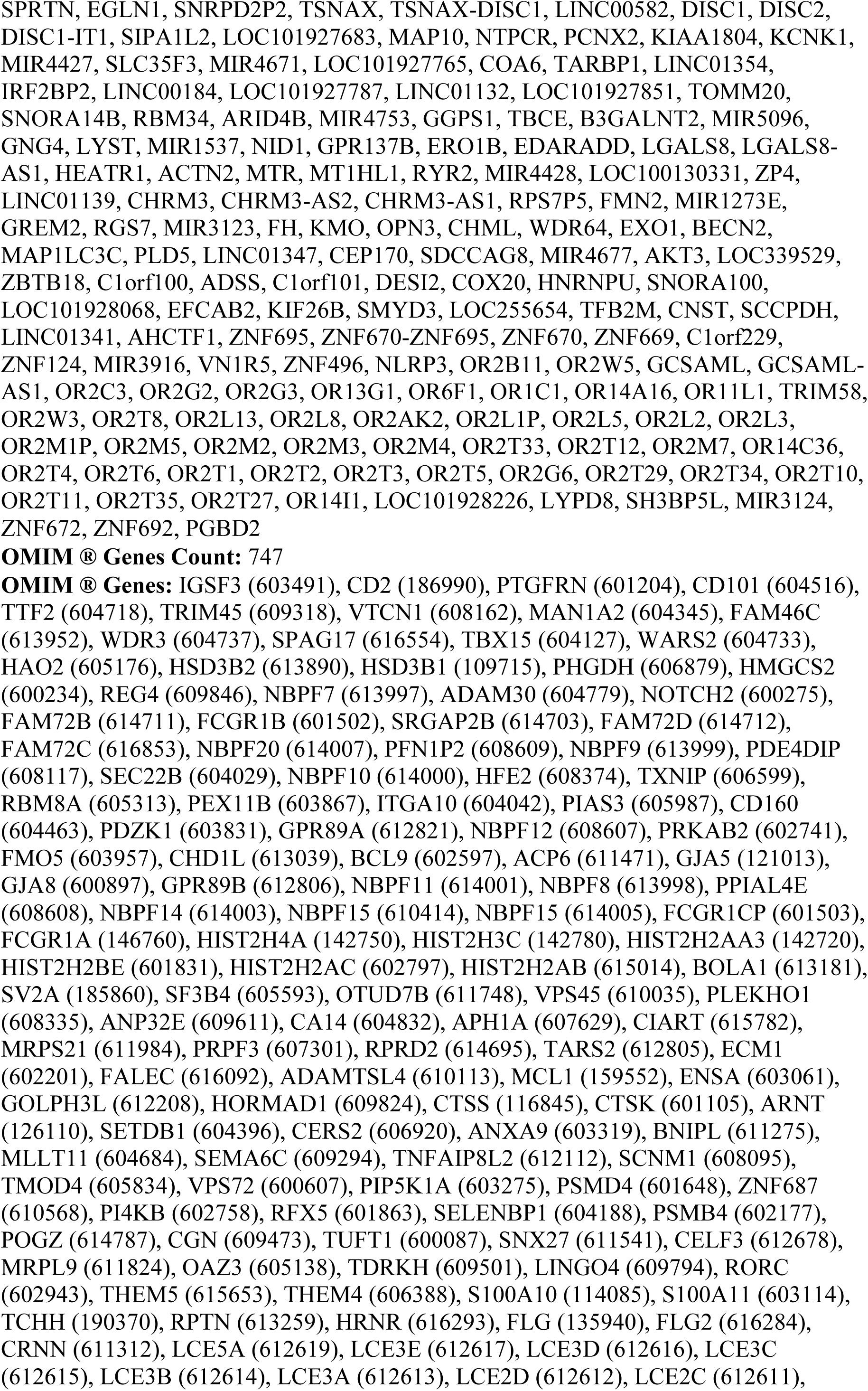
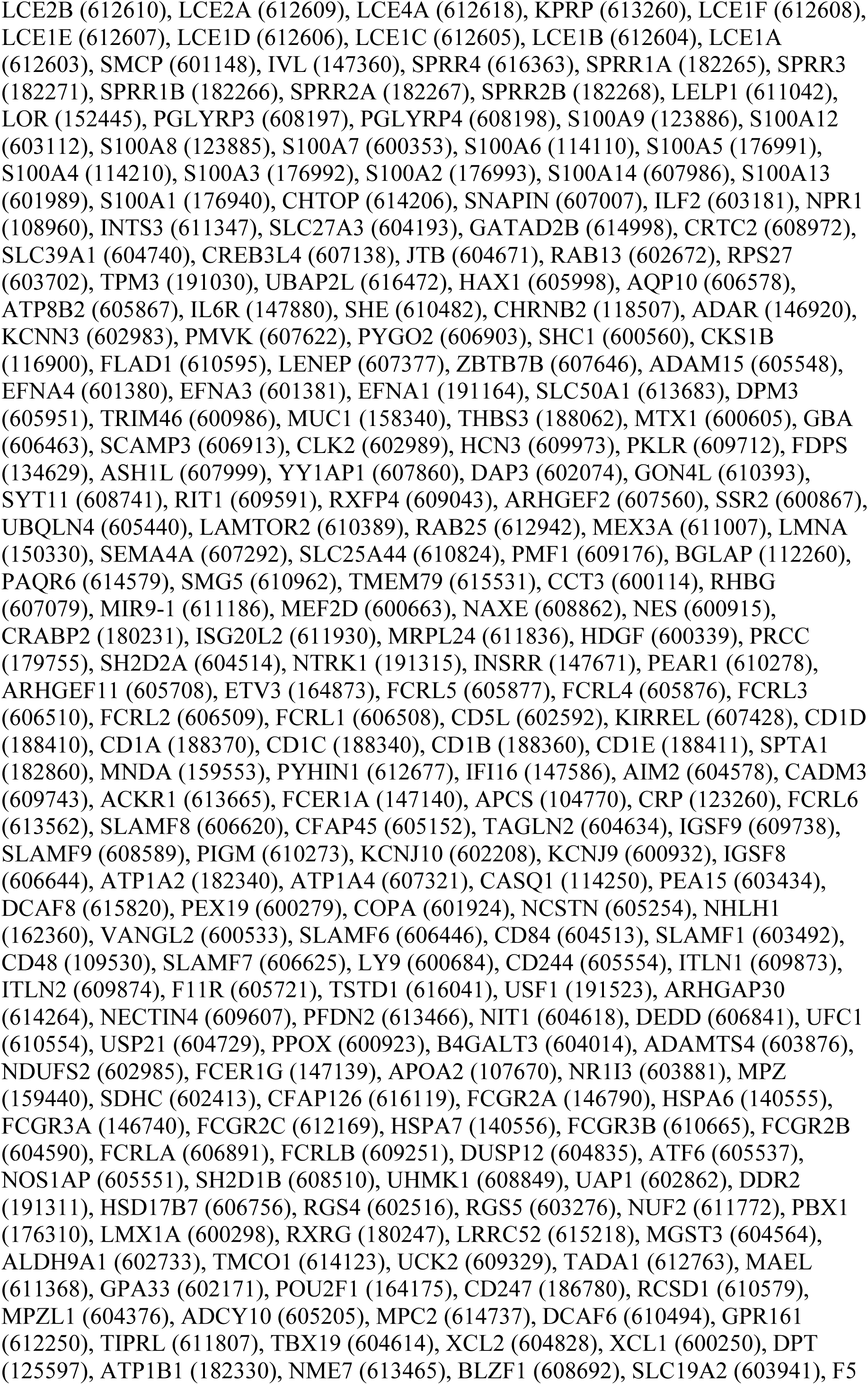
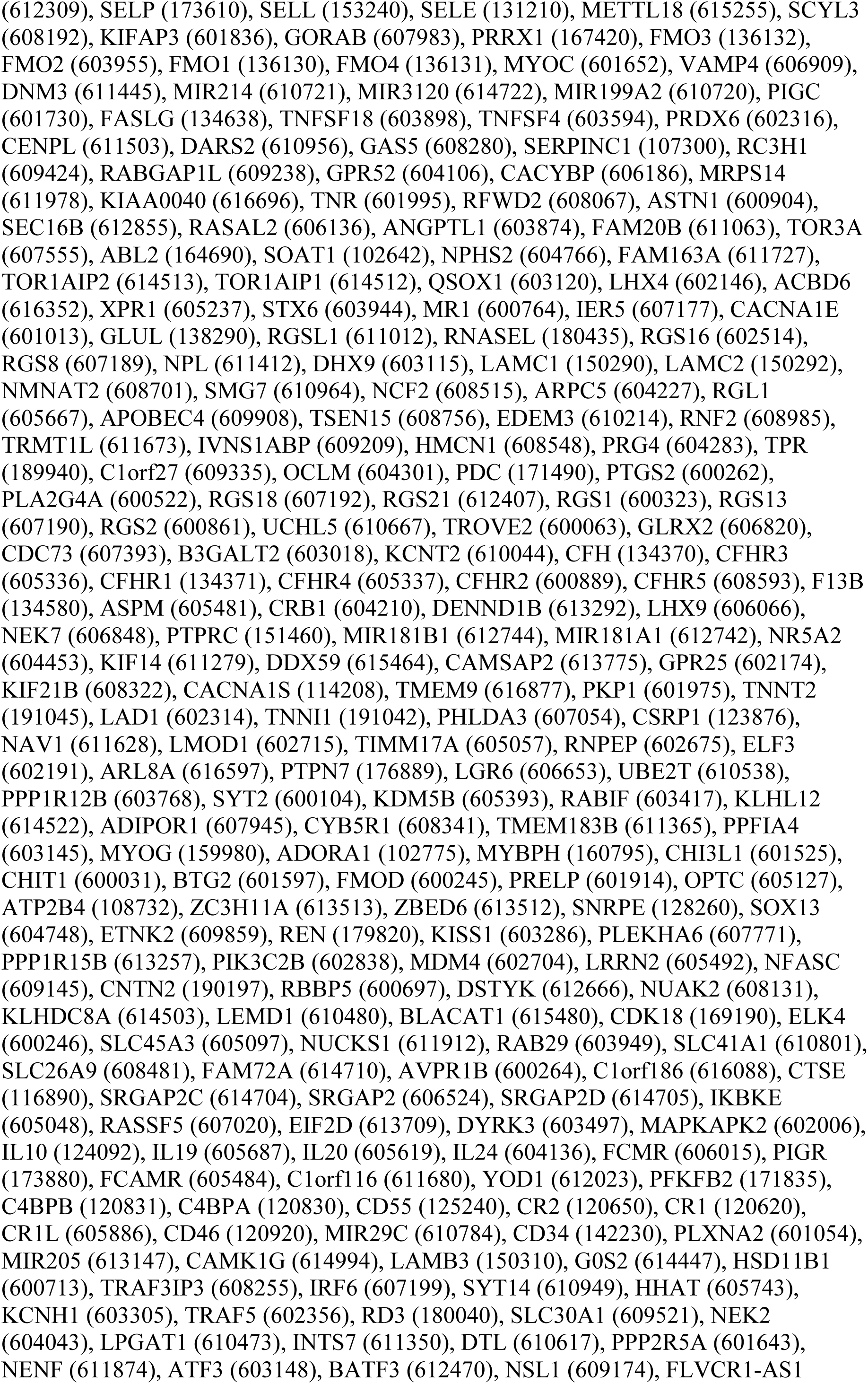
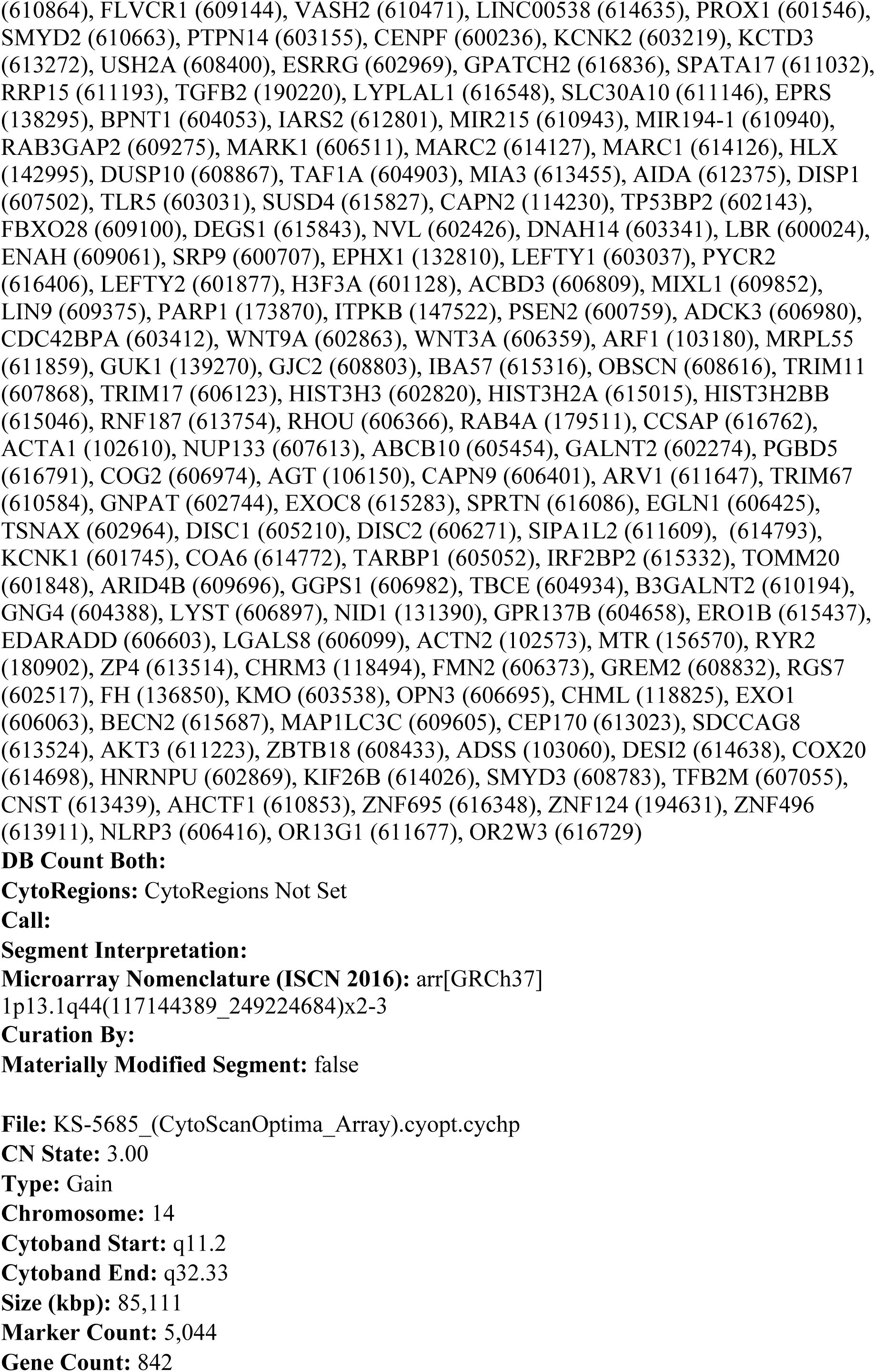
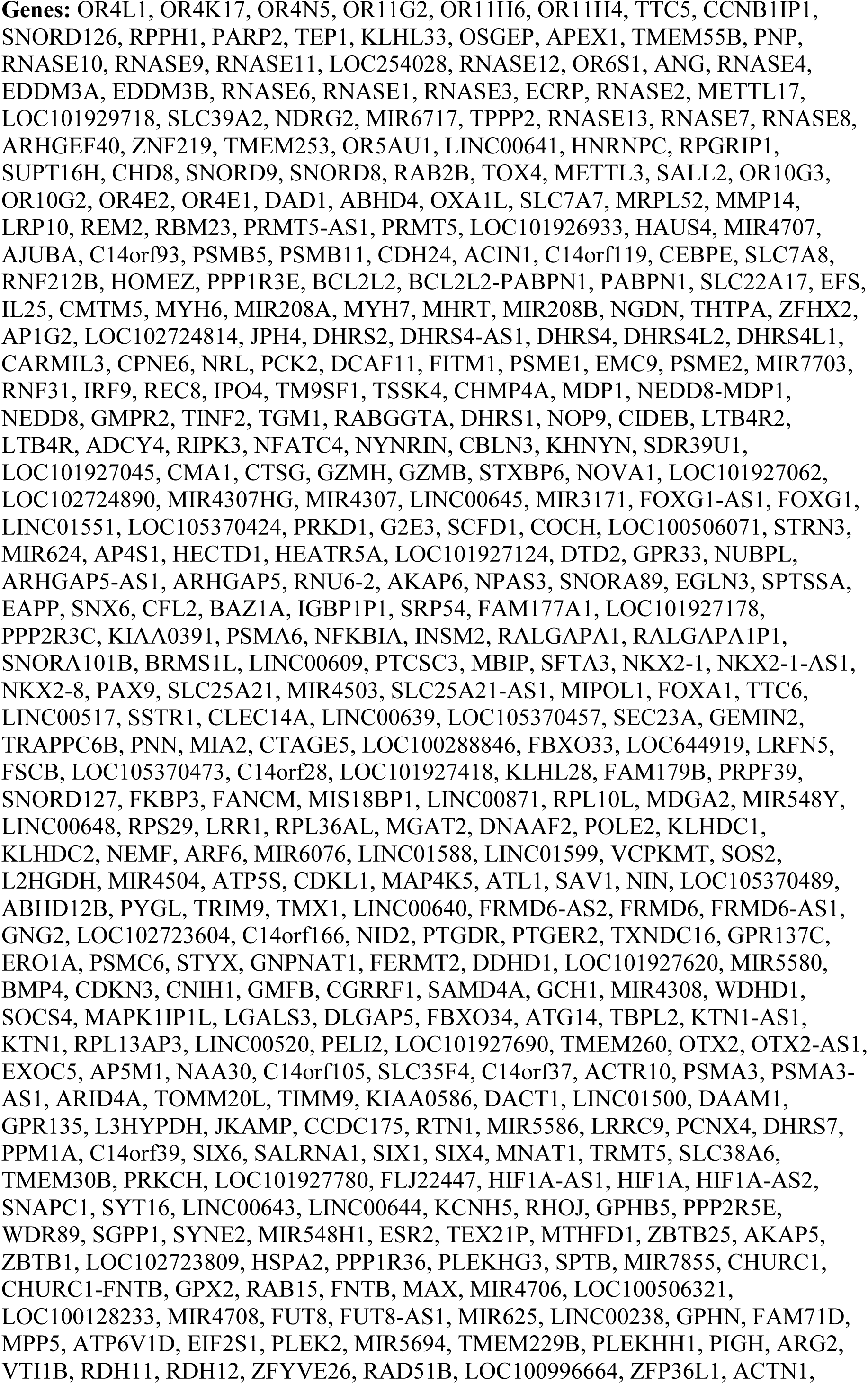
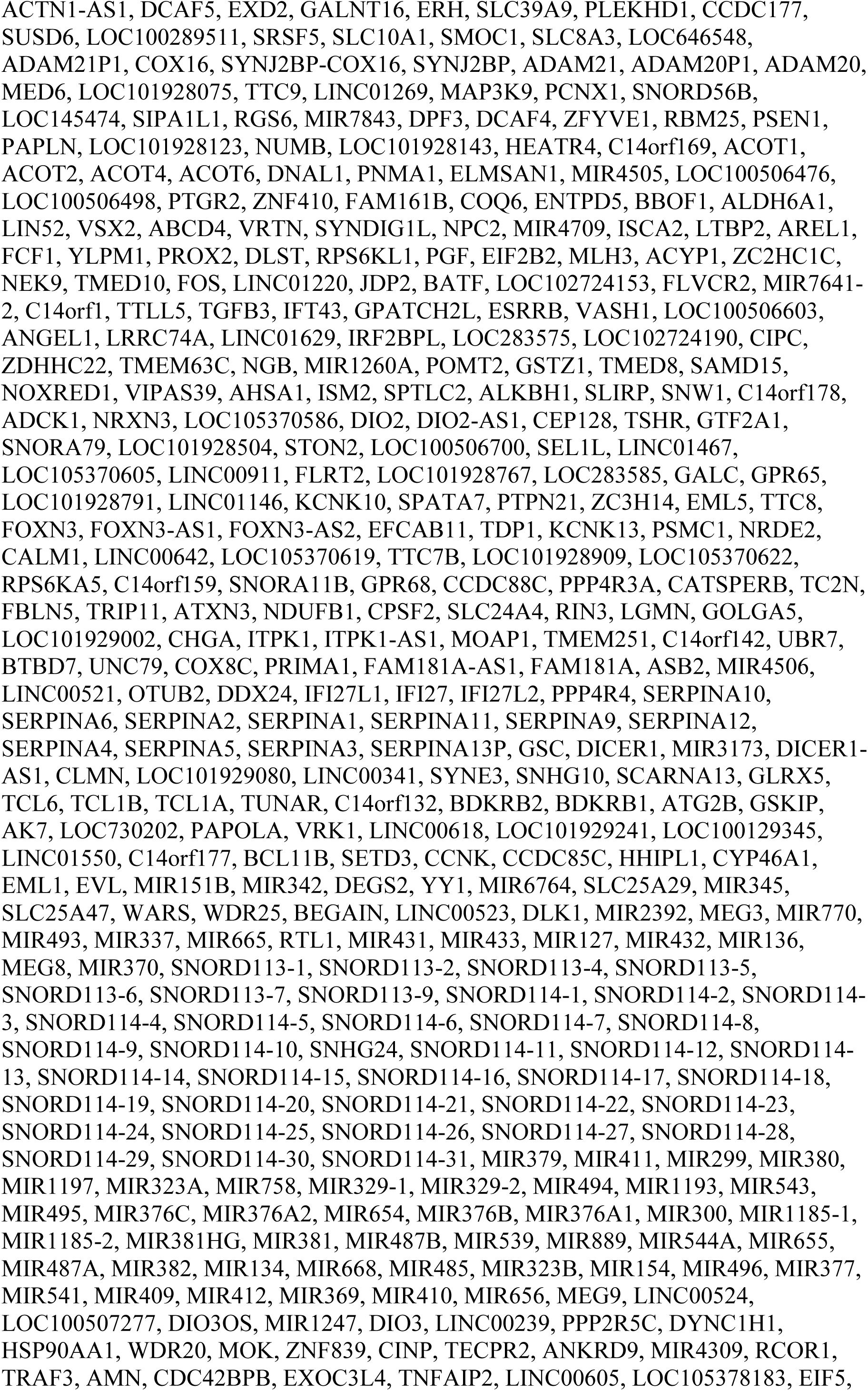
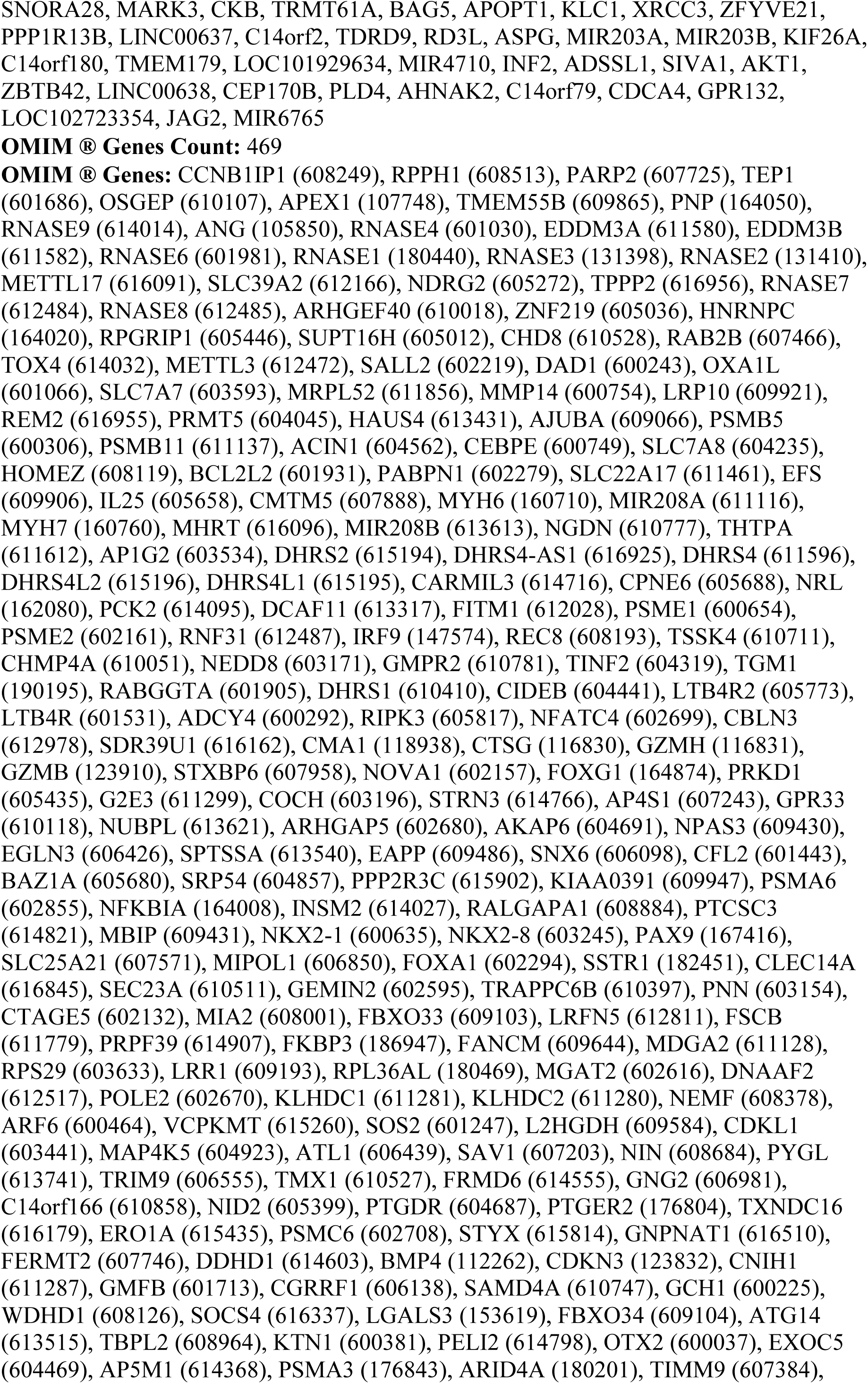
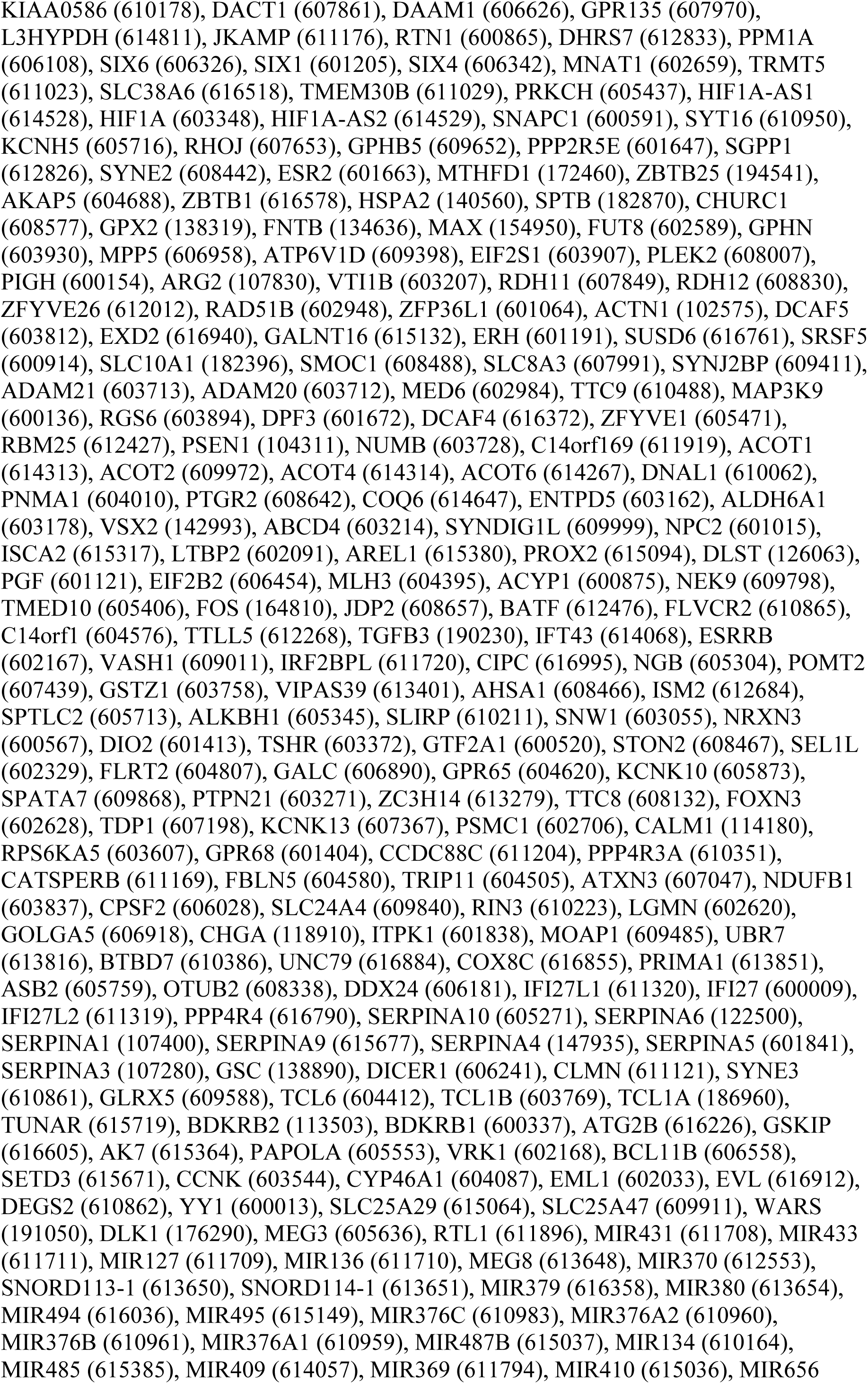
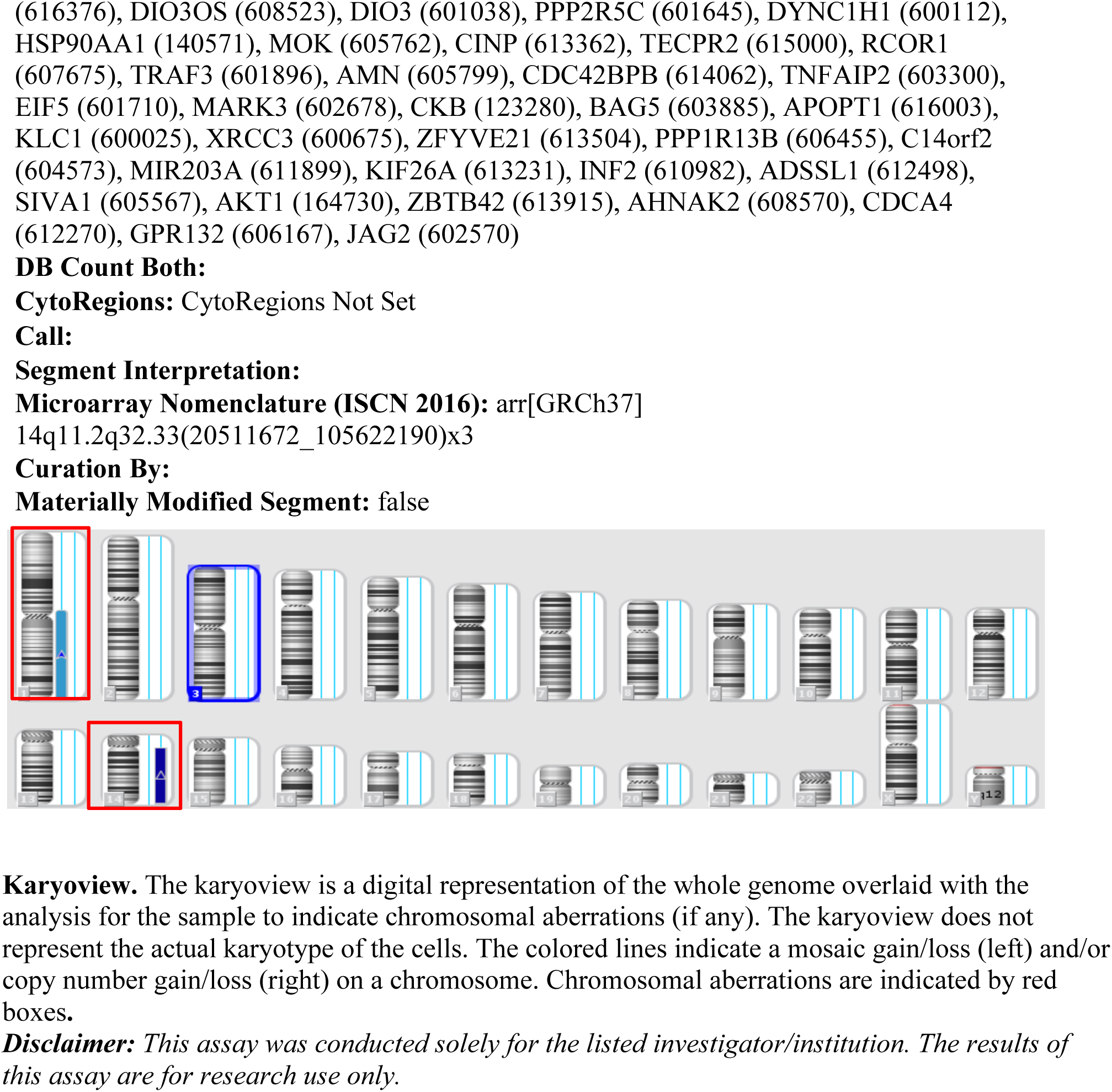

**Supplementary Table S2 related to Fig. 3C.**
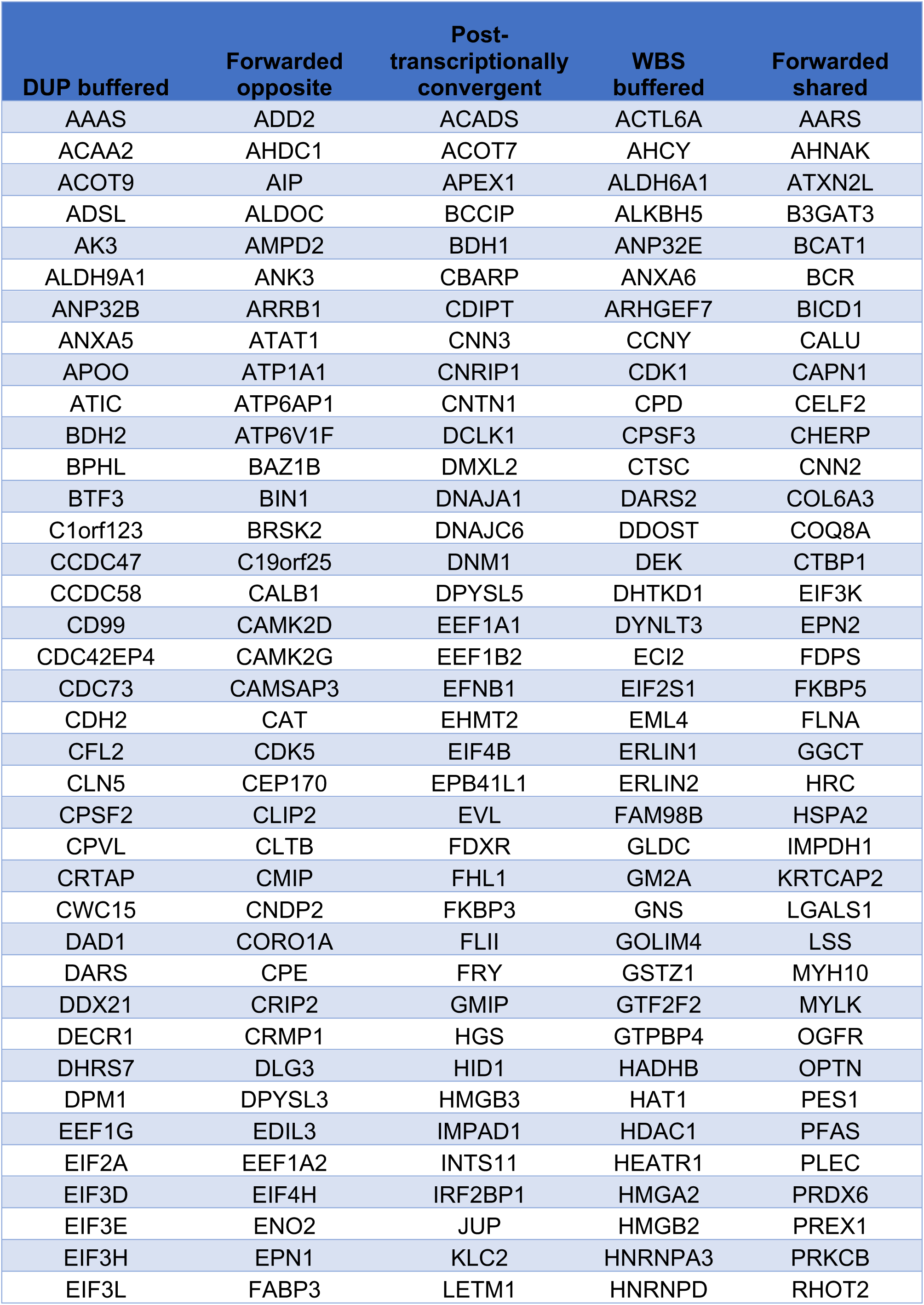
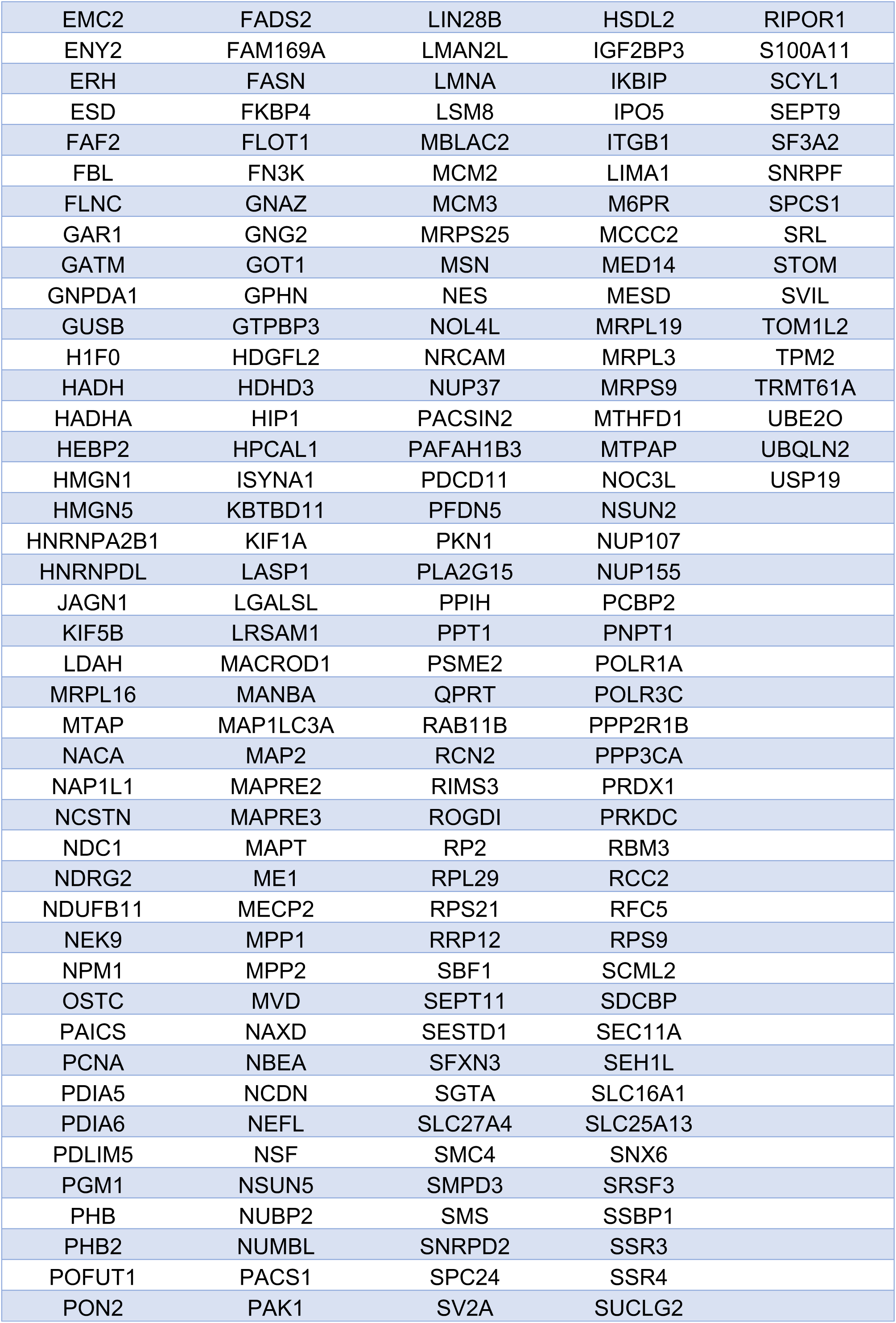
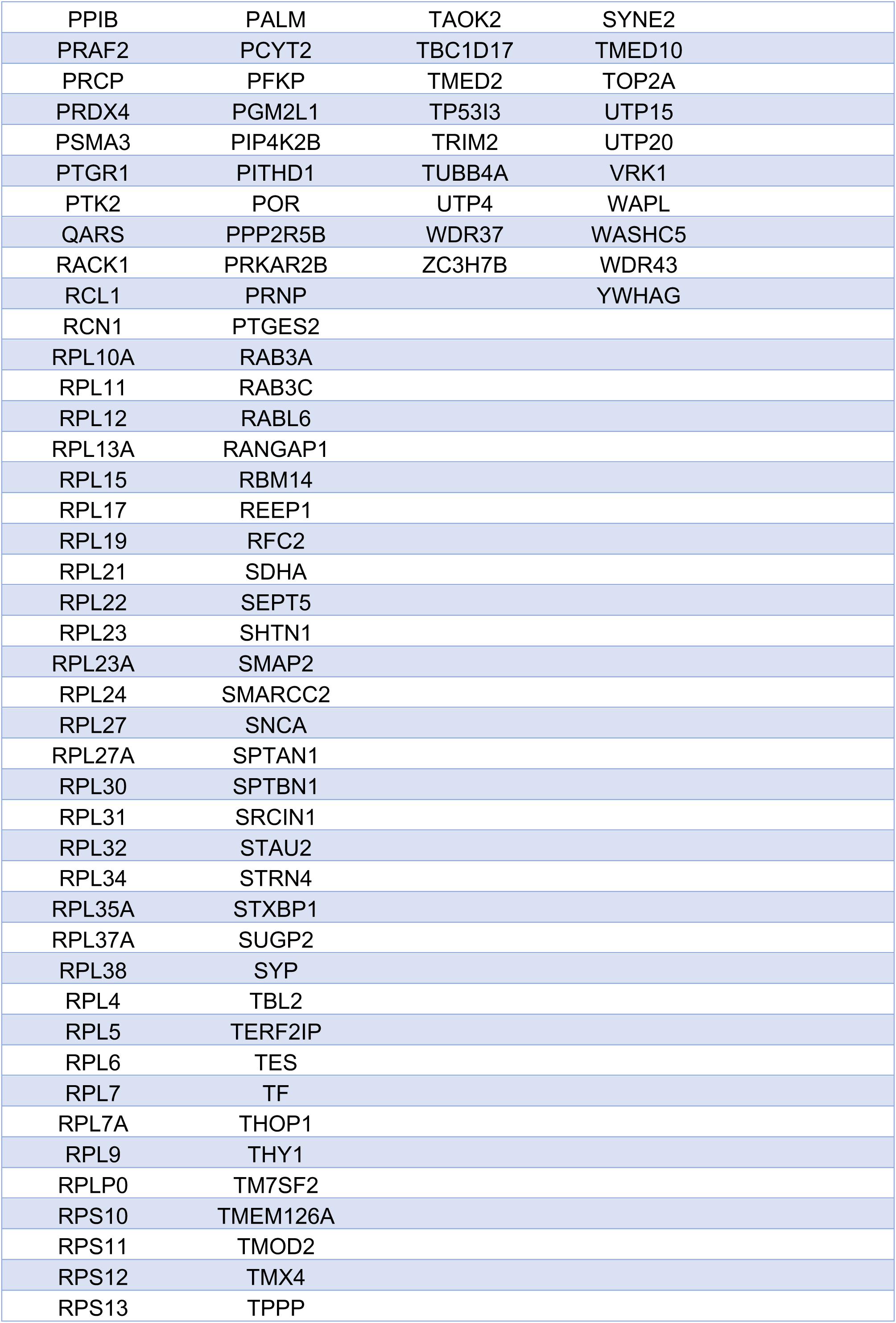
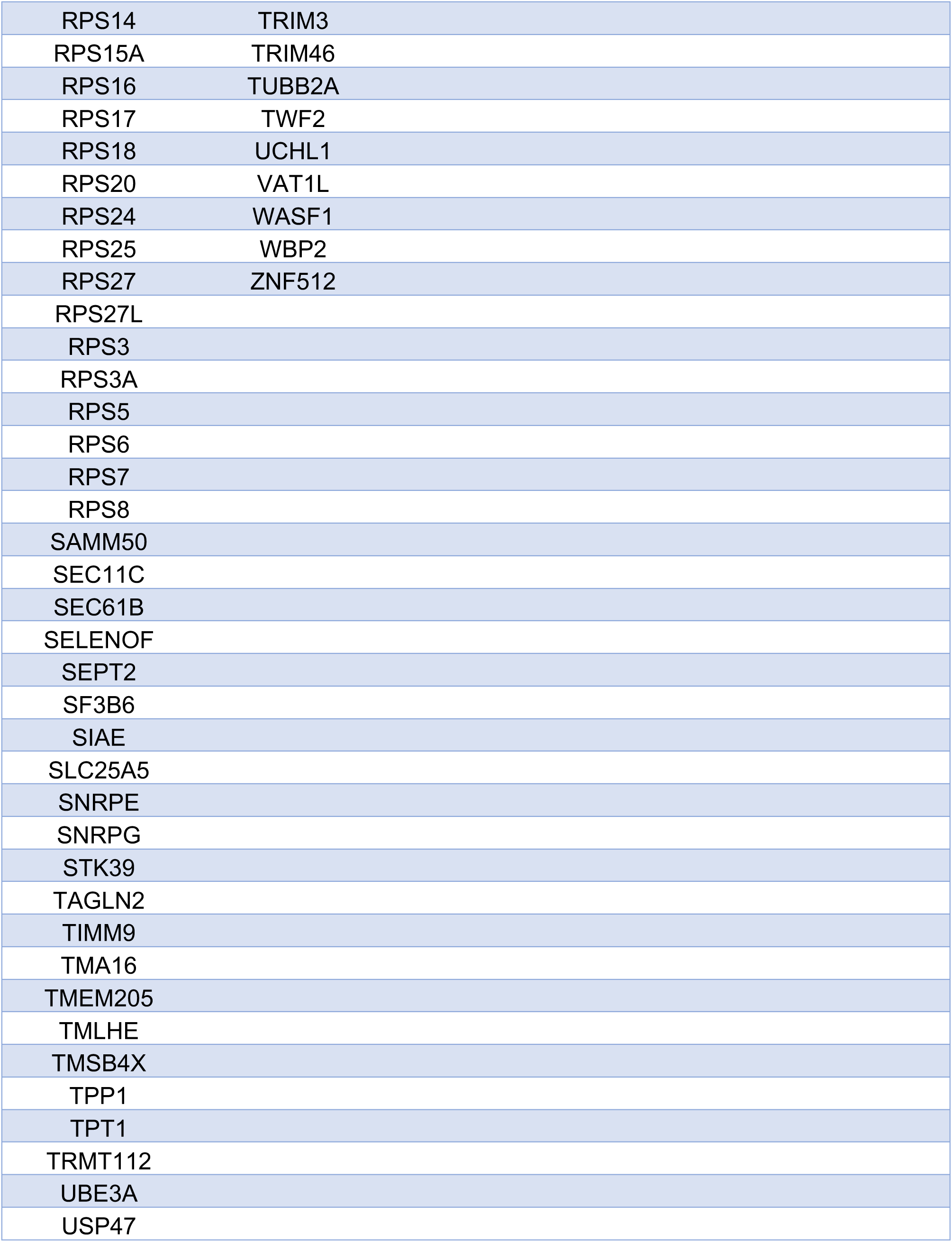

| **DUP buffered** | **Forwarded opposite** | **Post-transcriptionally convergent** | **WBS buffered** | **Forwarded shared** |
| --- | --- | --- | --- | --- |
| AAAS | ADD2 | ACADS | ACTL6A | AARS |
| ACAA2 | AHDC1 | ACOT7 | AHCY | AHNAK |
| ACOT9 | AIP | APEX1 | ALDH6A1 | ATXN2L |
| ADSL | ALDOC | BCCIP | ALKBH5 | B3GAT3 |
| AK3 | AMPD2 | BDH1 | ANP32E | BCAT1 |
| ALDH9A1 | ANK3 | CBARP | ANXA6 | BCR |
| ANP32B | ARRB1 | CDIPT | ARHGEF7 | BICD1 |
| ANXA5 | ATAT1 | CNN3 | CCNY | CALU |
| APOO | ATP1A1 | CNRIP1 | CDK1 | CAPN1 |
| ATIC | ATP6AP1 | CNTN1 | CPD | CELF2 |
| BDH2 | ATP6V1F | DCLK1 | CPSF3 | CHERP |
| BPHL | BAZ1B | DMXL2 | CTSC | CNN2 |
| BTF3 | BIN1 | DNAJA1 | DARS2 | COL6A3 |
| C1orf123 | BRSK2 | DNAJC6 | DDOST | COQ8A |
| CCDC47 | C19orf25 | DNM1 | DEK | CTBP1 |
| CCDC58 | CALB1 | DPYSL5 | DHTKD1 | EIF3K |
| CD99 | CAMK2D | EEF1A1 | DYNLT3 | EPN2 |
| CDC42EP4 | CAMK2G | EEF1B2 | ECI2 | FDPS |
| CDC73 | CAMSAP3 | EFNB1 | EIF2S1 | FKBP5 |
| CDH2 | CAT | EHMT2 | EML4 | FLNA |
| CFL2 | CDK5 | EIF4B | ERLIN1 | GGCT |
| CLN5 | CEP170 | EPB41L1 | ERLIN2 | HRC |
| CPSF2 | CLIP2 | EVL | FAM98B | HSPA2 |
| CPVL | CLTB | FDXR | GLDC | IMPDH1 |
| CRTAP | CMIP | FHL1 | GM2A | KRTCAP2 |
| CWC15 | CNDP2 | FKBP3 | GNS | LGALS1 |
| DAD1 | CORO1A | FLII | GOLIM4 | LSS |
| DARS | CPE | FRY | GSTZ1 | MYH10 |
| DDX21 | CRIP2 | GMIP | GTF2F2 | MYLK |
| DECR1 | CRMP1 | HGS | GTPBP4 | OGFR |
| DHRS7 | DLG3 | HID1 | HADHB | OPTN |
| DPM1 | DPYSL3 | HMGB3 | HAT1 | PES1 |
| EEF1G | EDIL3 | IMPAD1 | HDAC1 | PFAS |
| EIF2A | EEF1A2 | INTS11 | HEATR1 | PLEC |
| EIF3D | EIF4H | IRF2BP1 | HMGA2 | PRDX6 |
| EIF3E | ENO2 | JUP | HMGB2 | PREX1 |
| EIF3H | EPN1 | KLC2 | HNRNPA3 | PRKCB |
| EIF3L | FABP3 | LETM1 | HNRNPD | RHOT2 |
| EMC2 | FADS2 | LIN28B | HSDL2 | RIPOR1 |
| ENY2 | FAM169A | LMAN2L | IGF2BP3 | S100A11 |
| ERH | FASN | LMNA | IKBIP | SCYL1 |
| ESD | FKBP4 | LSM8 | IPO5 | SEPT9 |
| FAF2 | FLOT1 | MBLAC2 | ITGB1 | SF3A2 |
| FBL | FN3K | MCM2 | LIMA1 | SNRPF |
| FLNC | GNAZ | MCM3 | M6PR | SPCS1 |
| GAR1 | GNG2 | MRPS25 | MCCC2 | SRL |
| GATM | GOT1 | MSN | MED14 | STOM |
| GNPDA1 | GPHN | NES | MESD | SVIL |
| GUSB | GTPBP3 | NOL4L | MRPL19 | TOM1L2 |
| H1F0 | HDGFL2 | NRCAM | MRPL3 | TPM2 |
| HADH | HDHD3 | NUP37 | MRPS9 | TRMT61A |
| HADHA | HIP1 | PACSIN2 | MTHFD1 | UBE2O |
| HEBP2 | HPCAL1 | PAFAH1B3 | MTPAP | UBQLN2 |
| HMGN1 | ISYNA1 | PDCD11 | NOC3L | USP19 |
| HMGN5 | KBTBD11 | PFDN5 | NSUN2 |  |
| HNRNPA2B1 | KIF1A | PKN1 | NUP107 |  |
| HNRNPDL | LASP1 | PLA2G15 | NUP155 |  |
| JAGN1 | LGALSL | PPIH | PCBP2 |  |
| KIF5B | LRSAM1 | PPT1 | PNPT1 |  |
| LDAH | MACROD1 | PSME2 | POLR1A |  |
| MRPL16 | MANBA | QPRT | POLR3C |  |
| MTAP | MAP1LC3A | RAB11B | PPP2R1B |  |
| NACA | MAP2 | RCN2 | PPP3CA |  |
| NAP1L1 | MAPRE2 | RIMS3 | PRDX1 |  |
| NCSTN | MAPRE3 | ROGDI | PRKDC |  |
| NDC1 | MAPT | RP2 | RBM3 |  |
| NDRG2 | ME1 | RPL29 | RCC2 |  |
| NDUFB11 | MECP2 | RPS21 | RFC5 |  |
| NEK9 | MPP1 | RRP12 | RPS9 |  |
| NPM1 | MPP2 | SBF1 | SCML2 |  |
| OSTC | MVD | SEPT11 | SDCBP |  |
| PAICS | NAXD | SESTD1 | SEC11A |  |
| PCNA | NBEA | SFXN3 | SEH1L |  |
| PDIA5 | NCDN | SGTA | SLC16A1 |  |
| PDIA6 | NEFL | SLC27A4 | SLC25A13 |  |
| PDLIM5 | NSF | SMC4 | SNX6 |  |
| PGM1 | NSUN5 | SMPD3 | SRSF3 |  |
| PHB | NUBP2 | SMS | SSBP1 |  |
| PHB2 | NUMBL | SNRPD2 | SSR3 |  |
| POFUT1 | PACS1 | SPC24 | SSR4 |  |
| PON2 | PAK1 | SV2A | SUCLG2 |  |

|  |  |  |  |
| --- | --- | --- | --- |
| PPIB | PALM | TAOK2 | SYNE2 |
| PRAF2 | PCYT2 | TBC1D17 | TMED10 |
| PRCP | PFKP | TMED2 | TOP2A |
| PRDX4 | PGM2L1 | TP53I3 | UTP15 |
| PSMA3 | PIP4K2B | TRIM2 | UTP20 |
| PTGR1 | PITHD1 | TUBB4A | VRK1 |
| PTK2 | POR | UTP4 | WAPL |
| QARS | PPP2R5B | WDR37 | WASHC5 |
| RACK1 | PRKAR2B | ZC3H7B | WDR43 |
| RCL1 | PRNP |  | YWHAG |
| RCN1 | PTGES2 |  |  |
| RPL10A | RAB3A |  |  |
| RPL11 | RAB3C |  |  |
| RPL12 | RABL6 |  |  |
| RPL13A | RANGAP1 |  |  |
| RPL15 | RBM14 |  |  |
| RPL17 | REEP1 |  |  |
| RPL19 | RFC2 |  |  |
| RPL21 | SDHA |  |  |
| RPL22 | SEPT5 |  |  |
| RPL23 | SHTN1 |  |  |
| RPL23A | SMAP2 |  |  |
| RPL24 | SMARCC2 |  |  |
| RPL27 | SNCA |  |  |
| RPL27A | SPTAN1 |  |  |
| RPL30 | SPTBN1 |  |  |
| RPL31 | SRCIN1 |  |  |
| RPL32 | STAU2 |  |  |
| RPL34 | STRN4 |  |  |
| RPL35A | STXBP1 |  |  |
| RPL37A | SUGP2 |  |  |
| RPL38 | SYP |  |  |
| RPL4 | TBL2 |  |  |
| RPL5 | TERF2IP |  |  |
| RPL6 | TES |  |  |
| RPL7 | TF |  |  |
| RPL7A | THOP1 |  |  |
| RPL9 | THY1 |  |  |
| RPLP0 | TM7SF2 |  |  |
| RPS10 | TMEM126A |  |  |
| RPS11 | TMOD2 |  |  |
| RPS12 | TMX4 |  |  |
| RPS13 | TPPP |  |  |

|  |  |
| --- | --- |
| RPS14 | TRIM3 |
| RPS15A | TRIM46 |
| RPS16 | TUBB2A |
| RPS17 | TWF2 |
| RPS18 | UCHL1 |
| RPS20 | VAT1L |
| RPS24 | WASF1 |
| RPS25 | WBP2 |
| RPS27 | ZNF512 |
| RPS27L |  |
| RPS3 |  |
| RPS3A |  |
| RPS5 |  |
| RPS6 |  |
| RPS7 |  |
| RPS8 |  |
| SAMM50 |  |
| SEC11C |  |
| SEC61B |  |
| SELENOF |  |
| SEPT2 |  |
| SF3B6 |  |
| SIAE |  |
| SLC25A5 |  |
| SNRPE |  |
| SNRPG |  |
| STK39 |  |
| TAGLN2 |  |
| TIMM9 |  |
| TMA16 |  |
| TMEM205 |  |
| TMLHE |  |
| TMSB4X |  |
| TPP1 |  |
| TPT1 |  |
| TRMT112 |  |
| UBE3A |  |
| USP47 |  |

